# Dual Inhibitors Targeting G9a and GSK-3β: Translational Perspectives on Alzheimer’s Disease Treatment

**DOI:** 10.1101/2025.10.24.684310

**Authors:** Aina Bellver-Sanchis, Ainoa Sanchez-Arfelis, Bhanwar Singh Choudhary, Alba Irisarri, Marta Ribalta- Vilella, Sukanya Sukanya, Deb Ranjan Banerjee, José A. Morales-García, Carolin Wüst, Aida Castellanos, David Soto, Núria Casals, Rut Fadó, Ana Guerrero, Gemma Navarro, Juan A. Fafián-Labora, Anna Perez-Bosque, Lluïsa Miró, Muhammad Ali, Manahil Saqib, Zaeem Ali, Sajid Rashid, Belén Pérez, Antón Leandro Martínez Rodríguez, Jose Brea, Ruchi Malik, Santiago Vázquez, Mercè Pallas, Rafael Franco, David Valle-García, Carmen Escolano, Christian Griñán-Ferré

**Affiliations:** Department of Pharmacology, Toxicology and Therapeutic Chemistry, Institut de Neurociències-Universitat de Barcelona, Barcelona, Spain; Laboratory of Medicinal Chemistry. Department of Pharmacology, Toxicology and Medicinal Chemistry, Faculty of Pharmacy and Food Sciences, University of Barcelona, Av. Joan XXIII, 27-31, 08028 Barcelona, Spain; Department of Pharmacy, Central University of Rajasthan, Bandarsindari, Ajmer, Rajasthan, 305817, India; Department of Pharmacognosy, SVKM’s Dr. Bhanuben Nanavati College of Pharmacy, Vile Parle (West), Mumbai, Maharashtra, India, 400056; Department of Chemistry, National Institute of Technology Durgapur, MG Avenue, Durgapur 713209, India; Department of Cell Biology. Faculty of Medicine, Computense University of Madrid. (UCM), Madrid 28040, Spain; Neurophysiology Laboratory, Department of Biomedicine, Faculty of Medicine and Health Sciences, Institute of Neurosciences, University of Barcelona, 08036 Barcelona, Spain; August Pi i Sunyer Biomedical Research Institute (IDIBAPS), Barcelona, 08036 Spain; Basic Sciences Department, Faculty of Medicine and Health Sciences, Universitat Internacional de Catalunya, E-08195 Sant Cugat del Vallès, Spain; Centro de Investigación Biomédica en Red de Fisiopatología de la Obesidad y la Nutrición (CIBEROBN), Instituto de Salud Carlos III, E-28029 Madrid, Spain; Centro de Investigación en Red, Enfermedades Neurodegenerativas (CIBERNED), Instituto de Salud Carlos III, Madrid, Spain; Department of Biochemistry and Physiology, School of Pharmacy and Food Sciences, Universitat de Barcelona, 08028 Barcelona, Spain; Institut de Neurociències UB, Campus Mundet, Passeig de la Vall d’Hebron 171, 08035 Barcelona, Spain; Grupo de Terapia Celular y Medicina Regenerativa, Dpto. de Fisioterapia, Medicina y Ciencias Biomédicas. Facultad de Ciencias de la Salud, Universidad de da Coruña, INIBIC-CHUAC, Centro Interdisciplinar de Química y Biología (CICA), 15006, A Coruña, Spain; Institut de Nutrició I Seguretat Alimentària de la Universitat de Barcelona, Av. Prat de la Riba, 171, 08921 Santa Coloma de Gramenet, Spain; Functional Informatics Lab, National Center for Bioinformatics, Quaid-i-Azam University, Islamabad, Pakistan; Department of Pharmacology, Therapeutic and Toxicology, Universitat Autònoma de Barcelona, 08193 Barcelona, Spain; Health Research Institute of Santiago de Compostela (IDIS), University Hospital of Santiago de Compostela (SERGAS), Trav. Choupana s/n, 15706 Santiago de Compostela, Spain; Institute of Biomedicine of the University of Barcelona (IBUB), Universitat de Barcelona, 08028 Barcelona, Spain; Laboratorio de Epigenómica del Envejecimiento, Centro de Investigación sobre el Envejecimiento, Cinvestav Sede Sur, 14330, Mexico City, Mexico

**Keywords:** Alzheimer’s Disease, GSK-3β, G9a, BBB permeation, small molecule, SAMP8, Transcriptomic analysis

## Abstract

Epigenetic dysregulation and abnormal kinase signaling are key factors in Alzheimer’s disease (AD), yet no existing treatment targets both. We introduce a dual-targeting strategy that inhibits the histone methyltransferase G9a and the serine/threonine kinase GSK-3β—two synergistic drivers of neurodegeneration—using a single CNS-penetrant small molecule. This report covers the rational design, synthesis, and preclinical testing of T2 as a new therapeutic candidate. Structure-based pharmacophore modeling and medicinal chemistry refinement led to a focused library of dual inhibitors, with T2 standing out due to its balanced potency against both targets, ability to cross the blood-brain barrier, low off-target effects, and a favorable safety profile. In the SAMP8 mouse model of late-onset AD, T2 markedly improved memory, restored social behaviors, and increased synaptic complexity. Molecular studies showed that T2 treatment lowered tau phosphorylation, the Aβ42/40 ratio, and neurofilament light chain levels, while boosting neurotrophic factors and anti-inflammatory pathways. Multi-omics analyses indicated that T2 causes widespread transcriptional and proteomic changes, activating synaptic plasticity genes (like *Arc and Nectin3*), reducing ferroptosis and senescence-related pathways, and altering chromatin accessibility. Electrophysiological tests confirmed the restoration of AMPAR-mediated synaptic transmission. Overall, these results support dual G9a/GSK-3β inhibition as a promising disease-modifying approach for AD, with T2 emerging as a strong candidate for further therapeutic development.

## INTRODUCTION

Approximately 55 million people worldwide live with dementia, and nearly 10 million new cases arise each year. Alzheimer’s disease (AD), the most common form of dementia, makes up 60-70% of these cases and mainly affects the aging population.^1^ AD is a progressive neurodegenerative disorder marked by cognitive decline, memory loss, and behavioral changes. Its complex development involves multiple mechanisms, including the buildup of beta-amyloid (Aβ) plaques, tau neurofibrillary tangles, neuroinflammation, synaptic dysfunction, and epigenetic dysregulation.^2^

Epigenetic mechanisms have also become key contributors to AD pathology. G9a, a histone methyltransferase, catalyzes the methylation of histone H3 at lysine 9 (H3K9me2), leading to the repression of target gene transcription.^3^ Dysregulated G9a activity has been associated with cognitive deficits in AD through its effects on synaptic plasticity, neuronal survival, and gene expression.^4^ Notably, inhibiting G9a has been shown to reverse Aβ-induced memory impairments, highlighting its potential as a therapeutic target. Additionally, G9a plays important roles in autophagy and synaptic regulation, processes that are essential for maintaining neuronal integrity.^4,5^

Moreover, glycogen synthase kinase-3 beta (GSK-3β) is a serine/threonine kinase involved in various physiological functions, including cell regulation, glycogen metabolism, and apoptosis.^6^ In the context of AD, GSK-3β plays a crucial role in tau hyperphosphorylation, which contributes to neurofibrillary tangle formation, a key feature of AD pathology.^6,7^ The connection between GSK-3β activity and recently identified tau phosphorylation sites has provided new insights into disease progression. Therefore, inhibition of GSK-3β shows promise in reducing both Aβ deposition and tau phosphorylation in transgenic AD mouse models.^6,8^ Additionally, GSK-3β inhibition lessens Aβ-induced neurotoxicity in cultured neurons and influences neuroinflammatory responses, making it a promising therapeutic target.^9^

Interestingly, recent advances in tau biomarker research have significantly expanded our understanding of disease progression and diagnostic capabilities. The development of novel phosphorylated tau (p-tau) biomarkers, including p-tau212, p-tau205,^10,11^ brain-derived tau (BD-tau),^12^ and N-terminal tau fragments (NTA-tau),^13^ has revealed distinct temporal dynamics and biological significance in AD pathogenesis. Plasma p-tau217 has emerged as the most clinically validated biomarker, demonstrating exceptional diagnostic accuracy equivalent to cerebrospinal fluid biomarkers and superior performance compared to traditional imaging approaches.^14^ These biomarkers provide complementary information about different stages of tau pathology. While p-tau231 and p-tau217 detect the earliest amyloid-induced changes, p-tau205 emerges later and correlates strongly with tau-PET signal and Braak staging, positioning it as a sensitive proxy for neurofibrillary tangle burden.^10,14,15^ Novel microtubule-binding region tau fragments (MTBR-tau), particularly MTBR-tau243, have shown remarkable potential in bridging the gap between fluid and imaging tau biomarkers by reflecting ongoing pathological changes in brain tissue.^16^ Additionally, brain-derived tau (BD-tau) has demonstrated unique specificity for AD-type neurodegeneration, outperforming plasma total-tau and showing independence from peripheral tau sources, thereby completing the AT(N) biomarker framework in blood-based diagnostics.^12^ Furthermore, the integration of multiple tau biomarkers into comprehensive panels has revealed that different phosphorylation sites provide complementary temporal information: p-tau231 and p-tau217 indicate early amyloid-β changes before the onset of overt plaque pathology. At the same time, p-tau205 and MTBR-tau fragments reflect later-stage neurofibrillary pathology and ongoing neurodegeneration. This multi-biomarker approach has profound implications for clinical trial design, enabling more precise patient stratification, earlier intervention windows, and objective monitoring of therapeutic response. The development of dried blood spot sampling methods has further enhanced accessibility, potentially enabling population-scale screening and remote monitoring capabilities.^17^

Emerging evidence shows a functional interaction between G9a and GSK-3β, suggesting that epigenetic regulation can influence kinase signaling and tau pathology.^18^ This crosstalk might be especially important for regulating new tau phosphorylation sites, as epigenetic changes could alter the expression and activity of kinases responsible for site-specific tau modifications. Recent research indicates that G9a-driven histone modifications can affect the cellular processes involved in tau processing and secretion, potentially impacting the production of brain-derived tau fragments and MTBR-tau species, which are used as disease biomarkers. This interaction underscores the intricate relationship between epigenetic and signaling pathways in AD development. Targeting G9a could not only regulate gene expression but also indirectly influence GSK-3β activity, providing a multifaceted therapeutic approach. This approach, known as polypharmacology, involves targeting multiple biological pathways simultaneously to enhance treatment effectiveness for complex diseases. Therefore, dual inhibition of G9a and GSK-3β offers a promising strategy for creating effective therapies that address the multifactorial aspects of AD, with the potential to slow cognitive decline and enhance patient outcomes.

Advanced tau biomarker research has significant therapeutic implications that go beyond diagnosis, encompassing treatment monitoring and drug development.^19,20^ Detecting specific tau forms, like BD-tau and MTBR-tau fragments, allows for highly precise evaluation of tau-targeted therapies. These biomarkers can act as pharmacodynamic endpoints in clinical trials involving dual-target inhibitors, such as G9a/GSK-3β modulators, providing real-time insights into drug engagement and therapeutic effects. Additionally, the distinct temporal patterns of tau biomarkers can support personalized treatments: early biomarkers (p-tau231, p-tau217) help identify candidates for prevention, while later markers (p-tau205, MTBR-tau) assess therapeutic response in symptomatic patients. ^10,12–14,16,19–22^

In this study, we report a rationally designed dual-inhibitor strategy targeting G9a and GSK-3β. We synthesized and characterized a series of tetrahydropyrimidine-based compounds. Among these, compound T2 emerged as a promising lead, characterized by high selectivity, brain permeability, and favorable pharmacokinetic and safety profiles. T2 demonstrated robust neuroprotective effects in cellular and animal models of AD, reducing Aβ aggregation and tau hyperphosphorylation, enhancing synaptic plasticity, and modulating gene expression. Importantly, T2 treatment resulted in significant reductions in plasma levels of multiple tau phosphorylation sites (p-tau181, p-tau217, p-tau231), providing direct evidence of therapeutic efficacy using clinically validated biomarkers. The biomarker profile observed with T2 treatment suggests modulation of both early amyloid-associated tau changes and later neurofibrillary pathology, consistent with the dual-target mechanism of action. These findings provide compelling preclinical evidence supporting the therapeutic potential of simultaneous G9a and GSK-3β inhibition in the treatment of AD.

## RESULTS

To validate our polypharmacological strategy targeting G9a and GSK-3β, we conducted a comparative analysis of post-mortem cortical brain tissue from AD patients and age-matched non-demented (ND) controls. As previously reported by our group,^3^ G9a protein levels were significantly elevated in the AD cohort relative to ND individuals (Figure 1a–b), supporting its pathological upregulation in AD. Simultaneously, the active form of GSK-3β, phosphorylated at tyrosine 216 (p-GSK-3β Tyr216), was also increased in AD brains. The ratio of p-GSK-3β Tyr216 to total GSK-3β was significantly higher in AD samples (Figure 1a, c–e), indicating enhanced kinase activity. Additionally, elevated levels of phosphorylated tau at disease-relevant residues ( T231, T217, T181, and S202/T205) were observed (Figure 1f-g). Phosphorylations like p-T231, p-T217, and p-T181 start to rise in the cerebrospinal fluid at the same time as the initial buildup of Aβ aggregates.^14^ In contrast, phosphorylation at T205 may occur later and appears to be more closely linked to tau-PET imaging findings.^10,20^ Together, these findings substantiate the pathological activation of both G9a and GSK-3β in human AD brain tissue, supporting their relevance as dual therapeutic targets.

**Figure 1.**
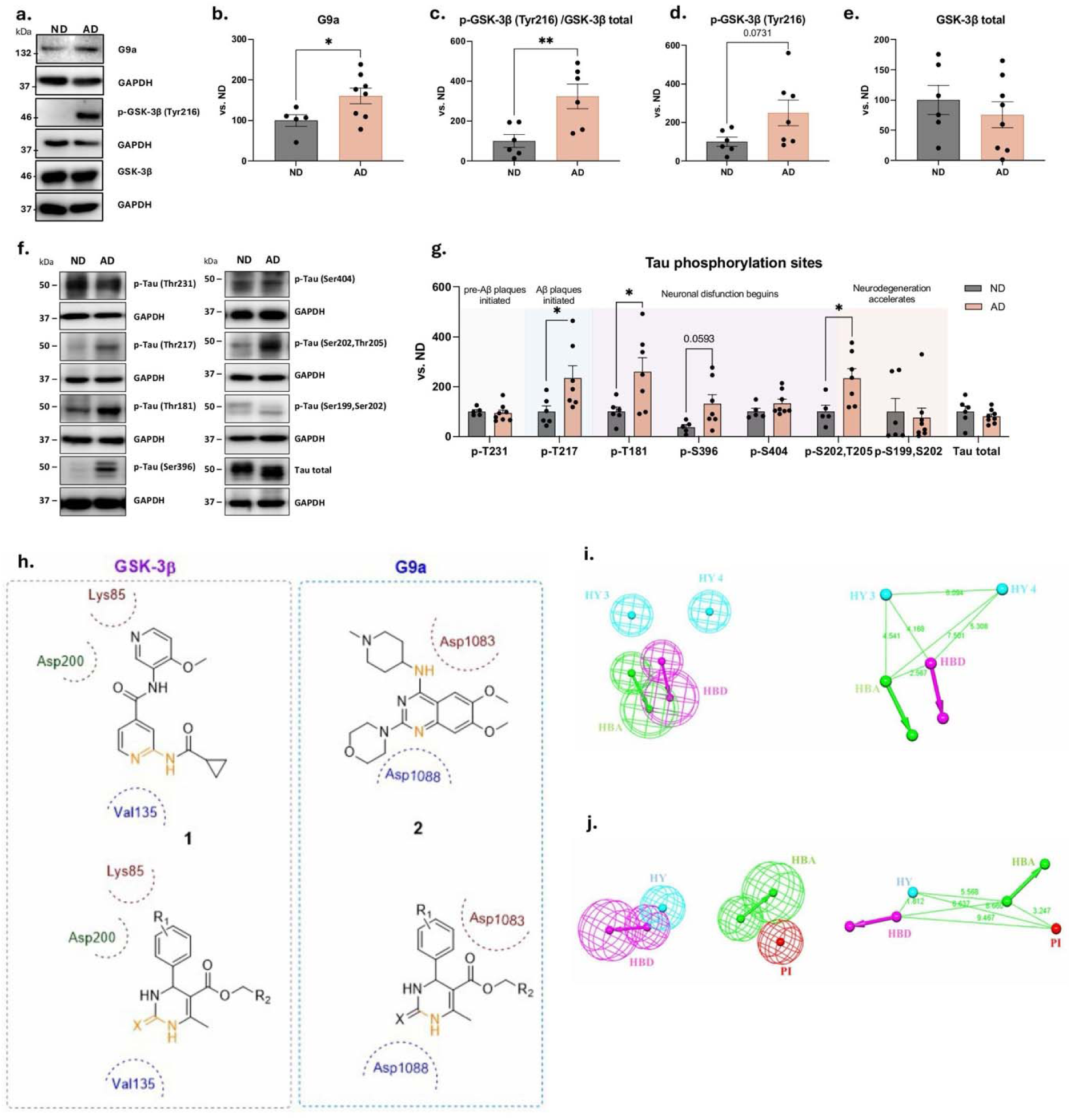
a. WB representative images, and **b.** quantification of G9a*, **c.** p-GSK-3β (Tyr216)/ GSK-3β total, **d.** p-GSK-3β (Tyr216) and **e.** GSK-3β total level in AD patients compared to ND patients. **f.** WB representative images, and **g.** quantification of Tau phosphorylation landscape in AD and ND human patients. Values presented are mean ± SEM; (ND n = 5, and AD n = 7-8); Statistical analysis: Student’s t-test; *p<0.05;**p<0.01. **h.** Design strategies for tetrahydropyrimidines as GSK-3β/G9a dual inhibitors. **I-J.** Generated 3D QSAR pharmacophores against GSK-3β (i) and G9a (**j)**. Both pharmacophores contain three identical features, hydrogen bond acceptor (HBA), hydrogen bond donor (HBD), and hydrophobic (HY), in common. Importantly, the HBA and HBD features are closer (2.5 Å) in space in GSK-3β pharmacophore, while apart in space (∼ 6 Å) in G9a pharmacophore. *Figure 1a-b reproduces data previously reported by our group, included here to provide a cohesive narrative within the present manuscript.^3^

### Design strategy for dual G9a/GSK-3β inhibitors

Multitarget drug design was approached through a fragment-based design strategy, allowing us to identify shared pharmacophoric features between known inhibitors of G9a and GSK-3β, including compounds from our previous studies. Through this comparative analysis, the tetrahydropyrimidines scaffold emerged as a potential starting point (Figure 1h) in dual design hypothesis.

To the inclusion of common pharmacophoric features present in tetrahydropyrimidines derivatives, we carried out advanced 3D QSAR based pharmacophore generation of G9a and GSK-3β, respectively. A total of sixteen compounds with IC_50_ values ranging from 0.9nM to 18,200nM were used as the training set to generate 3D pharmacophore models of G9a. In comparison, twenty-one training set compounds with IC_50_ values ranging from 5nM to 10,000nM were used to develop pharmacophore models of GSK-3β. The pharmacophore models were built and ranked based on their statistical parameters using the HypoGen algorithm of the commercially purchased BIOVIA Discovery Studio Academic Research Suite.^23^ The final two pharmacophores against G9a and GSK-3β were selected through proper validation methods (internal validation using statistical methods, and external validations via predicting activities of the test set) as explained in detail in the Supplementary information. The comparative analysis of the pharmacophores of G9a and GSK-3β was carried out to gain insight into the features required in the 3D space for the dual inhibition hypothesis (Figure 1i-j). As shown in Figure 1i-j, it was concluded that the 3D pharmacophores of G9a and GSK-3β contain three identical features, hydrogen bond acceptor (HBA), hydrogen bond donor (HBD), and hydrophobic (HY), in common. However, the HBA and HBD features are closer (2.5Å) in space in GSK-3β pharmacophore and can be complemented by the opposite features present in tetrahydropyrimidine ring itself. On the contrary, the HBA and HBD features are apart (∼ 6 Å) in space in the G9a pharmacophore. While the tetrahydropyrimidine ring –NH can complement the HBD feature in G9a, still one additional HBA feature (*e.g.,* ester moiety in our design) is required to satisfy the ∼6 Å distance in space from the HBD feature for the efficient interaction. The common hydrophobic (HY) features present in both pharmacophores were covered by putting methyl/phenyl group attached to the ring in our design. Based on the above-said dual hypothesis using common tetrahydropyrimidine ring and comparative analysis of the 3D QSAR pharmacophores, we designed a series of tetrahydropyrimidine derivatives having suitable functionalities, ester and hydrophobic moieties at suitable distance, for the dual inhibition G9a and GSK-3β.

Having identified the tetrahydropyrimidine core as a suitable scaffold for achieving dual inhibition, we next built two representative collections of tetrahydropyrimidine derivatives, featuring a urea or a thiourea group, and an ester (Series 1) or an amide (Series 2) functionality (see Table S1 and S2). These compounds were either commercially available or readily synthesized in just one synthetic step by a Biginelli reaction from a benzaldehyde derivative, a urea (or thiourea), and a β-ketoester or a β-ketoamide (see Supplementary information for details). All the compounds were fully characterized, and details are included in the Supplementary Information file.

### *In vitro* and *in silico* selection of the lead

To achieve effective dual-target inhibition, it is critical to attain a balanced potency toward both G9a and GSK-3β. We therefore assessed the inhibitory activity of all synthesized compounds under biochemical assay conditions. Several compounds demonstrated a relevant range of activity at both targets. Notably, compounds T2, T3, and T4 emerged as potential good candidates due to their dual-target activity (Table S1-S2, Figure S1).

Off-target inhibition of GLP, a homologous histone methyltransferase, was evaluated to assess compound selectivity. Most of the tested compounds showed <50% inhibition of GLP at the evaluated concentration of 10µM, representing a very favorable selectivity profile compared to the reference inhibitor UNC0638 (GLP IC_50_ = 19nM).^24^ Among them, compounds T2, U1, U5, and CT2 showed the lowest GLP inhibition, indicating enhanced specificity for G9a (Table S1-S2, Figure S1).

To determine the CNS drug-likeness of the synthesized molecules, we assessed blood–brain barrier (BBB) permeability using the PAMPA-BBB assay. Compounds from Series 1 generally exceeded the permeability threshold established by Di et al. (Pe > 5.16 × 10⁻L cm·s⁻¹), suggesting adequate CNS exposure. In contrast, Series 2 compounds displayed lower permeability. For example, compounds T1, T3, T4, and U4, from Serie 1, were more permeable than their counterparts from Serie 2, CT1, CT3, CT4, and CU4, respectively. As a result, subsequent investigations focused exclusively on the Series 1 family (Table S1).

Pharmacokinetic and toxicity profiles were evaluated to prioritize compounds with optimal CNS drug properties. Parameters such as human plasma and microsomal stability, plasma protein binding, hERG channel inhibition, and CYP450 inhibition were systematically analyzed (see Supporting Information for details, Table S3-S5).

To assess functional efficacy, *Caenorhabditis elegans* (*C. elegans*) CL2006 transgenic strain, which expresses human AβL–LL in muscle cells and develops age-dependent paralysis^25^ were treated with the dual inhibitors. Multiple compounds, including T1, T2, T12, U2, U4, U6 and U13 (Figure 2a-n), significantly improved locomotion, with T2 exhibiting the most robust dose-dependent effect. Furthermore, Aβ aggregation was reduced by all candidates, except Tideglusib, with T2 showing the strongest effect (52% reduction, Figures 2o-p). These results position T2 as the lead compound for preclinical development.

**Figure 2.**
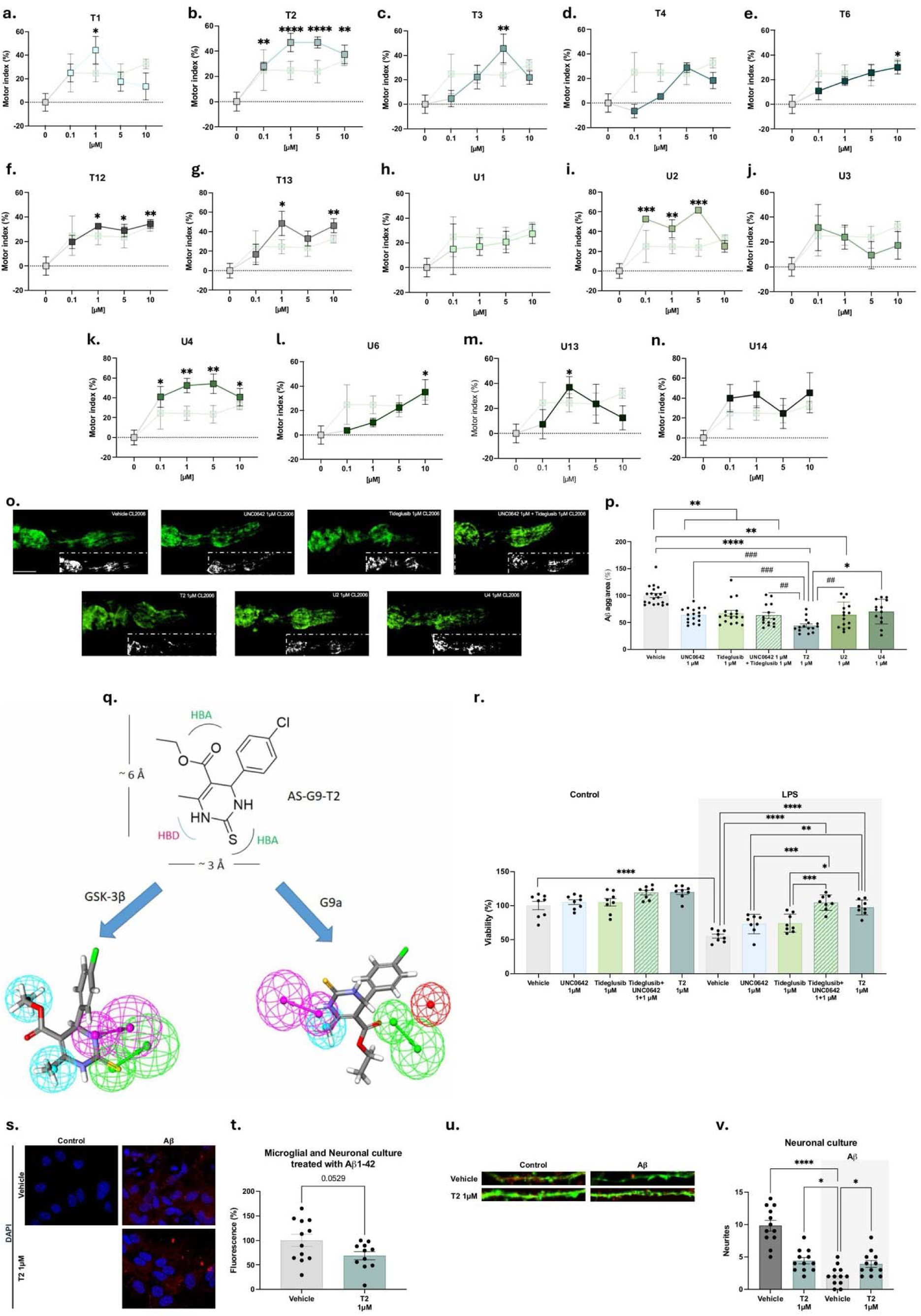
a.-n. Treatment with the inhibitors ameliorates the motor dysfunction of CL2006 worms. For dose-response: Values represented are mean ± Standard error of the mean (SEM); n=3 with at least 90–100 worms in each group. Statistics: One-Way ANOVA, followed by Tukey post-hoc analysis. *p<0.05, **p<0.01, ***p<0.001, ****p<0.0001. **o.** and **p.** Quantification of ThS-positive particles in the head region of CL2006 strain, and representative images. Values represented are mean ± Standard error of the mean (SEM); n=3 with 25 worms in each group. Statistical analysis: Kruskal–Wallis test followed by Dunn’s post-hoc analysis, *p<0.05, **p<0.001, ****p<0.0001; Mann-Whitney test, ^##^p<0.01, ^###^p<0.001. **q**. Mapping the designed lead into the G9a and GSK-3β pharmacophores separately reveals complementary interactions between the features within the specified distance in 3D space, as predicted by our dual design hypothesis. In the GSK-3β pharmacophore, the ring– NH and the thioketo moieties of the designed lead complemented the HBD and HBA features within < 3 Å distance. In the G9a pharmacophore, the ring–NH complemented the HBD feature, and the incorporated ester moiety provided the HBA feature at a proper distance (∼6Å) for efficient interaction. The HY features of both the pharmacophores were complemented by incorporating methyl/phenyl rings to the tetrahydropyrimidine core in our design. **r.** Cell viability after LPS insult. **s.** Representative images and **t.** Quantification of the effect of T2 (1µM) on Aß_1-42_ aggregation in primary neuronal and microglial cultures. Data are presented as the mean ± SEM of 5 independent experiments. Statistical analysis: One-way ANOVA; *p < 0.05 (n ≥ 3). **u.** and **v.** Neurite formation was analyzed in primary neuronal cultures treated with Aβ_1-42_. Data are presented as the mean ± SEM of 5 independent experiments. Statistical analysis: One-way ANOVA; *p < 0.05 (n ≥ 3).

### *In silico* validation of T2 binding via pharmacophore modeling and molecular dynamics

The lead compound, T2, was mapped independently onto the pharmacophore models of G9a and GSK-3β (Figure 2q), revealing complementary spatial interactions consistent with our dual-inhibition design hypothesis. As shown in Figure 2q, the ring–NH and the thioketo moieties of T2 complemented the HBD and HBA features within < 3 Å distance into the GSK-3β pharmacophore. The ring –NH complemented the HBD feature, and the incorporated ester moiety provided the HBA feature at a proper distance (∼ 6Å) for the efficient interaction of T2 with the G9a pharmacophore. The HY features of both the pharmacophores were complemented by the incorporated methyl groups of T2.

The molecular dynamics (MD) simulation of G9a-T2 and GSK-3β-T2 complex was conducted over 200 ns to evaluate protein stability, binding modes, protein–ligand interactions, and functional characteristics. The RMSD of the GSK-3β-T2 complex was stable throughout the simulation duration and is stable around 1.8 Å (Figure S2a). The RMSD for the T2-bound substrate site G9a-T2 complex was stabilized after 60 ns and remained stable until 160 ns at 10 Å. At 175 ns, it dropped to 6Å and remained stable around this value until the end of the simulation (Figure S2b). While in the case of T2 bound at the SAM site, the G9a-T2 complex RMSD remains unstable throughout the simulation (Figure S2c).

To evaluate the stability of amino acid residues in the binding pocket, the RMSF of the protein backbone was calculated for the entire simulation duration. Fluctuation of amino acid residues in the binding pocket of GSK3β-T2 complex (Figure S2d), as well as for T2 bound at the substrate site G9a-T2 complex (Figure S2e), remained stable below 2.5 Å (denoted by green line) (Figure S2d). While in the case of T2 bound at the SAM site, G9a-T2 complex binding pocket amino acid residues are volatile and fluctuating up to 7.5 Å (Figure S2f).

T2 series is already reported as an ATP competitive GSK-3β inhibitor in our previous research^18,19^, and the stability of protein ligand complex throughout simulation also proves that T2 binds at the ATP site. In the case of G9a, the stability of T2 bound at the substrate site and the instability of T2 bound at the SAM site indicate that the G9a-T2 complex proves T2 is a substrate-competitive G9a inhibitor, a finding further validated by *in vitro* studies.

### Neuroprotective Activity in Mixed Neuronal Cultures

To model neuroinflammation, primary cortical neurons and microglia were exposed to lipopolysaccharide (LPS) (1 µM) and 200 U/mL interferon-γ (IFN-γ) for a period of 48 hours. Thereafter, the cells were stimulated with our dual approach (T2 or UNC0642 and Tideglusib), either singly or in combination, as well as with vehicle, for an additional 48 hours. Finally, a combination of single inhibitors (UNC0642 and Tideglusib) was used. The dual approach (UNC0642 + Tideglusib) and T2 alone significantly improved cell survival, exceeding the efficacy of single-agent treatments (Figure 2R). Furthermore, T2 reduced Aβ aggregation and restored neurite complexity, thereby reversing the disruptive effects of AβL–LL exposure (500nM, 48h) *in vitro* (Figure 2s–v).

### Preclinical profiling of T2 covering selectivity, safety, and pharmacokinetics

To evaluate the off-target profile of the lead dual inhibitor T2, a comprehensive selectivity panel was conducted against a range of epigenetic and enzymatic targets at a concentration of 10 µM (Figure 3a). T2 exhibited minimal inhibition (<50%) against the majority of other histone methyltransferase family members, including SETD7, SUV39H1, PRMT5, and DOT1L (Figure 3b). Additionally, T2 showed high specificity across a broad kinase panel, with residual activity toward unrelated kinases, such as CDKs, MAPKs, and PI3K isoforms, remaining above 70%, reinforcing its clean selectivity profile.

**Figure 3.**
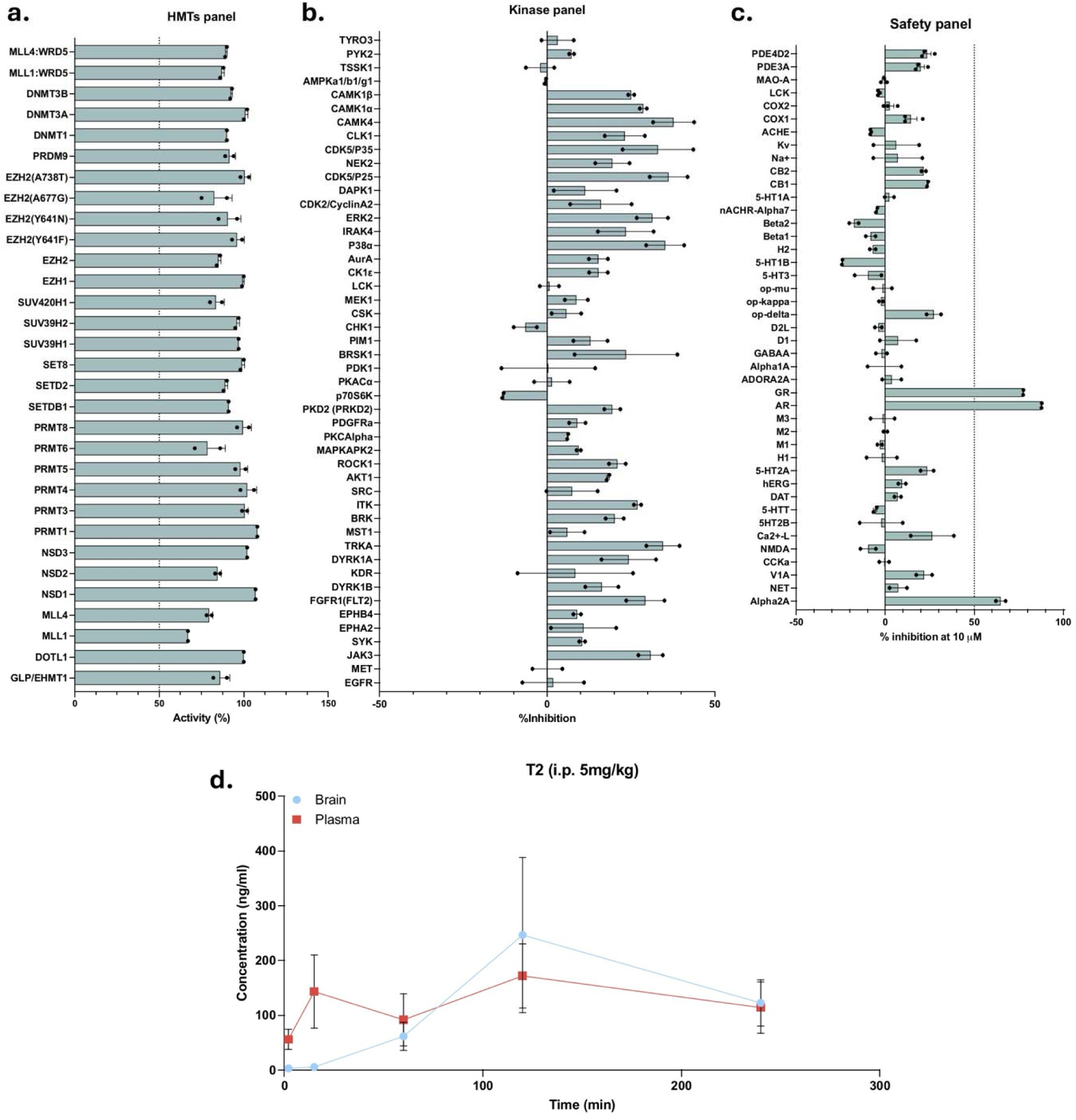
Selectivity, safety, and permeability profile of T2. **a.** Histone methyltransferases, and **b.** kinase and. **c.** safety panel of T2 at 10µM. **d.** Timeline of the T2 plasma and brain concentrations (ng/mL) after i.p. administration at 5mg/Kg.

Given the necessity of long-term exposure in chronic central nervous system (CNS) conditions such as AD, establishing preclinical safety is a critical prerequisite. A safety pharmacology panel at 10 μM revealed negligible inhibition across several key biological targets. Specifically, COX-1, COX-2, voltage-gated Na⁺ and K⁺ channels, 5HT1A, dopamine D1, hERG, DAT, NET, NK1, and TRPV1 receptors showed <20% inhibition, indicating a low risk of interaction with these systems. At the same concentration, T2 displayed >50% inhibition toward glucocorticoid receptor (GR), androgen receptor (AR), and alpha-2A adrenergic receptor. However, these findings were observed at concentrations tenfold higher than the effective dose, suggesting a limited *in vivo* risk under therapeutic conditions (Figure 3c). Further *in vitro* mutagenic (Ames test) and genotoxicity assessment (micronucleus test (MNT)) confirmed that T2 does not contain base-pair or frame-shift mutagens and does not induce chromosomal damage at relevant concentrations, supporting its classification as a non-mutagenic and non-genotoxic compound (Table S6-S7, Figure S3a-d). Furthermore, to limit the maximal tolerated dose (MTD), an acute oral toxicity assay was conducted. Accordingly, the MTD in mice of T2 orally administered was 1 g/Kg (Figure S3e-k).

Next, we validated the CNS druggability of orally administered T2 at 5 mg/Kg through an in-depth pharmacokinetic assessment (Figure 3d). The concentration of T2 in the brain (blue circles) increased gradually, reaching a peak at 2 hours post-dose, followed by a slow decline, indicating sustained central exposure. In contrast, plasma concentrations exhibited a more rapid rise and fall, with an earlier peak and lower overall exposure relative to brain levels. Notably, brain concentrations exceeded those in plasma at intermediate and later time points, suggesting efficient penetration across the blood–brain barrier and possible retention within brain tissue. By the final time point (4 hours), concentrations in both compartments converged. These results highlight the favorable brain distribution of T2 and confirm its potential as a therapeutic candidate for targeting CNS disorders.

### T2 treatment improves age-related behavioral abnormalities and cognitive decline in SAMP8 mice

To assess efficacy, we employed the senescence-accelerated mouse-prone 8 (SAMP8) mouse model, widely recognized as a model of late-onset Alzheimer’s disease (LOAD). As a first step, we confirmed the disease-relevant expression profile of the targets: both G9a expression and the p-GSK-3β/total GSK-3β ratio were significantly increased in SAMP8 mice compared to senescence-resistant SAMR1 controls (Figure 4a-c, S4a-b), validating the model’s suitability for proof-of-concept (PoC) evaluation. SAMP8 mice were then treated with T2 (5 mg/kg, administered by oral gavage daily for 1 month). Cognitive performance was assessed using the Novel Object Recognition Test (NORT) and the Object Location Test (OLT), which evaluate short-term and long-term memory, respectively. Exploration times during the habituation phases did not differ across groups (Figures S4c-d). T2 treatment led to significant improvements in both short-and long-term memory performance in NORT (Figures 4d-e) and OLT compared to untreated SAMP8 controls (Figure 4f).

**Figure 4.**
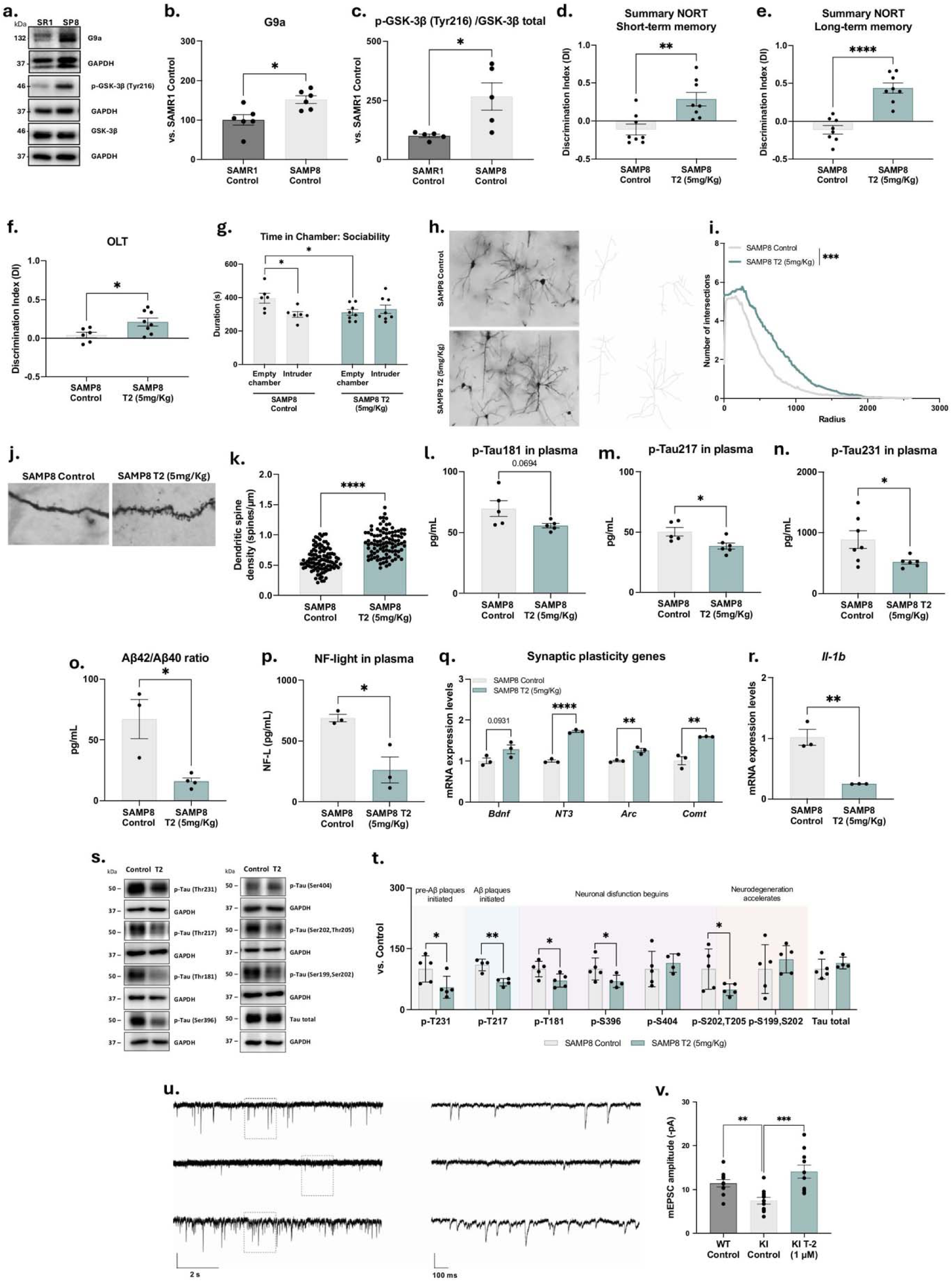
T2 efficacy in the SAMP8 mouse model. **a.** WB representative images, and **b.** quantification of G9a*, **c.** p-GSK-3β (Tyr216)/ GSK-3β total levels in SAMP8 compared to SAMR1. Values presented are the mean ± SEM (SAMR1 Control, n = 5-6, and SAMP8, n = 5-6). Statistical analysis: Student’s t-test analysis; *p < 0.05. **d.** and **e.** NORT evaluation by DI after 2h and 24h, respectively. **f.** OLT evaluation by DI after 24 h. **g.** Time in chamber: Sociability in the three-chamber test. Values presented are the mean ± SEM; (SAMP8 Control n = 8, and SAMP8 T2 (5mg/Kg) n = 8); Statistical analysis: Student t-test; *p<0.05; **p<0.01; ****p<0.0001.**h.** Representative images of Golgi-stained neurons (scale bar = 100 µm). **i.** Number of neuronal intersections. Values presented are the mean ± SEM; (N = 2 groups (SAMP8 Control n = 90, and SAMP8 T2 (5mg/Kg) n = 90); Statistical analysis: Kolmogorov-Smirnov; p-value represented in the graph. **j.** Representative images of the spine density. **k.** Spine density quantification of neurons. Values presented are the mean ± SEM; (N = 2 groups; SAMP8 Control n = 60, and SAMP8 T2 (5mg/Kg) n = 60 dendrites (of different neurons) were analyzed from 6 mice per group; Statistical analysis: Student t-test; ****p<0.0001. **l.** Plasma levels of p-TauT181, **m.** p-TauT217, **n.** p-TauT231, **o.** Aβ_42_/Aβ_40_ ratio, **p.** NF-light in SAMP8 treated with both vehicle and T2. **q.** Gene expression of *Bdnf*, *NT3*,Arc, *Comt*, *and* **r.** *Il-1b.* **s.** WB representative images, and **t.** Quantification of the Tau phosphorylation landscape in SAMP8 and SAMP8 mice treated with T2. Values presented are mean ± SEM; (SAMP8 Control n = 5-4, and SAMP8 T2 (5mg/Kg) n = 5-4); Statistical analysis: Student’s t-test; *p<0.05; **p<0.01; ****p<0.0001. **u.** Representative traces of AMPAR-mediated miniature EPSCs recorded from hippocampal neurons. Recordings were obtained from wild-type neurons treated with vehicle (WT-DMSO; top traces), from knock-in mice neurons treated with vehicle (KI-DMSO; middle traces), and from knock-in neurons treated with 1 μM T2 for 72 hours before recordings (KI-T2; lower traces). The left panels display 10-second continuous recordings, while the right panels display magnified segments corresponding to the dotted boxes in the left panels. **v.** Quantification of AMPAR-mediated mEPSC amplitudes under the three experimental conditions shown in panel u. The bar graph represents the mean ± standard error of the mean (SEM) for each group. Individual data points (open dots) indicate the number of individual recordings: 10 for WT-DMSO, 11 for KI-DMSO, and 10 for KI-T2, collected from three independent neuronal cultures. *Figure 4a-b reproduces data previously reported by our group, included here to provide a cohesive narrative within the present manuscript.^3^

Social behavior was evaluated using the Three-Chamber Test (TCT). No changes were observed during the habituation phase (Figure S4e). However, SAMP8 control mice showed reduced time in the chamber with the intruder, confirming the social withdrawal phenotype. T2-treated mice showed a significant reduction in time spent in the empty chamber, indicating partial restoration of social engagement (Figure 4g). Notably, sniffing time toward the intruder remained unchanged (Figure S4f), suggesting a specific effect on social preference rather than olfactory-driven investigation.

Morphological analysis revealed a significant increase in neuronal arborization (intersections) and spine density in the T2-treated group compared to the control (Figures 4h-k), consistent with enhanced synaptic plasticity.

Given the central involvement of tau and Aβ in AD pathology, we evaluated key molecular markers. T2 treatment reduced phosphorylation of tau at multiple disease-relevant sites (T181, T217, T231) (Figures 4l-n), as well as the Aβ_42_/Aβ_40_ plasma ratio and neurofilament light chain (NF-L) levels (Figures 4o-p, S4g-h), suggesting an attenuation of neurodegeneration. Additional tau hyperphosphorylation sites (S396, S202/T205) in brain tissue were also reduced following treatment (Figures 4s-t).

To elucidate mechanisms of synaptic repair, we analyzed neurotrophic and plasticity-related gene expression. Levels of *Brain-derived neurotrophic factor (Bdnf)* and *neurotrophin 3 (NT3)* were increased in SAMP8-treated mice, promoting synapse formation in response to learning (Figure 4q). We also demonstrated an increase in the gene expression of immediate early genes (IEGs) involved in memory formation, such as *activity-regulated cytoskeleton-associated protein (Arc),* after T2 treatment (Figure 4q)*. Catechol-O-methyltransferase (COMT),* engaged in working memory, was also found to increase in the SAMP8-treated group (Figure 4q). In addition, we also examined the gene expression of pro-inflammatory cytokines, such as *interleukin*-*1*β *(Il*-*1*β), which was observed to be reduced after T2 treatment (Figure 4r).

Finally, to assess the functional consequences of dual G9a/GSK-3β inhibition on synaptic transmission, we recorded AMPA receptor–mediated miniature excitatory postsynaptic currents (mEPSCs) in primary cortical neurons from knock-in (KI) mice following 24-hour T2 treatment. Pharmacological isolation of AMPAR-mEPSCs revealed a significant increase in mEPSC amplitude, indicative of enhanced postsynaptic AMPAR expression, while mEPSC frequency remained unchanged, suggesting that presynaptic function was unaffected (Figures 4u-v).

### T2 is a potent H3K9me2 inhibitor that reshapes chromatin landscape

To assess the effect of T2 in chromatin, we performed H3K9me2 ChIP-seq analyses in SAMP8 control and SAMP8 T2-treated samples. After observing significant peaks in both samples, we notice a global decrease in H3K9me2 (Figure 5a), with most peaks exhibiting a negative fold change in the T2-treated samples (Figures 5b-c). We next focus on identifying peaks with a significant enrichment decrease in T2-treated samples and found 326 with a greater than 2-fold decrease (Figures 5d-e). We next analyzed the closest genes to those decreased peaks and performed enrichment analysis to investigate their function. We found that genes close to H3K9me2 altered peaks are enriched with genes responsive to TGFb, which is an essential modulator of immune and neural cells (Figure 5f). Furthermore, we demonstrated that H3K9me2/H3 total levels were decreased after T2 treatment in the SAMP8 group compared to the Control group (Figure 5g-h).

**Figure 5.**
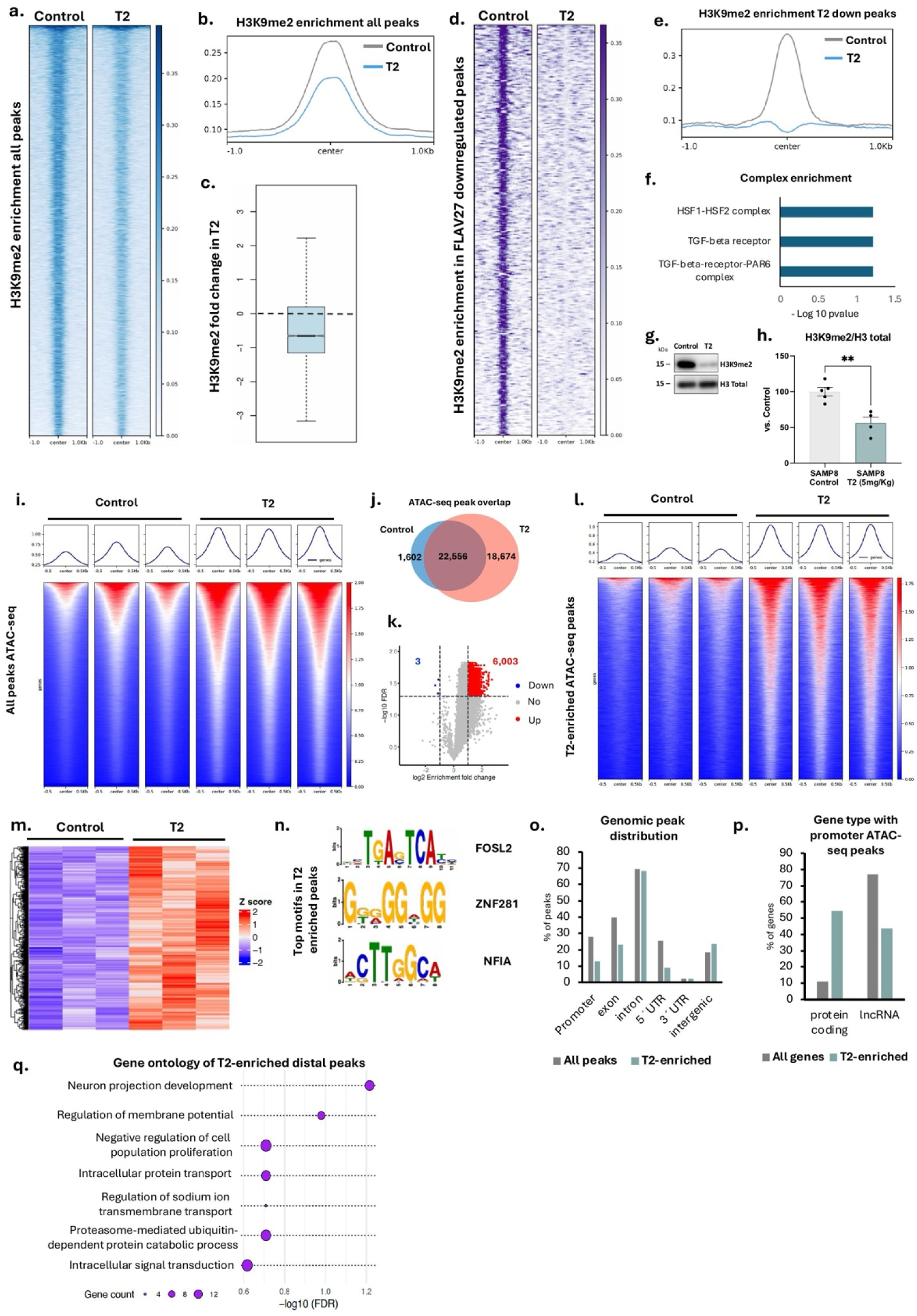
ChIP-seq and ATAC-seq analyses of T2 effects in cortical tissue. **a.** H3K9me2 enrichment at significant peaks in Control (left) and T2-treated (right) SAMP8 mice. **b.** Average H3K9me2 enrichment of control (gray) and T2 (blue) samples. **c.** Fold change in H3K9me2 enrichment of T2 compared to control. The dashed line indicates 0 (no fold change). Values below 0 indicate a decrease in T2, while values above 0 indicate an increase. **d.** H3K9me2 enrichment at peaks with a considerable (fold change > 2) decrease in T2 samples. **e.** Average H3K9me2 enrichment at peaks with a considerable decrease in T2 samples. **f.** Gene set enrichment analysis of genes near peaks with a significant decrease in H3K9me2 in T2 samples. **g.** WB representative images, **h.,** and quantification of H3K9me2 in SAMP8. Values presented are the mean ± SEM (N = 2 groups: SAMP8 Control, n = 5, and SAMP8 FLAV-27 (5 mg/kg), n = 4). Statistical analysis: Student’s t-test; **p < 0.001. **i.** ATAC-seq (accessibility) enrichment at significant peaks in Control (left) and T2-treated (right) SAMP8 mice. Heatmaps show enrichment at all peaks (bottom) while line plots show average enrichment (top). **j.** Venn diagram of overlapping peaks between control and T2 samples.**k.** Volcano plot of differentially enriched ATAC-seq peaks comparing T2 vs control samples. Numbers indicate the number of peaks with a significant (FDR < 0.05, FC > 2) change for peaks with increased (red) or decreased (blue) accessibility in T2. **l.** ATAC-seq enrichment at peaks with a significant increase in T2 (T2-enriched). **m.** Heatmap of average ATAC-seq enrichment at peaks with a significant increase in T2. **n.** DNA motifs enriched (Evalue < 0.05) in T2-enriched peaks. **o.** Genomic distribution of T2-enriched peaks (black) and all ATAC-seq peaks (grey). Results are presented as percentages with respect to the total peak number. **p.** Gene type of genes associated with T2-enriched peaks at promoters (grey) and all genes (blue). **q.** Gene ontology analysis of biological functions of genes associated with distal T2-enriched peaks. The X-axis shows significance (-log10(FDR), and the circle size corresponds to the number of genes in each category.

To further analyze if T2 has a global effect on chromatin conformation beyond H3K9me2 changes, we performed ATAC-seq in triplicate from control and T2-treated SAMP8 mice. We found significant peaks for all samples and analyzed chromatin accessibility. As expected, we found a clear increase in accessibility in T2-treated samples (Figure 5i), which again suggests that T2 has a global effect on chromatin conformation. Interestingly, the increase in accessibility is mostly composed of new peaks in T2 samples, while most peaks in control samples are maintained in T2-treated mice (Figure 5j). Moreover, confirming that T2 is an inhibitor of heterochromatin, most significantly changed peaks in T2-treated samples are peaks that show an increase (6,003) rather than a decrease (3) in accessibility (Figure 5k). Thus, we predict that T2 will have a significant effect in inducing different gene sets, rather than by increasing accessibility at promoters or distal regulatory regions. To analyze this, we focus on peaks that exhibit a significant increase in accessibility in T2-treated samples, which we refer to as T2-enriched peaks (Figure 5l-m). We first performed motif analysis of these peaks and found three significant motifs that are similar to binding sites of transcription factors FOSL2, ZNF281, and NFIA (Figure 5n). We then analyze the genomic distribution of T2-enriched peaks compared to all ATAC-seq peaks. Surprisingly, T2-enriched peaks are more frequent in intron an intergenic region than in promoter regions (Figure 5o). This suggests that the phenotypic effect of T2 may derive mainly from the derepression of distal regulatory elements. To dissect the effect of T2 treatment on direct promoter regulation and distal element regulation, we separated our T2-enriched peaks into those overlapping with gene promoters and those present in intergenic regions. We first analyzed the genes associated with promoter T2-enriched peaks. Interestingly, most genes with T2-enriched peaks at promoters are coding rather than non-coding genes, compared with the distribution of all genes (Figure 5p). We then performed GO analysis and found that promoter T2-enriched genes belong to global categories such as regulation of DNA replication or protein ubiquitination (data not shown or supp), which do not fully explain T2 role as a neuroprotector. We next looked at genes close (<35 kb) to T2-enriched distal peaks and repeated our GO analysis. This time, we found that genes associated with distal T2-enriched genes are involved in neural projection development and the regulation of membrane potential. This suggests that most of the neuroprotective effects we observe may be due to distal regulatory regions that are opened upon T2-treatment, rather than promoters. However, further studies are needed to confirm this hypothesis (Figure 5q).

### Single-cell RNA-seq reveals a neuroprotective and anti-inflammatory effect in T2-treated mice

To understand the transcriptomic effects of T2 treatment at the molecular level, we performed single-cell RNA-seq experiments in control and T2-treated SAMP8 mice. After removing cells with high mitochondrial content and those with very high RNA and feature counts, we analyzed 5,696 cells in the control samples and 9,550 in the T2-treated samples. We perform clustering analysis, UMAP projections, and annotation of cell identity using singleR (Figure 6a). We analyzed the cell type distribution and found that, accordingly to all our data, T2-treated mice have a higher number of neurons and a lower number of microglial cells (Figure 6b). These data suggest that T2-treatment has neuroprotective and anti-inflammatory effects. To confirm this, we analyzed marker genes specific to T2-cells and found that they are correlated with axon guidance (which is consistent with our ATAC-seq findings) (Figure 6c-d). Gene Ontology (GO) analysis of the upregulated gene set prominently highlighted pathways associated with synaptic structure, neurotransmission, and plasticity. Among the top-ranked terms was “vesicle-mediated transport in synapse”, with a strong statistical enrichment (adjusted p < 1e-25).

**Figure 6.**
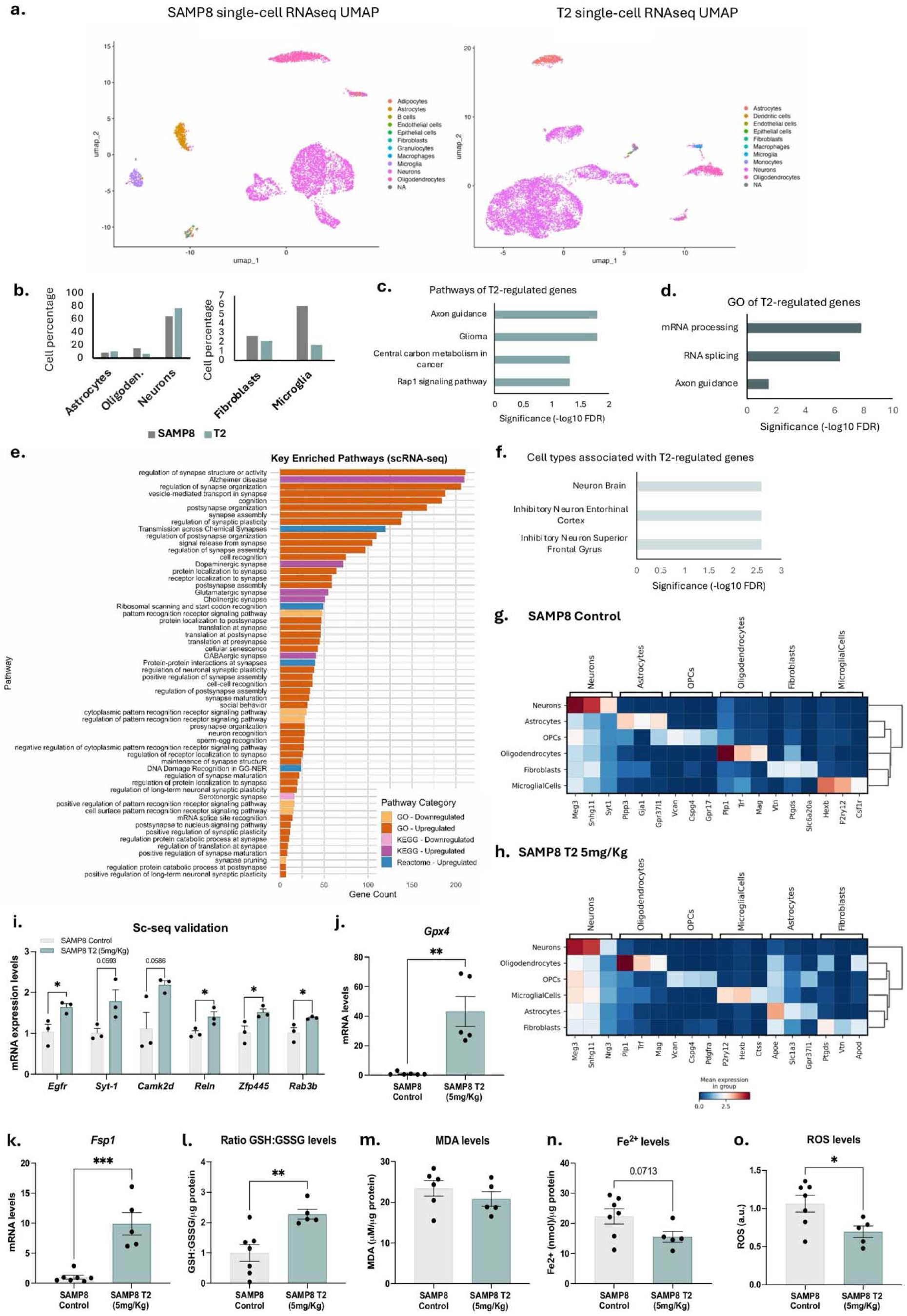
Single-cell RNA-seq in control and T2 mice. **a.** UMAP plots of control (left) and T2-treated mice (right) derived from single-cell RNA-seq data. **b.** Percentage of cells corresponding to different cell categories in control (gray) and T2 (blue) according to single-cell RNA-seq data. **c.** Enrichment analyses showing enriched pathways, **d.** Gene ontology biological process, **e.** GO, KEGG, and Reactome analyses and **f.** Cell type of marker genes specific to T2-treated cells. **g.** Expression heatmap of gene markers for each cell type population in control and **h.** T2-treated samples. **i.** qPCR validation of genes with significant changes in different single-cell clusters: *Egfr, Syt-1*, *Camk2d*, ReIn, *Zfp445,* and *Rab3b*, respectively. Values presented are the mean ± SEM (N = 2 groups: SAMP8 Control, n = 3, and SAMP8 T2 (5 mg/kg), n = 2). Student’s t-test analysis was performed: *p < 0.05. **j.** Gene expression of *Gpx4* and **k.** *Fsp1*. **l.** Levels of the GSG:GSSG ratio, **m.** ROS, **n.** Fe^2+^, and **o.** MDA in SAMP8 control and T2-treated (5 mg/kg) samples. Values presented are the mean ± SEM (N = 2 groups: SAMP8 Control, n = 7-5, and SAMP8 T2 (5 mg/kg), n = 5-4). Statistical tests: Student’s t-test: *p < 0.05; **p < 0.01; ***p < 0.001.

Additionally, enriched GO terms such as “translation at synapse” and “translation at postsynapse” suggest the reactivation of localized protein synthesis at dendritic sites, an essential process for memory consolidation (Figure 6e). From the GO downregulated pathways, the trend suggests a shift away from stress and immune-related gene expression. These terms indicate transcriptional silencing of genes involved in chronic immune surveillance, potentially tied to microglial and astrocytic reactivity (Figure 6e). Furthermore, the genes that T2 upregulates are linked to neuron cell types, particularly inhibitory neurons (Figure 6f).

Finally, we analyzed whether our scRNA-seq list is correlated with the list of genes that show ATAC-seq enrichment at distal regions. We identified 28 genes that were present in both lists (Supplementary Table 9). The gene number is too low for a gene set enrichment analysis; however, many genes are correlated with proteasome function and RNA processing, and are likely transcriptional regulators.

Although some top synaptic KEGG pathways were not captured in the most significantly enriched set, trends related to neurotransmitter release and neurotrophic signaling were evident. More notably, several downregulated KEGG pathways reflected suppression of degenerative mechanisms such as ferroptosis (Figures 6e). In line with this, T2 treatment in SAMP8 mice resulted in a significant upregulation of *Gpx4* and *Fsp1*, two key ferroptosis suppressor genes that act through glutathione-dependent and-independent mechanisms, respectively (Figures 6j-k). Additionally, T2 restored the redox balance, as evidenced by a higher GSH:GSSG ratio and reduced levels of ROS and Fe²⁺ compared to the Control group (Figures 6l-n). However, no differences were observed in the MDA levels between both groups (Figure 6o).

Overall, our results indicate that at the molecular level, T2 functions as a potent gene regulator, enhancing the expression of genes related to neuronal function while decreasing the number of microglial cells. These results suggest a dual role for T2, which can act by promoting both neural growth and survival, while also reducing the immune response characteristic of AD and aging.

### Global proteome remodeling in response to T2 uncovers mechanisms of neuroprotection

To investigate the molecular basis of T2’s neuroprotective effects, we performed unbiased proteomic profiling in brains from aged SAMP8 mice. Principal component analysis (PCA) revealed clear segregation between treated and control groups, indicating substantial proteome remodeling following treatment (Figure 7a). Differential expression analysis identified numerous proteins that were significantly upregulated or downregulated in T2-treated animals (Figure 7b). Hierarchical clustering confirmed consistent expression shifts across replicates (Figure 7c).

**Figure 7.**
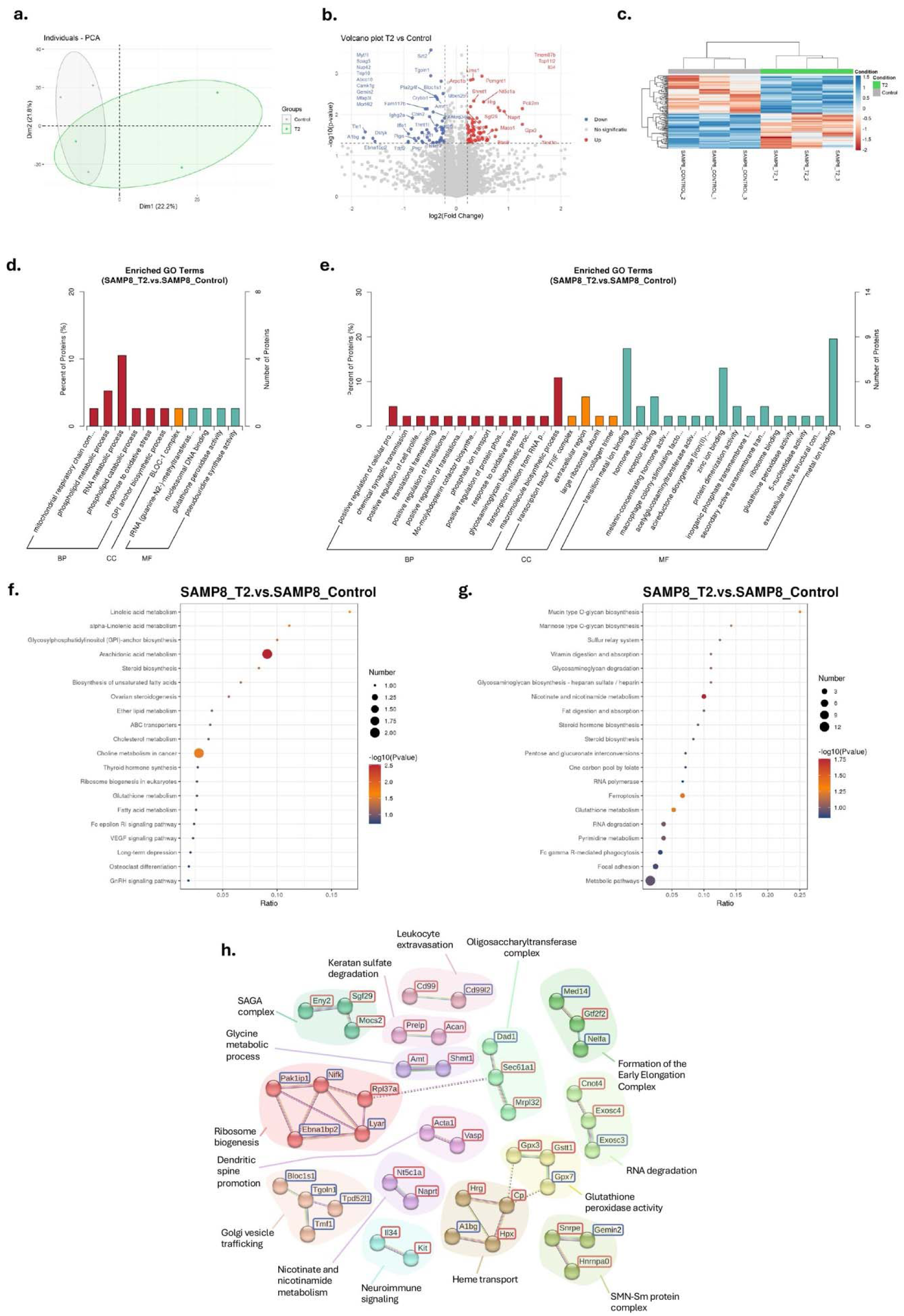
Proteomic overview of treatment effects in aged SAMP8 mouse brains. a. Principal component analysis (PCA) plot of all identified proteins, illustrating the overall separation between SAMP8 (grey ellipse) and the treated group (green ellipse). **b.** Volcano plots displaying differentially expressed proteins between treated and control SAMP8 mice. Statistically significant upregulated (red) and downregulated (blue) proteins are highlighted based on adjusted p-value and fold-change thresholds. **c.** Heatmap of top differentially expressed proteins, clustered by Euclidean distance across samples. **d-e**. Gene Ontology (GO) enrichment analysis of downregulated (D) and upregulated (E) proteins, grouped into biological processes (BP), cellular components (CC), and molecular functions (MF). **f-g.** KEGG pathway enrichment analysis for downregulated (F) and upregulated (G) proteins. Dot size represents the number of proteins associated with each pathway, and color indicates statistical significance (–log10(p-value)). **h.** String protein–protein interaction network of differentially expressed proteins in SAMP8 mice treated with T2. Only proteins with a minimum interaction score ≥ 0.5 were included. Clustering was performed using the Markov Cluster Algorithm (MCL) with an inflation parameter of 2.0, and functionally enriched groups were annotated based on STRING biological processes. Red-framed nodes indicate proteins upregulated, and blue-framed nodes indicate downregulated proteins in treated animals relative to untreated controls. Functional clusters include: ribosome biogenesis, Golgi vesicle trafficking, RNA degradation, glutathione peroxidase activity, immune-regenerative signaling, epigenetic regulation (SAGA complex), and dendritic spine remodeling, among others. *Clustering performed using the MCL algorithm (Enright et al., 2002)*.

To understand the functional impact of these changes, we performed a Gene Ontology (GO) enrichment analysis. Downregulated proteins were significantly associated with RNA metabolic processes (Figure 7d), whereas upregulated proteins were associated with metal ion binding and macromolecular biosynthetic processes (Figure 7e). Further analysis of KEGG pathways revealed that arachidonic acid metabolism was the most significantly downregulated pathway, while nicotinate/nicotinamide metabolism, glutathione metabolism, and ferroptosis were among the most upregulated (Figures 7f-g).

The suppression of arachidonic acid metabolism is of particular interest, as this pathway is implicated in neuroinflammation and synaptic degeneration in both humans and animal models of AD. Its downregulation may reflect reduced production of pro-inflammatory eicosanoids, supporting an anti-inflammatory role for T2.^26^

Conversely, the enrichment of ferroptosis-related proteins suggests that T2 activates antioxidant defense mechanisms against iron-induced neurotoxicity. This is relevant since iron accumulation and ferroptotic cell death are increasingly recognized as drivers of neurodegeneration in AD.^27^ Upregulated proteins included hemopexin (Hpx) and ceruloplasmin (Cp), both involved in iron and heme detoxification.^28,29^

Additionally, antioxidant enzymes such as GPX3 and GSTT1 were elevated. GPX3 acts as a key extracellular ROS scavenger, while GSTT1 catalyzes the conjugation of glutathione to electrophilic compounds, supporting the notion that T2 enhances the brain’s redox defense and mitigates ferroptotic stress.^30^

A protein–protein interaction (PPI) network, constructed using the STRING database, grouped the differentially expressed proteins into functional modules (Figure 7h). This revealed downregulated clusters related to ribosome biogenesis and Golgi vesicle trafficking, and upregulated clusters associated with glutathione peroxidase activity, heme transport, epigenetic regulation, and neuroimmune signaling.

Among the upregulated factors, interleukin-34 (IL-34) was of particular interest. IL-34 has been shown to modulate microglial activity, induce insulin-degrading enzyme (IDE), and upregulate heme oxygenase-1 (HO-1), thereby promoting Aβ clearance and reducing oxidative damage without inducing neurotoxicity.^31^

Importantly, proteins involved in ribosome biogenesis (e.g., Ebna1bp2, Pak1ip1, Rpl37a) and vesicle trafficking (e.g., Bloc1s1, Tgoln1, Tpd52l1) were consistently downregulated, suggesting suppression of the senescence-associated secretory phenotype (SASP), which contributes to chronic neuroinflammation in aging brains.^32^

T2 also upregulated SAGA complex members such as Eny2 and Sgf29, which are known to modulate gene expression through histone acetylation and deubiquitylation and are essential for transcriptional plasticity.^33^

Interestingly, Cd99l2, a gene implicated in leukocyte trafficking into the CNS, was downregulated. This is consistent with reduced neuroinflammatory signaling. Additionally, Cd99l2 has recently been identified as a negative regulator of neuronal activation via its role as a synaptic adhesion molecule,^34,35^ suggesting that its suppression may also support neuroadaptive plasticity.

Finally, key proteins involved in dendritic spine remodeling, including VASP and ACTA1, were significantly upregulated. VASP regulates actin polymerization in dendritic spines, promoting their growth and stabilization.^36^ The concurrent upregulation of ACTA1, a structural actin isoform, further supports the idea that T2 enhances synaptic structure and function, potentially contributing to improved cognitive performance in aged animals.

## DISCUSSION

AD presents a complex pathological landscape, including amyloid buildup, tau pathology, synaptic dysfunction, and neuroinflammation, which can overwhelm therapies that target only single mechanisms.^37^ Developing compound T2, a dual G9a/GSK-3β inhibitor, signifies a shift from traditional single-target treatments that have often failed to produce meaningful clinical results for AD patients. By simultaneously addressing epigenetic dysregulation and kinase hyperactivation—two key drivers of neurodegeneration—this polypharmacological approach better tackles AD’s multifactorial nature than selective inhibitors alone.^4,6,38,39^ Our findings show that inhibiting G9a and GSK-3β together results in strong improvements at molecular, synaptic, and behavioral levels, such as normalizing tau phosphorylation and restoring dendritic complexity, which are often missing in single-target therapies. Additionally, genome-wide and multi-omics studies demonstrate that dual inhibition offers significantly better neuroprotection than targeting either enzyme alone. This supports the growing evidence that network redundancy and compensatory signaling pathways can weaken the effectiveness of single-target treatments.

From a chemical standpoint, the optimized tetrahydropyrimidine scaffold overcame key limitations of previous frameworks by achieving balanced potency at both targets and a favorable BBB profile. This is a critical advance, as many promising GSK-3β or G9a inhibitors fail to reach adequate brain exposure or cause dose-limiting toxicities with prolonged administration.^24,38,39,41^ For example, studies using excessive GSK-3β inhibition with agents like SAR502250 or SB216763 have documented concerning adverse effects, including neurotoxicity during prolonged treatment.^38,39^ Our balanced dual mechanism successfully achieves disease modification while avoiding the safety concerns and efficacy limitations observed with single-target or poorly balanced dual agents. Extensive molecular dynamics simulations and computational analyses confirm T2’s unique and stable engagement at both target sites, a characteristic that has not been well-documented for previously published dual ligands. Furthermore, pharmacokinetic and safety data indicate that T2 achieves sustained central concentrations with minimal off-target activity and no detectable genotoxicity or mutagenicity at therapeutic doses, supporting its translational potential for chronic CNS use.

Behaviorally, T2 caused notable improvements in cognitive performance and social interaction in SAMP8 mice, a model of LOAD, exceeding the improvements seen with selective GSK-3β or G9a inhibitors like SAR502250, SB216763, UNC0642, or Tideglusib.^3,8,38,39^ This greater effectiveness was also reflected in enhanced synaptic plasticity and AMPAR-mediated transmission, along with increased expression of neurotrophic and immediate-early genes (*Bdnf*, *NT3*, *Arc*, *Comt*) and a decrease in pro-inflammatory cytokines (*Il-1*β). Overall, these results suggest a mechanism where T2 simultaneously encourages neural growth and survival while reducing neuroimmune activation.

A key aspect of our work is the integration of multi-omics analyses to define the range of T2’s effects. It showed that dual inhibitor therapy uniquely remodels the epigenome, transcriptome, and cellular networks with a level of molecular reprogramming that exceeds single-target approaches. ChIP-seq and ATAC-seq revealed a widespread reduction of H3K9me2 and a significant increase in chromatin accessibility at distal regulatory elements enriched for motifs such as FOSL2, ZNF281, and NFIA. This indicates that T2’s neuroprotective effect largely comes from derepression of genes involved in neuronal projection and membrane potential located at enhancer regions, complementing direct effects at promoters. At the same time, single-cell RNA-seq analysis showed an increased proportion of neurons and a decreased proportion of microglia in treated animals, with transcriptional signatures pointing to axon guidance, synaptic vesicle transport, and localized translation at dendritic sites — processes crucial for memory consolidation.^18,27,30^

A key innovative feature of our work is demonstrating that T2 effectively inhibits ferroptosis and related cell death pathways, as shown by proteomic and redox analyses. This layered neuroprotective mechanism sets our approach apart from most current treatments, which often overlook iron-dependent neurotoxicity and oxidative stress. Our proteomic data support this, showing a coordinated decrease in pathways related to neuroinflammation—such as arachidonic acid metabolism and SASP components^26^—and an increase in ferroptosis inhibitors and antioxidant enzymes like Hpx, Cp, GPX3, GSTT1,^28–30^, along with IL-34 and SAGA complex proteins.^31,33^ These changes suggest T2 promotes iron and heme detoxification, reduces oxidative stress, and restores transcriptional flexibility, countering multiple degenerative processes. This comprehensive molecular and functional evidence sets a new standard for disease-modification strategies, especially considering the growing recognition of iron-induced neurotoxicity and ferroptosis as key factors in AD development.27,30 Importantly, our identification and consistent modulation of plasma tau phosphorylation species (pT181, pT217, pT231), well-recognized clinical and pharmacodynamic biomarkers,^15,17,21,22,42^ represent a significant breakthrough for clinical translation. In previous therapeutic efforts, such biomarkers were often weakly linked to treatment response or absent from study endpoints, which greatly hindered progress toward precision medicine. Our dual-target intervention reliably modulates these biomarkers in direct relation to functional improvements, strongly supporting their use in clinical development and patient monitoring.

Taken together, our findings show that dual inhibition of G9a and GSK-3β by the small molecule T2 offers an innovative and potentially transformative therapeutic approach for AD. T2 not only demonstrated remarkable effectiveness in reversing cognitive and morphological deficits in the SAMP8 model but also positively influenced key neurodegeneration biomarkers, leading to transcriptomic and proteomic reprogramming toward neuroprotection and synaptic plasticity, and activating essential antioxidant mechanisms. These effects, combined with a promising safety profile and strong ability to penetrate the central nervous system, position T2 as a highly promising candidate for developing disease-modifying therapies. Compared to the growing evidence supporting single-target and earlier multi-target strategies, our data highlight that true disease modification in AD requires simultaneous, rational targeting of both epigenetic and kinase pathways. The balanced dual G9a/GSK-3β inhibitor presented here achieves multidimensional efficacy, safety, and mechanistic validation where single or simpler multi-target approaches have failed, reinforcing the idea that the future of AD drug development lies in precise, multi-target pharmacological design driven by molecular insights to address the disease’s inherent complexity and overcome the limitations of less comprehensive strategies.

## ASSOCIATED CONTENT

Supporting information.

The Supporting Information is available free of charge at https//pubs.acs.org/doi/

Experimental details, characterization data, and spectra for the compounds of the synthesized compounds (PDF).

## ACKNOWLEDGEMENTS

This study was supported by the Ministerio de Economía, Industria Economía, Industria y Competitividad (Agencia Estatal de Investigación, AEI) and European Union NextGenerationEU/PRTR (PID2022-139016OA-I00, PDC2022-133441-I00,to CGF and MP), MICIU/AEI/10.13039/501100011033 and FEDER, UE and PDC2022-133441-I00 MICIU/AEI /10.13039/501100011033 Europea Next GenerationEU/ PRTR), Generalitat de Catalunya (2021 SGR 00357). This work was supported by Grant PID2022-1380790B-I00 funded by MCIN/AEI/10.13039/501100011033 and by ERDF A way of making Europe (to C.E.), Grant PDC2022-133441-I00 funded by MCIN/AEI/10.13039/501100011033 and by the European Union NextGenerationEU/PRTR (to C.E.), and Grant 2021 SGR 00357 funded by Generalitat de Catalunya (to C.E.). This study was co-financed by Secretaria d’Universitats i Recerca del Departament d’Empresa i Coneixement de la Generalitat de Catalunya 2023 (Product 0092; Llavor 005 and Llavor 007 to CGF). SS and BSC are grateful to Council for Scientific and Industrial Research, India, for Senior Research Fellowship (Grant No 09/1131(0026)/2019-EMR-I and 09/1131(0004)/2016-EMR-I) and Central University of Rajasthan for providing computational facilities. ALM and JB are grateful to Xunta de Galicia (ED431C 2018/21 and ED431G 2019/02) and European Regional Development Fund (ERDF) in the frame of the Recovery Assistance for Cohesion and the Territories of Europe (REACT-EU) funds. As mentioned in the respective figure legends, Figures 1a–b and 4a–b reproduce data previously reported by our group (Bellver-Sanchis et al., Aging Dis. 2024;15(1):311), included here to provide a cohesive narrative within the present manuscript.

## ABBREVIATIONS

GSK-3β: glycogen synthase kinase 3 beta
AD: Alzheimer’s disease
Aβ: beta-amyloid
C. elegans: Caenorhabditis elegans
MNT: micronucleus test
MTD: maximal tolerated dose
SAMP8: senescence-accelerated mouse-prone 8
LOAD: late-onset Alzheimer’s disease
PoC: proof-of-concept
NORT: Novel Object Recognition Test
OLT: Object Location Test
TCT: Three-Chamber Test
mEPSCs: miniature excitatory postsynaptic currents.

---

**Figure S1.**
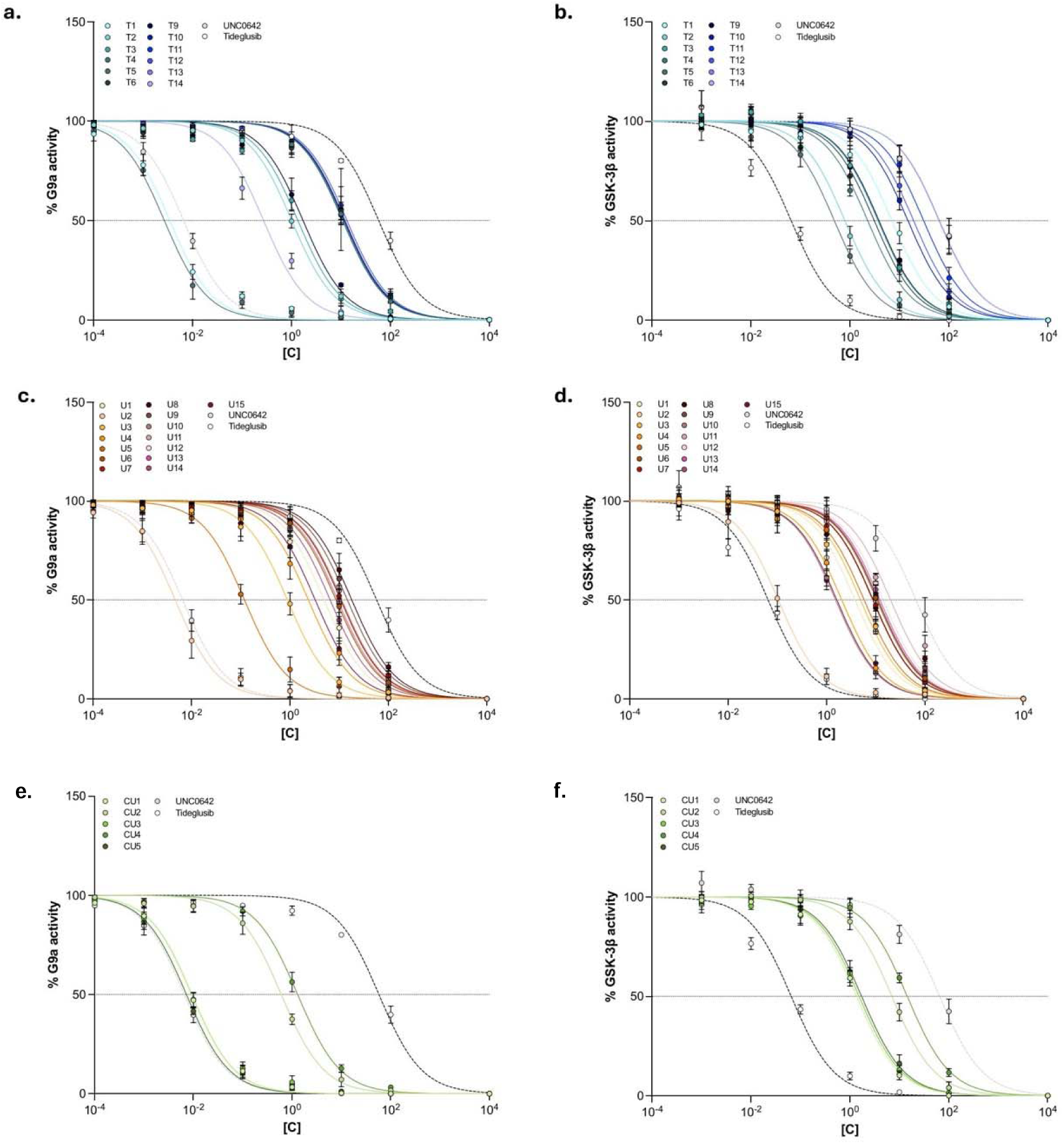
Representative IC_50_ graph against (**a,c** and **e**) G9a and (**b,d,** and **f**) GSK-3β of the Series 1 and Series 2 compounds.

**Figure S2.**
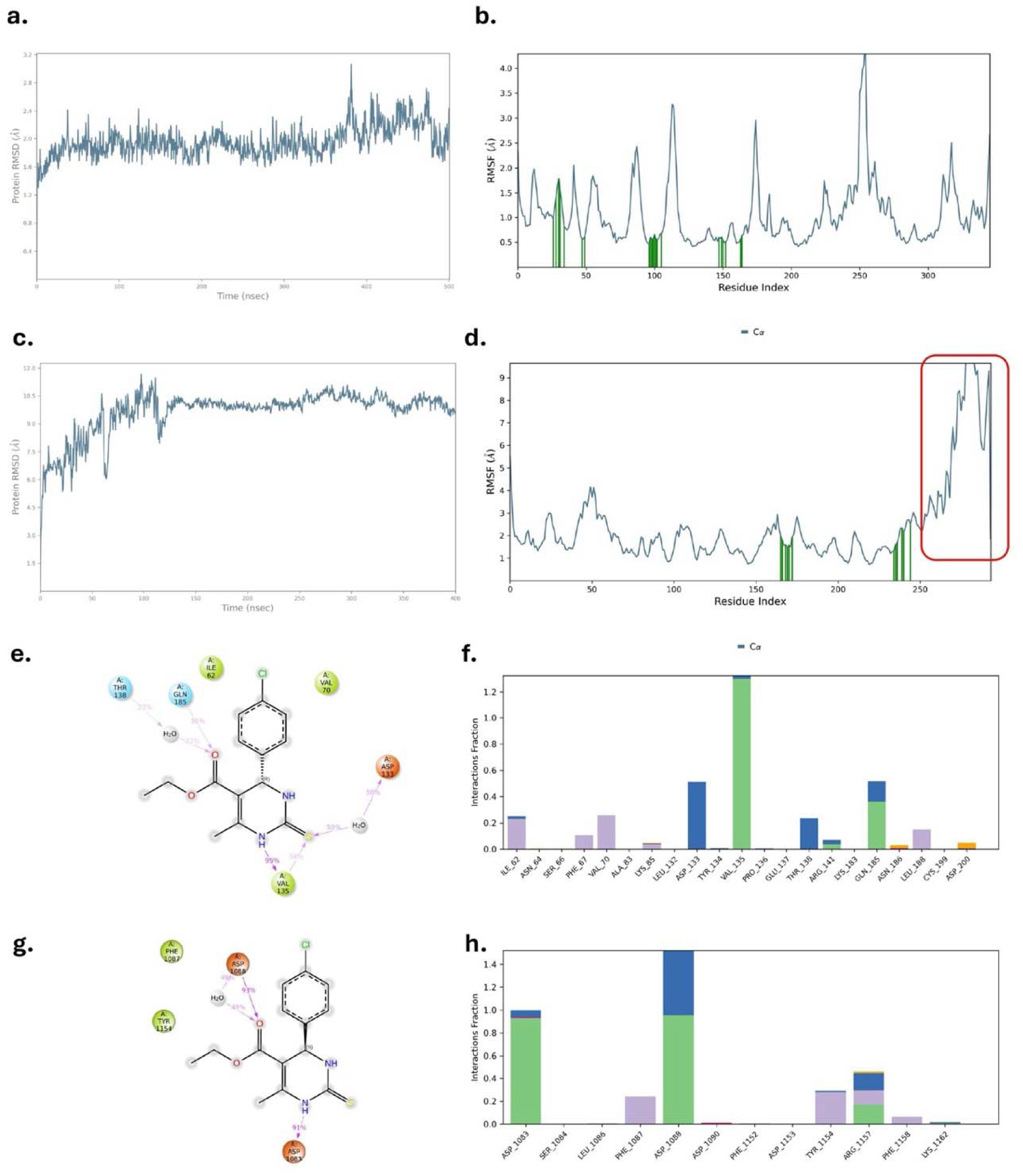
a.RMSD & **b.** RMSF of GSK3β-T2 complex remain stable around 2Å through out simulation, **c**. RMSD of G9a-T2 remain stable around 10Å through out simulation, reason of this higher RMSD is RMSF graph **d.** Higher fluctuation marked in red square of solvent exposed C terminal of G9a, apart from that all residues of G9a are stable around 2Å. **e.-f.** 2D interaction diagram and protein-ligand interaction histogram of GSK3β-T2 show strong hydrogen bond interaction with Val135 of kinase hinge domain, **g.-h.** 2D interaction diagram and protein-ligand interaction histogram of G9a-T2 show strong interaction with Asp1083 and Asp1088 of substrate binding region.

**Figure S3.**
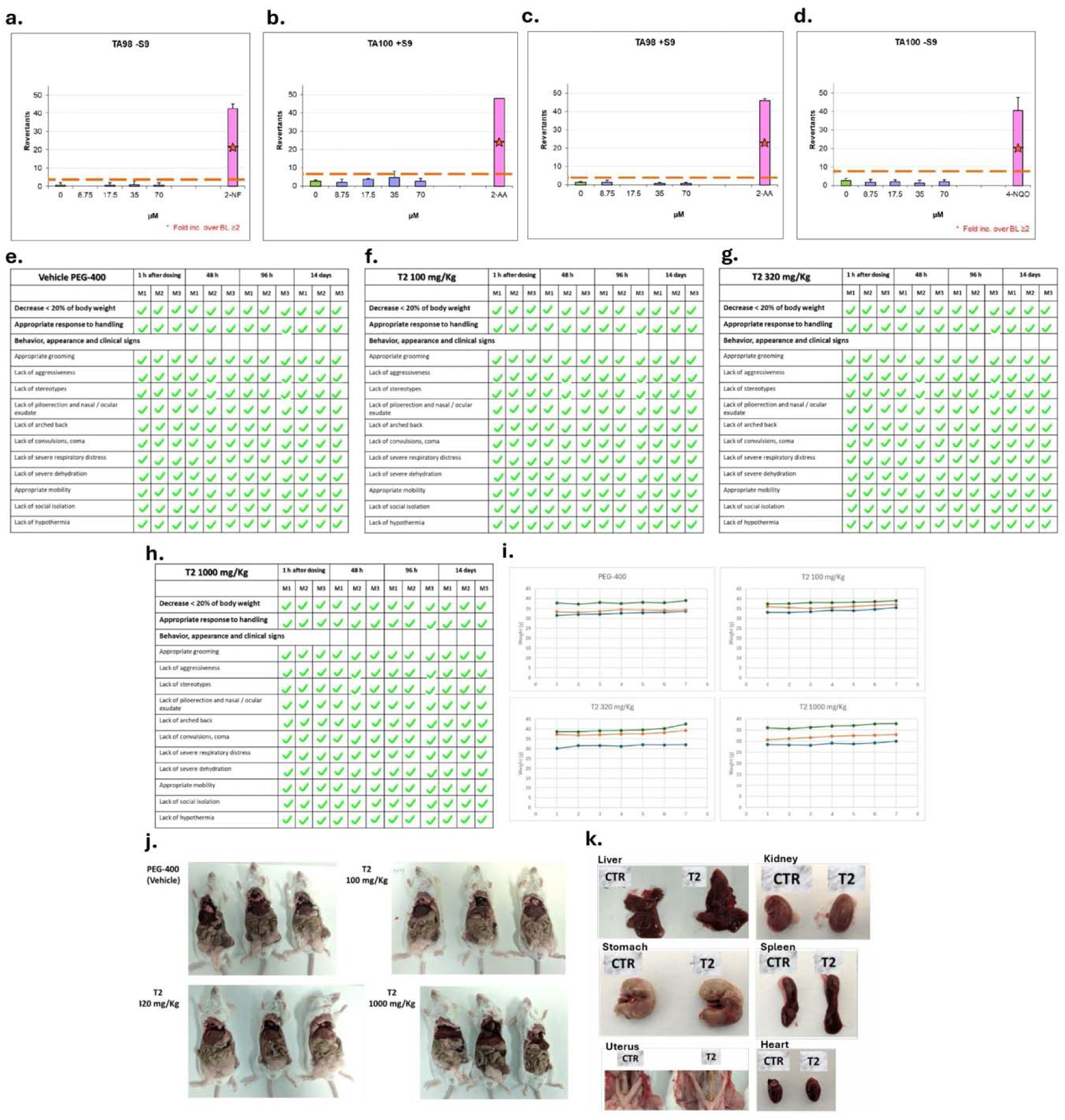
a. and **b.** Number of positive mutated wells of T2 using *Salmonella typhimurium (S. typhimurium)* TA98**.c.** and **d.** Number of positive mutated wells of T2 using S. typhimurium TA98 and **E.** TA100 with S9 activation (Left column). **e.-h.** The results of supervision protocol observation are summarized in the figure. **g.** Weights of mice during 14 days of treatment with T2. The weight (g) of the mice is represented (every 2 days). **i.** Mice were sacrificed 14 days after treatment and gross necropsy were performed, photographs of necropsy are collected. **j.** Mice treated with 1000 mg/Kg T2 compound after sacrificing compared to control mice: organs comparison.

**Figure S4.**
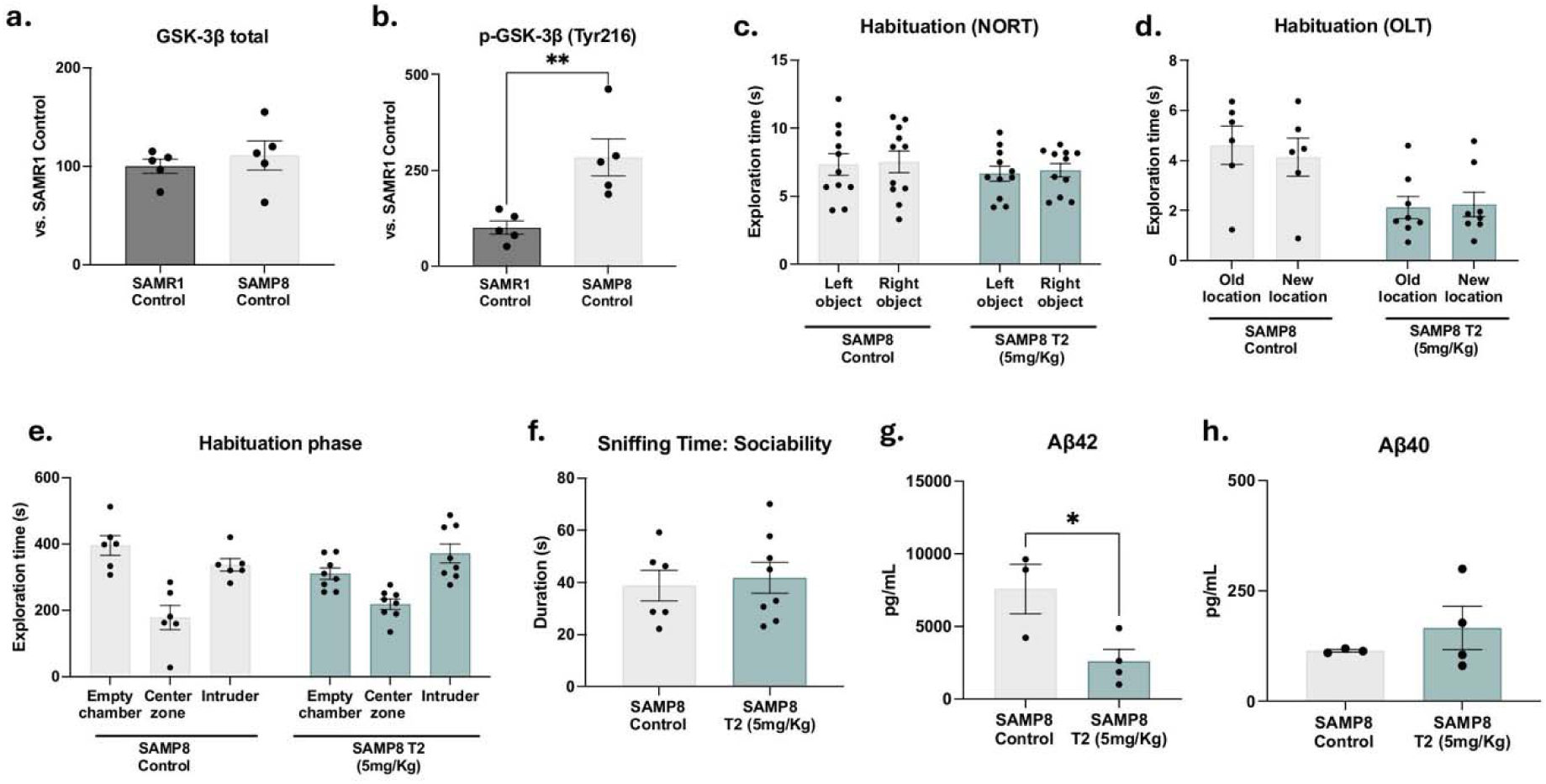
a. Quantification of GSK-3β and **b.** p-GSK-3β (Tyr216) protein levels in SAMP8 compared to SAMR1. Values presented are the mean ± SEM (SAMR1 Control, n = 5-6, and SAMP8, n = 5-6). Statistical analysis: Student’s t-test analysis; *p < 0.05; **p<0.01. **c.** NORT, **d.** OLT and **e.** TCT habituation. **f**. Sniffing time evaluated during TCT. **g**. Levels of Aβ42 and **h**. Aβ40. Values presented as mean ± SEM; (SAMP8 Control n = 10, and SAMP8 T2 (5mg/Kg) n = 10); Statistics analysis: Student’s t-test analysis; *p<0.05.

**Table S1.**
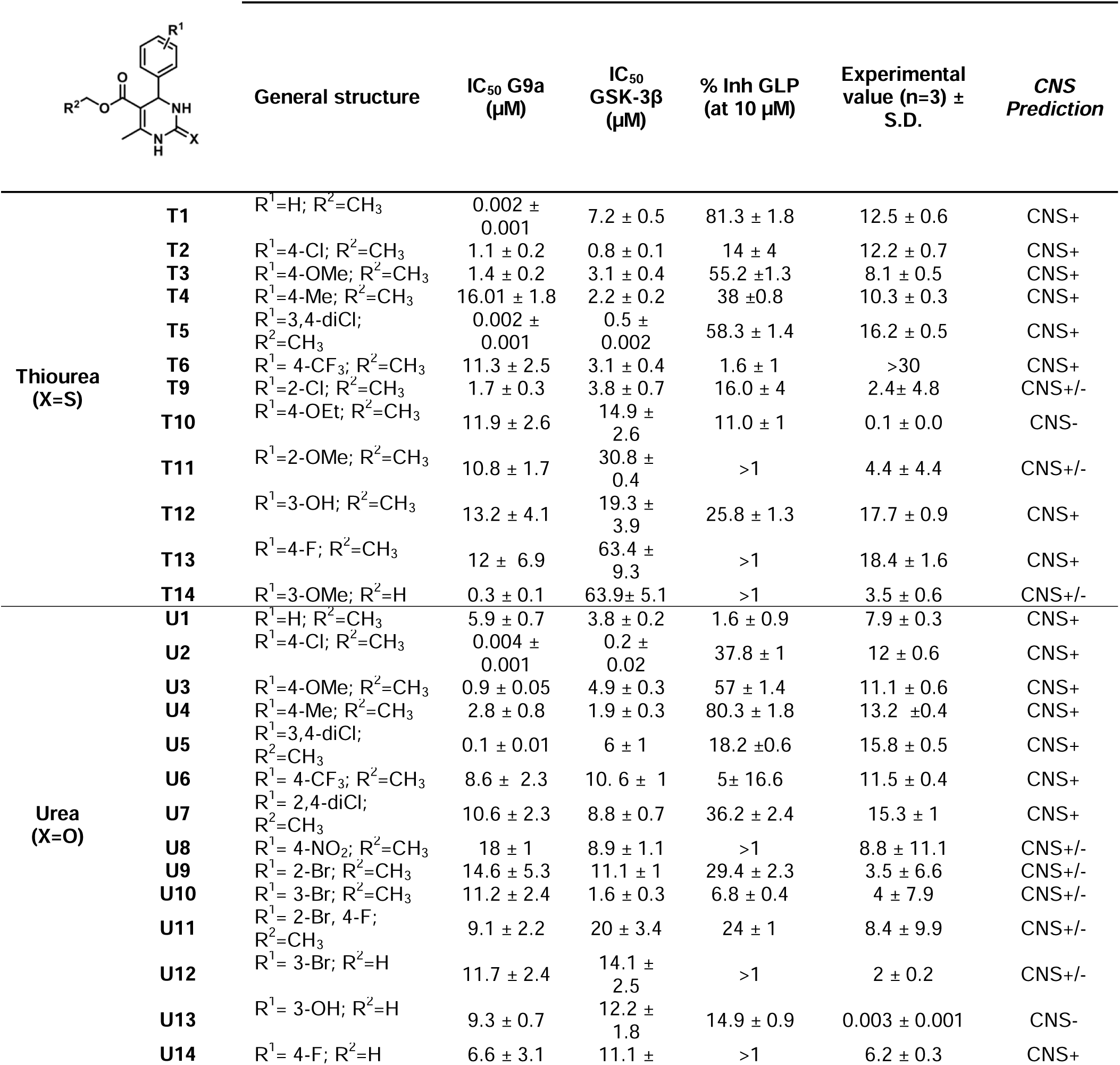
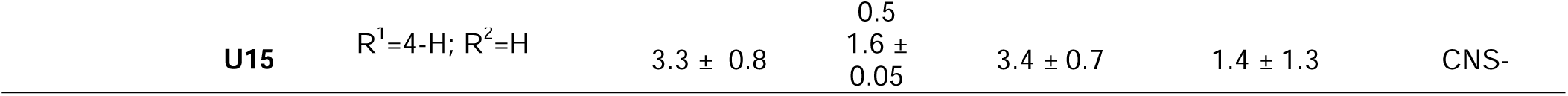
Designed and synthesized compounds from Series 1. This table contain IC_50_ for both targets (G9a and GSK-3β, µM), percentage of GLP inhibition at 10 µM and Permeability results (*Pe* 10-6 cm s-1) from the PAMPA-BBB assay and BBB permeation prediction. For the reference compounds used for the PAMPA-BBB assay see Table S8.

| | | General structure | IC<sub>50</sub> G9a<br>(μM) | IC<sub>50</sub> GSK-3β<br>(μM) | % Inh GLP<br>(at 10 μM) | Experimental<br>value (n=3) ±<br>S.D. | CNS<br>Prediction |
| --- | --- | --- | --- | --- | --- | --- | --- |
| Thiourea<br>(X=S) | T1 | R<sup>1</sup>=H; R<sup>2</sup>=CH<sub>3</sub> | 0.002 ±<br>0.001 | 7.2 ± 0.5 | 81.3 ± 1.8 | 12.5 ± 0.6 | CNS+ |
| | T2 | R<sup>1</sup>=4-Cl; R<sup>2</sup>=CH<sub>3</sub> | 1.1 ± 0.2 | 0.8 ± 0.1 | 14 ± 4 | 12.2 ± 0.7 | CNS+ |
| | T3 | R<sup>1</sup>=4-OMe; R<sup>2</sup>=CH<sub>3</sub> | 1.4 ± 0.2 | 3.1 ± 0.4 | 55.2 ± 1.3 | 8.1 ± 0.5 | CNS+ |
| | T4 | R<sup>1</sup>=4-Me; R<sup>2</sup>=CH<sub>3</sub> | 16.01 ± 1.8 | 2.2 ± 0.2 | 38 ± 0.8 | 10.3 ± 0.3 | CNS+ |
| | T5 | R<sup>1</sup>=3,4-diCl;<br>R<sup>2</sup>=CH<sub>3</sub> | 0.002 ±<br>0.001 | 0.5 ±<br>0.002 | 58.3 ± 1.4 | 16.2 ± 0.5 | CNS+ |
| | T6 | R<sup>1</sup>= 4-CF<sub>3</sub>; R<sup>2</sup>=CH<sub>3</sub> | 11.3 ± 2.5 | 3.1 ± 0.4 | 1.6 ± 1 | >30 | CNS+ |
| | T9 | R<sup>1</sup>=2-Cl; R<sup>2</sup>=CH<sub>3</sub> | 1.7 ± 0.3 | 3.8 ± 0.7 | 16.0 ± 4 | 2.4 ± 4.8 | CNS+/- |
| | T10 | R<sup>1</sup>=4-OEt; R<sup>2</sup>=CH<sub>3</sub> | 11.9 ± 2.6 | 14.9 ±<br>2.6 | 11.0 ± 1 | 0.1 ± 0.0 | CNS- |
| | T11 | R<sup>1</sup>=2-OMe; R<sup>2</sup>=CH<sub>3</sub> | 10.8 ± 1.7 | 30.8 ±<br>0.4 | >1 | 4.4 ± 4.4 | CNS+/- |
| | T12 | R<sup>1</sup>=3-OH; R<sup>2</sup>=CH<sub>3</sub> | 13.2 ± 4.1 | 19.3 ±<br>3.9 | 25.8 ± 1.3 | 17.7 ± 0.9 | CNS+ |
| | T13 | R<sup>1</sup>=4-F; R<sup>2</sup>=CH<sub>3</sub> | 12 ± 6.9 | 63.4 ±<br>9.3 | >1 | 18.4 ± 1.6 | CNS+ |
| | T14 | R<sup>1</sup>=3-OMe; R<sup>2</sup>=H | 0.3 ± 0.1 | 63.9 ± 5.1 | >1 | 3.5 ± 0.6 | CNS+/- |
| Urea<br>(X=O) | U1 | R<sup>1</sup>=H; R<sup>2</sup>=CH<sub>3</sub> | 5.9 ± 0.7 | 3.8 ± 0.2 | 1.6 ± 0.9 | 7.9 ± 0.3 | CNS+ |
| | U2 | R<sup>1</sup>=4-Cl; R<sup>2</sup>=CH<sub>3</sub> | 0.004 ±<br>0.001 | 0.2 ±<br>0.02 | 37.8 ± 1 | 12 ± 0.6 | CNS+ |
| | U3 | R<sup>1</sup>=4-OMe; R<sup>2</sup>=CH<sub>3</sub> | 0.9 ± 0.05 | 4.9 ± 0.3 | 57 ± 1.4 | 11.1 ± 0.6 | CNS+ |
| | U4 | R<sup>1</sup>=4-Me; R<sup>2</sup>=CH<sub>3</sub> | 2.8 ± 0.8 | 1.9 ± 0.3 | 80.3 ± 1.8 | 13.2 ± 0.4 | CNS+ |
| | U5 | R<sup>1</sup>=3,4-diCl;<br>R<sup>2</sup>=CH<sub>3</sub> | 0.1 ± 0.01 | 6 ± 1 | 18.2 ± 0.6 | 15.8 ± 0.5 | CNS+ |
| | U6 | R<sup>1</sup>= 4-CF<sub>3</sub>; R<sup>2</sup>=CH<sub>3</sub> | 8.6 ± 2.3 | 10.6 ± 1 | 5 ± 16.6 | 11.5 ± 0.4 | CNS+ |
| | U7 | R<sup>1</sup>= 2,4-diCl;<br>R<sup>2</sup>=CH<sub>3</sub> | 10.6 ± 2.3 | 8.8 ± 0.7 | 36.2 ± 2.4 | 15.3 ± 1 | CNS+ |
| | U8 | R<sup>1</sup>= 4-NO<sub>2</sub>; R<sup>2</sup>=CH<sub>3</sub> | 18 ± 1 | 8.9 ± 1.1 | >1 | 8.8 ± 11.1 | CNS+/- |
| | U9 | R<sup>1</sup>= 2-Br; R<sup>2</sup>=CH<sub>3</sub> | 14.6 ± 5.3 | 11.1 ± 1 | 29.4 ± 2.3 | 3.5 ± 6.6 | CNS+/- |
| | U10 | R<sup>1</sup>= 3-Br; R<sup>2</sup>=CH<sub>3</sub> | 11.2 ± 2.4 | 1.6 ± 0.3 | 6.8 ± 0.4 | 4 ± 7.9 | CNS+/- |
| | U11 | R<sup>1</sup>= 2-Br, 4-F;<br>R<sup>2</sup>=CH<sub>3</sub> | 9.1 ± 2.2 | 20 ± 3.4 | 24 ± 1 | 8.4 ± 9.9 | CNS+/- |
| | U12 | R<sup>1</sup>= 3-Br; R<sup>2</sup>=H | 11.7 ± 2.4 | 14.1 ±<br>2.5 | >1 | 2 ± 0.2 | CNS+/- |
| | U13 | R<sup>1</sup>= 3-OH; R<sup>2</sup>=H | 9.3 ± 0.7 | 12.2 ±<br>1.8 | 14.9 ± 0.9 | 0.003 ± 0.001 | CNS- |
| | U14 | R<sup>1</sup>= 4-F; R<sup>2</sup>=H | 6.6 ± 3.1 | 11.1 ± | >1 | 6.2 ± 0.3 | CNS+ |

|  |  |  |  |  |  |  |
| --- | --- | --- | --- | --- | --- | --- |
| **U15** | R<sup>1</sup>=4-H; R<sup>2</sup>=H | 3.3 ± 0.8 | 0.5<br>1.6 ±<br>0.05 | 3.4 ± 0.7 | 1.4 ± 1.3 | CNS- |

**Table S2.**
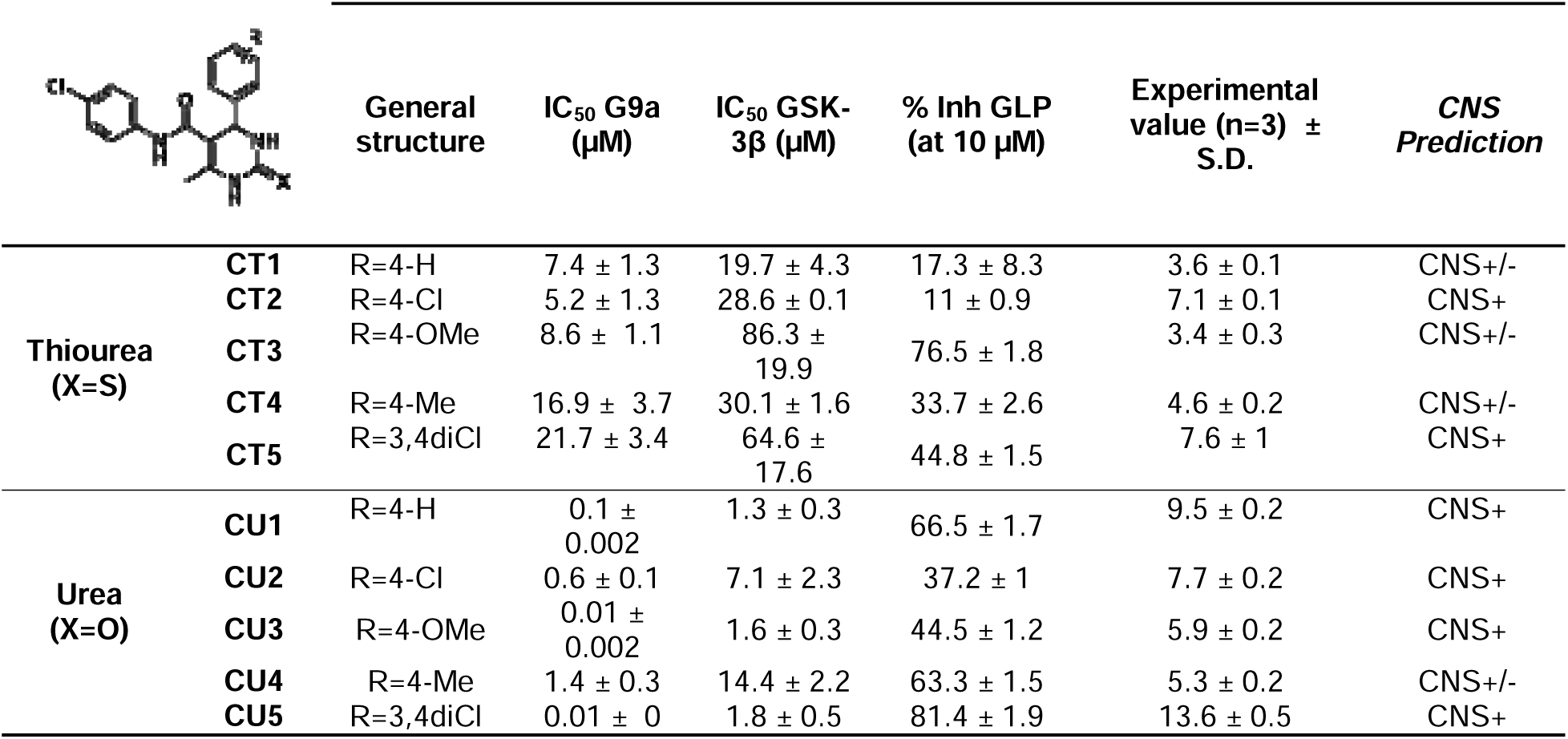
Designed and synthesized compounds from Series 2. This table contain IC_50_ for both targets (G9a and GSK-3β, µM), percentage of GLP inhibition at 10 µM and Permeability results (*Pe* 10-6 cm s-1) from the PAMPA-BBB assay and BBB permeation prediction.

|  |  | General structure | IC<sub>50</sub> G9a (μM) | IC<sub>50</sub> GSK-3β (μM) | % Inh GLP (at 10 μM) | Experimental value (n=3) ± S.D. | CNS Prediction |
| --- | --- | --- | --- | --- | --- | --- | --- |
| Thiourea (X=S) | CT1 | R=4-H | 7.4 ± 1.3 | 19.7 ± 4.3 | 17.3 ± 8.3 | 3.6 ± 0.1 | CNS+/- |
|  | CT2 | R=4-Cl | 5.2 ± 1.3 | 28.6 ± 0.1 | 11 ± 0.9 | 7.1 ± 0.1 | CNS+ |
|  | CT3 | R=4-OMe | 8.6 ± 1.1 | 86.3 ± 19.9 | 76.5 ± 1.8 | 3.4 ± 0.3 | CNS+/- |
|  | CT4 | R=4-Me | 16.9 ± 3.7 | 30.1 ± 1.6 | 33.7 ± 2.6 | 4.6 ± 0.2 | CNS+/- |
|  | CT5 | R=3,4diCl | 21.7 ± 3.4 | 64.6 ± 17.6 | 44.8 ± 1.5 | 7.6 ± 1 | CNS+ |
| Urea (X=O) | CU1 | R=4-H | 0.1 ± 0.002 | 1.3 ± 0.3 | 66.5 ± 1.7 | 9.5 ± 0.2 | CNS+ |
|  | CU2 | R=4-Cl | 0.6 ± 0.1 | 7.1 ± 2.3 | 37.2 ± 1 | 7.7 ± 0.2 | CNS+ |
|  | CU3 | R=4-OMe | 0.01 ± 0.002 | 1.6 ± 0.3 | 44.5 ± 1.2 | 5.9 ± 0.2 | CNS+ |
|  | CU4 | R=4-Me | 1.4 ± 0.3 | 14.4 ± 2.2 | 63.3 ± 1.5 | 5.3 ± 0.2 | CNS+/- |
|  | CU5 | R=3,4diCl | 0.01 ± 0 | 1.8 ± 0.5 | 81.4 ± 1.9 | 13.6 ± 0.5 | CNS+ |

**Table S3.**
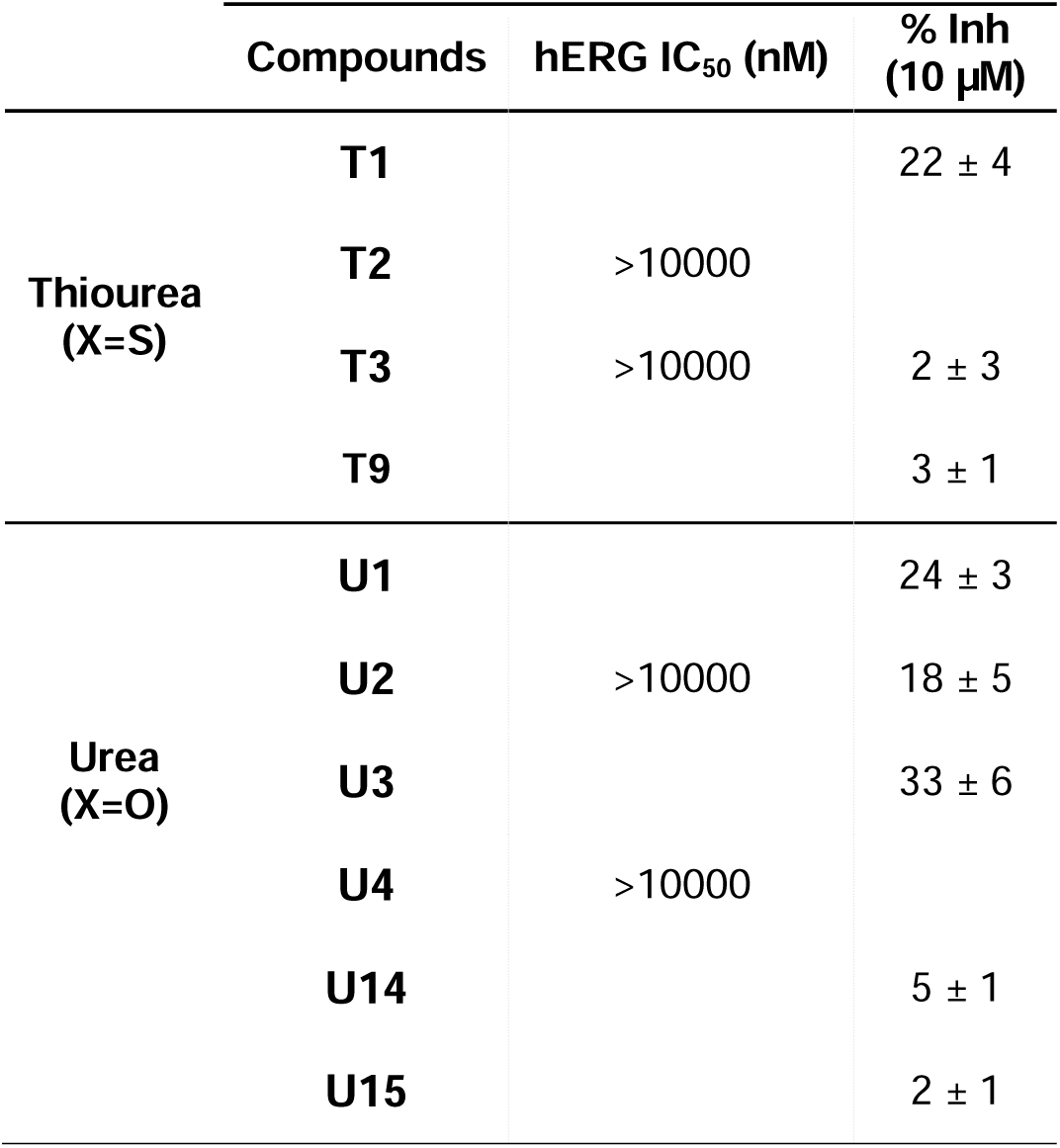
hERG IC_50_ and percentage of inhibition at 10 µM of selected compounds.

**Table S4.** Results of solubility (µM), and protein binding, and stability in human plasma for selected compounds from Series 1.

| | | Solubility<br>(μM) | Human plasma protein binding | | Stability in human plasma | | | |
| --- | --- | --- | --- | --- | --- | --- | --- | --- |
|  |  |  | Plasma protein binding (%) | Unbound fraction (%) | T (min) |  |  |  |
|  |  |  |  |  | 0 | 60 | 120 | 360 |
| Thiourea<br>(X=S) | T1 | 79 | 90.29 | 9.71 | 100 | 94.5 | 95.4 | 92.9 |
|  | T2 |  | 92.8 | 7.2 | 100 | 100.3 | 102.3 | 101.7 |
|  | T3 |  | 77.24 | 22.76 | 100 | 104.8 | 103.6 | 98.0 |
|  | T9 | > 100 | 96.21 | 3.79 | 100 | 94.1 | 94.0 | 64.0 |
| Urea | U1 |  | 78.89 | 21.11 | 100 | 94.8 | 108.3 | 109.9 |
| (X=O) | U2 |  | 96.83 | 3.17 | 100 | 94.9 | 84.7 | 95.6 |
|  | U3 |  |  |  |  |  |  |  |
|  | U4 |  | 92.74 | 7.26 | 100 | 104.9 | 95.5 | 103.7 |
|  | U14 | > 100 | 88.85 | 11.15 | 100 | 98.8 | 103.2 | 99.1 |
|  | U15 | > 100 | 77.51 | 22.49 | 100 | 96.6 | 87.3 | 87.9 |

**Table S5.** Results of microsomal stability and security panel at a recombinant human cytochrome P450 (CYP450) panel for selected compounds from Series 1.

|  |  | Microsomal stability |  |  | Security panel Recombinant human cytochrome P450 (CYP450) enzymes |  |  |  |
| --- | --- | --- | --- | --- | --- | --- | --- | --- |
|  |  | %Remanent (60min) | t1/2 (min) | Clint (μL/min* mg prot) | CYP1A2<br>% inh (10 μM, 1μM*) | CYP2C9<br>% inh (10 μM, 1μM*) | CYP2C19<br>% inh (10 μM, 1μM*)/IC<sub>50</sub> (nM)** | CYP2D6<br>% inh (10 μM, 1μM*)/IC<sub>50</sub> (nM)** |
| Thiourea (X=S) | T1 |  |  |  | 16.6 ± 4.3* | 12.8 ± 2* | -15.4 ± 12* | 15.7 ± 5.3* |
|  | T2 | 77.7 | 141.8 | 7.8 | 34 ± 1 | 62 ± 1 | 263.3** | 33 ± 1** |
|  | T3 |  |  |  | -13.3 ± 2* | 3.7 ± 1.1* | 11.7 ± 4.5* | -3.2 ± 0.2* |
|  | T9 | 82.4 | 221.2 | 3 | 12.9 ± 3.2 | 9.4 ± 4.9 | -11.6 ± 6.9 | 3 ± 10.1 |
| Urea (X=O) | U1 | 45.6 | 42.9 | 22.4 | -11.9 ± 1.7* | 6.4 ± 7.8* | 19.8 ± 0.3* | 5.3 ± 16.8* |
|  | U2 |  |  |  | -10.7 ± 1.8* | -0.7 ± 3.4* | -1.1 ± 3.9* | -7.6 ± 4.3* |
|  | U4 |  |  |  | 14.2 ± 7.6* | 1.5 ± 7.8* | -12.9 ± 7.8* | -12.8 ± 5* |
|  | U14 | 74.1 | 148.4 | 5.4 | -0.8 ± 7.8 | 4.3 ± 7.4 | 7.3 ± 20.3 | 7.7 ± 10.7 |
|  | U15 | 93.0 | 644.3 | 1.2 | 9.3 ± 6.6 | 4.5 ± 0.3 | -4.9 ± 7.2 | 4.7 ± 12.7 |

**Table S6.** Results of compound and positive control (CTR+) in MNT test. Compound is assayed at different concentrations and results are calculated in terms of number of binucleate cells, cytostasis percentage ± SD and percentage of micronuclei in binucleate cells ± SD.

| Sample | Concentration | No. binucleate cells | % Cytostasis | % MN |
| --- | --- | --- | --- | --- |
| T2 | 200 μM | 203 | 30.2 | 4.4 |
| T2 | 100 μM | 412 | 16.3 | 9.9 |
| T2 | 50 μM | 850 | 0 | 7.5 |
| T2 | 25 μM | 1298 | 0 | 7.5 |
| T2 | 12.5 μM | 1337 | 0 | 9.5 |
| CTR- |  | 1487 | - | 7.2 |
| CTR + (MitoC) | 1 μM | 411 | 37.7 | 28.9 |
| CTR + (MitoC) | 0.5 μM | 815 | 16.8 | 19.0 |

**Table S7.** a.Mutagenic activity of T2 using *S. typhimurium* TA98 and TA100 without S9 activation. **b.** Mutagenic activity of T2 using *S. typhimurium* TA98 and TA100 with S9 activation. Abbreviations: 2-NF: 2-nitrofluorene; 4-NQO: 4-nitroquinoline-N-oxide; 2-AA: 2-aminoanthracene.

| a. |  | TA98 - S9 |  |  | TA98 + S9 |  |  | Mutagenic Conc./ Cytotoxic effect |
| --- | --- | --- | --- | --- | --- | --- | --- | --- |
|  |  | Number positive wells |  |  | Number positive wells |  |  |  |
| Sample | Concentration (µM) | Replicate #1 | Replicate #2 | Replicate #3 | Replicate #1 | Replicate #2 | Replicate #3 |  |
| T2 | 0 | 0 | 2 | 0 | 1 | 2 | 1 |  |
|  | 8.75 | 0 | 0 | 0 | 0 | 2 | 2 |  |
|  | 17.5 | 0 | 2 | 0 | 0 | 0 | 0 |  |
|  | 35 | 0 | 0 | 3 | 1 | 1 | 0 |  |
|  | 70 | 0 | 2 | 0 | 1 | 0 | 1 |  |
| 2-NF | 2 µg/mL | 43 | 45 | 40 |  |  |  | Mutagenic Conc. |
| 2-AA | 1 µg/mL |  |  |  | 47 | 46 | 45 |  |

| b. |  | TA100 - S9 |  |  | TA100 + S9 |  |  | Mutagenic Conc./ Cytotoxic effect |
| --- | --- | --- | --- | --- | --- | --- | --- | --- |
|  |  | Number positive wells |  |  | Number positive wells |  |  |  |
| Sample | Concentration (µM) | Replicate #1 | Replicate #2 | Replicate #3 | Replicate #1 | Replicate #2 | Replicate #3 |  |
| T2 | 0 | 2 | 2 | 4 | 3 | 2 | 3 |  |
|  | 8.75 | 2 | 3 | 0 | 0 | 3 | 3 |  |
|  | 17.5 | 1 | 3 | 2 | 4 | 4 | 3 |  |
|  | 35 | 0 | 3 | 1 | 5 | 8 | 1 |  |
|  | 70 | 2 | 3 | 1 | 3 | 4 | 1 |  |
| 2-NQO | 0.1 µg/mL | 46 | 43 | 33 |  |  |  | Mutagenic Conc. |
| 2-AA | 2.5 µg/mL |  |  |  | 48 | 48 | 48 |  |

**Table S8.**
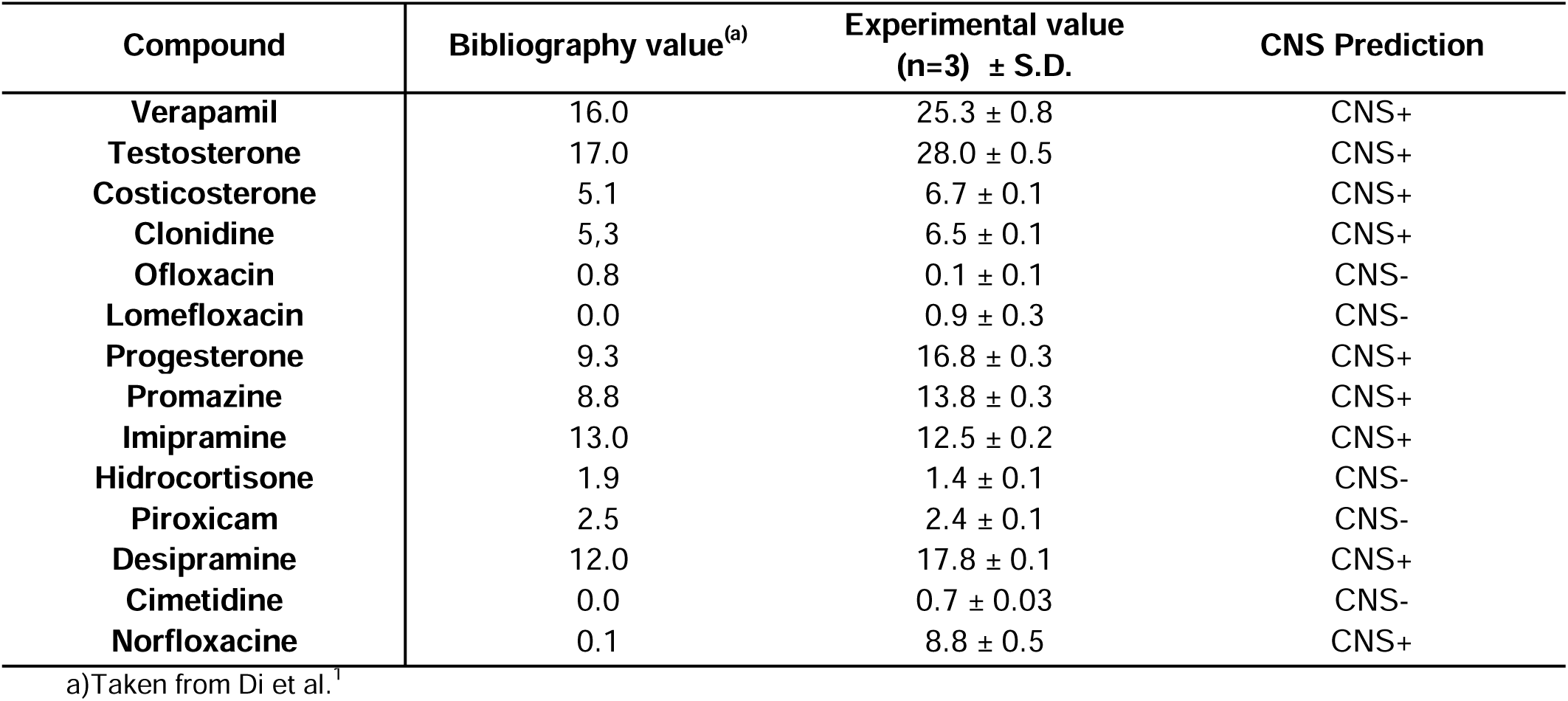
Permeability (*Pe* 10-6 cm s^-1^) in the PAMPA-BBB assay of the 14 commercial drugs predictive penetration in the CNS used as references.

**Table S9.** List of genes from ATAC-seq analysis that overlaps with the Single cell-seq analysis.

| Genes |
| --- |
| Ago2 |
| Alox5ap |
| Bco1 |
| Ccin |
| Cd83 |
| Cnga2 |
| Cop1 |
| D430019H16Rik |
| Ddo |
| Dkk3 |
| Dusp5 |
| Gstk1 |
| Hrh2 |
| Jazf1 |
| Lsm4 |
| Map4k3 |
| Pcgf5 |
| Ppp1r9a |
| Prickle2 |
| Rnase4 |
| Rph3a |
| Sh3bp4 |
| Siah3 |
| Slc44a1 |
| Sub1 |
| Tekt3 |
| Xkr6 |
| Zfp365 |

**Table S10.** Antibodies used in Western Blot.

**Table S11.**
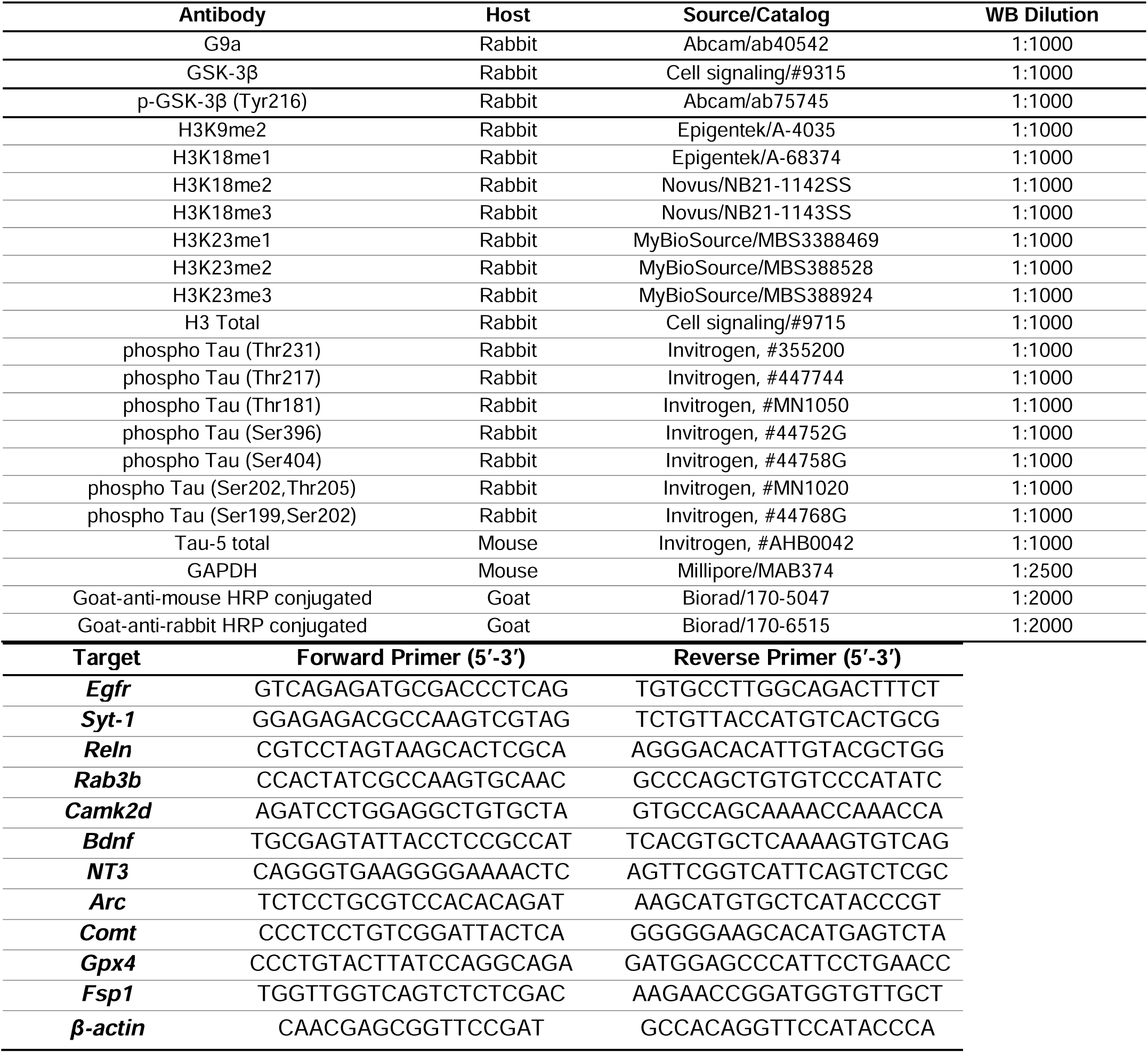
Primer sequences of real-time qPCR (SYBR Green primers).

**Table S12.** Patient data of the parietal brain samples from the Institute of Neuropathology-IDIBELL Brain Bank, Hospitalet de Llobregat, Barcelona, Spain.

| Gender | Age | Postmortem delay | Diagnosis |
| --- | --- | --- | --- |
| Male | 85 | 5 h 45 min | Control |
| Male | 78 | 2 h 15 min | Control |
| Male | 79 | 7h | Control |
| Male | 52 | 3 h 20 min | Control |
| Female | 72 | 2 h 45 min | Control |
| Female | 49 | 5 h | Control |
| Male | 93 | 7 h 20 min | AD BRAAK STAGE<br>V/C |
| Male | 85 | 5 h 15 min | AD BRAAK STAGE<br>V/C |
| Male | 74 | 6 h 30 min | AD BRAAK STAGE<br>V/C |
| Female | 83 | 3 h 20 min | AD BRAAK STAGE<br>V/C |
| Female | 82 | 1 h 45 min | AD BRAAK STAGE<br>V/B |
| Female | 72 | 9 h 30 min | AD BRAAK STAGE<br>V/B |
| Female | 75 | 4 h 15 min | AD BRAAK STAGE<br>V/B |
| Female | 85 | 2 h 10 min | AD BRAAK STAGE<br>V/B |

