## Supplementary 1 for "Dual Inhibitors Targeting G9a and GSK-3β: Translational Perspectives on Alzheimer’s Disease Treatment"

Experimental section...................................................................................................................................................S3

Chromatography and NMR Spectra.............................................................................................................................S10

References................................................................................................................................................................S73

**EXPERIMENTAL SECTION**

**Chemical Synthesis**

Commercially available reagents and solvents were used without further puriﬁcation unless stated otherwise. The progress of the reactions was monitored by thin-layer chromatography (TLC) with aluminium-backed sheets with silica gel 60 F254 (Merck, ref 1.05554), and spots were visualized with UV light. Column chromatography was performed on silica gel 60 Å (Sigma Aldrich, 40 - 63 μm, 230-400 mesh) or with a PuriFlash 5.052 Interchim provided with multi wavelength UV detection using pre-packed RediSep Rf silica gel cartridges normal phase (silica gel, 35−70 μm) and reversed phase (C18) columns with the indicated solvent system. Melting points were determined in open capillary tubes with an MFB 59510M Gallenkamp melting point apparatus. IR spectra were run on a FTIR Perkin-Elmer Spectrum RX I. Absorption values are expressed as wavenumbers (cm^-1^); only significant absorption bands are given. 400 MHz 1H NMR / 100.6 MHz 13C NMR spectra were recorded on a Bruker 400. The chemical shifts were reported in ppm (δ scale) relative to internal tetramethylsilane, or to solvent peak, and coupling constants are reported in Hertz (Hz). Assignments given for the NMR spectra have been carried out based on Distortionless Enhancement by Polarization Transfer (DEPT). The used abbreviations were: s, singlet; d, doublet; t, triplet; q, quadruplet; m, multiplet; cs, complex signal; broad s., broad singlet or combinations thereof. All NMR spectra can be found at the end of this file*.* Accurate mass spectra were recorded with ESI techniques on a Hewlett-Packard 5988a LC / MSD-TOF instrument at Unitat d’Espectrometria de Masses dels Centres Científics i Tecnològics de la Universitat de Barcelona (CCiTUB). The elemental analyses were carried out in a Flash 1112 series Thermofinnigan elemental microanalyzator (A5) to determine C, H, and N at the Servei de Microanàlisis of IIQAB (CSIC) of Barcelona. The analytical samples of all the new compounds, which were subjected to pharmacological evaluation, possessed purity ≥95% as evidenced by their HPLC-UV. HPLC-UV were determined with a HPLC Agilent 1260 Infinity II LC/MSD coupled to a photodiode array and mass spectrometer.  Samples (5 μL, 0.5 mg/mL) in a 1:1 mixture of water with 0.05% formic acid (A) and acetonitrile with 0.05% formic acid (B) were injected using an Agilent Poroshell 120 EC-C18 (2.7 μm, 50 mm × 4.6 mm) column at 40 °C. The mobile phase was a mixture of A and B, with a flow 0.6 mL/min, using the following gradients: from 95% A–5% B to 100% B in 3 min; 100% B for 3 min; from 100% B to 95% A–5% B in 1 min; and 95% A–5% B for 3 min. Purity is given as % of absorbance at 220 nm. HPLC enantioseparation was performed by using HPLC‐7‐NP with a LPG-3400SD pump equipped with a Dionex VWD‐3400‐RS detector. Sample (2000 μL, 2.5 mg/mL) in a 1:1 mixture of hexane (A) and 2-propanol (B) was injected using Phenomenex Lux-Cellulose 4 (250 mm × 0.46 mm, 5 μm) column. The mobile phase was a mixture of A and B, with a flow 1.0 mL/min, using the following gradients: 90% A–10% B for 17 min, 80% A–20% B for 8 min; 90% A–10% B for 4 min; and 90% A–10% B for 15 min.

**General Procedure A:**

The appropriate aldehyde (2.00 mmol), urea or thiourea (6.00 mmol) and ethyl acetoacetate or 1,3-dicarbonyl compound (2.00 mmol) were dissolved in ACN or DMF (1.00 or 2.00 M, respectively), or solvent-free in a 10.00 mL Pyrex microwave process vial. Afterwards, TMSCl (2.00 mmol) were added dropwise to the reaction mixture and the vessel was sealed and subsequently irradiated for 20 to 40 min at 120 ºC under microwave conditions. After it was cooled to room temperature, the reaction mixture was added into cold water, extracted with EA (3x), the combined organic phases were washed with brine, dried over Na_2_SO_4_, filtered and evaporated under reduced pressure. The crude was purified by reverse phase column chromatography to afford the desired product^1^.

**General Procedure B:**

The appropriate aldehyde (1.00 mmol), thiourea (1.00 mmol) and ethyl acetoacetate (1.00 mmol), were dissolved in DMF (2.00 M) in a 10.00 mL Pyrex microwave process vial. Afterwards, TMSCl (6.00 mmol) were added dropwise to the reaction mixture and the vessel was sealed and subsequently irradiated for 1 h at 120 ºC under microwave conditions. After it was cooled to room temperature, the reaction mixture was added into cold water, extracted with EA (3x), the combined organic phases were washed with brine, dried over Na_2_SO_4_, filtered and evaporated under reduced pressure. The crude was purified by reverse phase column chromatography to afford the desired product^2^.

**General Procedure C:**

The appropriate aldehyde (1.00 mmol), urea (1.50 mmol), ethyl acetoacetate (1.00 mmol) and Cu(OTf)_2_ (0.02 mmol) were dissolved in EtOH (0.50 M) in a 10.00 mL Pyrex microwave process vial. The vessel was sealed and subsequently for 1 h at 100 ºC under microwave conditions. After it was cooled to room temperature, a precipitate was formed which was filtered with H_2_O and Hexane. The crude was purified by phase column chromatography to afford the desired product^3^.

**General Procedure D**

A mixture of required aldehyde (1.0 mmol), ethyl acetoacetate or methyl acetoacetate (1.1 mmol), urea or thiourea (1.3 mmol) and citric acid (0.5 mmol) were stirred into a round bottom flask and the reaction mixture was then heated at 80 ºC for 6 to 8 h. After completion of the reaction as monitored by TLC, to the reaction mixture, cold water was added and stirred for 10 minutes. The white precipitates obtained was then filtered, washed with water and dried under vacuum. The crude product was further purified by recrystallization from EtOH^4^.

**Series 1: Ester bond**

1. *Ethyl 6-methyl-4-phenyl-2-thioxo-1,2,3,4-tetrahydropyrimidine-5-carboxylate* ***(T1).***

Product purchased from Apollo Scientific.

2. *Ethyl 4-(4-chlorophenyl)-6-methyl-2-thioxo-1,2,3,4-tetrahydropyrimidine-5-carboxylate (****T2****).*

Following the general **procedure B**, chlorobenzaldehyde (140.40 mg, 1.00 mmol, 1.00 eq), thiourea (76.00 mg, 1.00 mmol, 1.00 eq), ethyl acetoacetate (0.13 mL, 1.00 mmol, 1.09 eq) in 0.50 mL of DMF. Afterwards, 0.76 mL of TMSCl were added. Column chromatography (80.00 % Hex / 20.00 % AcOEt), gave comp **T2** (151.50 mg, 48.80 % yield) as a white solid, mp 196-197 ºC. IR: 3327, 3173, 2984, 1670, 1573, 1465, 1281, 1197, 1178, 1120, 1093, 1014, 805, 760, 746, 646 cm^-1^. ^1^H NMR (400 MHz, CDCl_3_) δ 1.19 (t, *J* = 7.1 Hz, 3H, C*H*_3_), 2.36 (d, *J* = 0.8 Hz, 3H, C*H*_3_), 4.07 – 4.15 (m, 2H, C*H*_2_), 5.38 (d, *J* = 3.1 Hz, 1H, C*H*), 7.13 (s, 1H, -N*H*), 7.20 – 7.25 (m, 2H, Ar*H*), 7.28 – 7.34 (m, 2H, Ar*H*), 7.66 (bs, 1H, -N*H*). ^13^C NMR (101 MHz, DMSO) δ 174.23, 164.99, 145.38, 142.37, 132.26, 128.60, 128.60, 128.31, 128.31, 100.29, 59.66, 53.44, 17.18, 14.01. HRMS (ESI-TOF (-)) calcd for [C_14_H_15_ClN_2_O_2_S, -H]^-^, 309.0540; found 309.0464.

3. *Ethyl 4-(4-methoxyphenyl)-6-methyl-2-thioxo-1,2,3,4-tetrahydropyrimidine-5-carboxylate* ***(T3).***

Following the general **procedure B**, *p*-anisaldehyde (0.12 mL, 1.00 mmol, 1.00 eq), thiourea (76.00 mg, 1.00 mmol, 1.00 eq), ethyl acetoacetate (0.13 mL, 1.00 mmol, 1.00 eq) in 0.50 mL of DMF. Afterwards, 0.76 mL of TMSCl were added. Column chromatography (76.00 % Hex / 24.00 % AcOEt) gave comp **T3** (49.90 mg, 14.70 % yield) as a white solid, mp 156-157 ºC. IR: 3312, 3168, 1666, 1609, 1573, 1509, 1462, 1269, 1251, 1194, 1181, 1170, 1121, 1109, 1028, 835, 819, 765, 654 cm^-1^. ^1^H NMR (400 MHz, CDCl_3_) δ 1.18 (t, *J* = 7.1 Hz, 3H, C*H*_3_), 2.35 (d, *J* = 0.8 Hz, 3H, C*H*_3_), 3.79 (s, 3H, C*H_3_*-Ph), 4.03 – 4.15 (m, 2H, C*H*_2_), 5.35 (d, *J* = 3.0 Hz, 1H, C*H*), 6.83 – 6.87 (m, 2H, Ar*H*), 7.17 – 7.25 (m, 2H, Ar*H*), 7.40 (bs, 1H, -N*H*).^13^C NMR (101 MHz, DMSO) δ 174.01, 165.16, 158.73, 144.74, 135.69, 127.60, 127.60, 113.88, 113.88, 100.95, 59.54, 55.10, 53.43, 17.13, 14.04. HRMS (ESI-TOF (-)) calcd for [C_15_H_18_N_2_O_3_S, -H]^-^, 305.1040; found 305.0958.

4. *Ethyl 6-methyl-2-thioxo-4-(p-tolyl)-1,2,3,4-tetrahydropyrimidine-5-carboxylate* ***(T4).***

Following the general **procedure B**, *p*-tolualdehyde (0.12 mL, 1.00 mmol, 1.00 eq), thiourea (76.00 mg, 1.00 mmol, 1.00 eq), ethyl acetoacetate (0.13 mL, 1.00 mmol, 1.00 eq) in 0.50 mL of DMF. Afterwards, 0.76 mL of TMSCl were added. Column chromatography (83.00 % Hex / 17.00 % AcOEt) gave comp **T4** (65.70 mg, 22.70 % yield) as a white solid, mp 192-193 ºC. IR: 3323, 3170, 1671, 1574. 1463, 1327, 1185, 1174, 1117, 786, 759, 763, 650 cm-^1^. ^1^H NMR (400 MHz, CDCl_3_) δ 1.18 (t, *J* = 7.1 Hz, 3H, C*H*_3_), 2.33 (s, 3H, O C*H*_3_), 2.35 (d, *J* = 0.8 Hz, 3H, C*H*_3_), 4.06 – 4.13 (m, 2H, C*H*_2_), 5.37 (d, *J* = 2.9 Hz, 1H, C*H*), 6.89 (d, *J* = 3.7 Hz, 1H, -N*H*), 7.13 (d, *J* = 8.3 Hz, 2H, Ar*H*), 7.18 (d, *J* = 8.3 Hz, 2H, Ar*H*), 7.42 (bs, 1H, -N*H*).^13^C NMR (101 MHz, DMSO) δ 174.14, 165.14, 144.87, 140.59, 136.90, 129.06, 126.29, 129.06, 126.29, 100.81, 59.56, 53.72, 20.67, 17.14, 14.03. HRMS (ESI-TOF (-)) calcd for [C_15_H_18_N_2_O_2_S -H]^-^, 289.1090; found 289.1007.

5. *Ethyl 4-(3,4-dichlorophenyl)-6-methyl-2-thioxo-1,2,3,4-tetrahydropyrimidine-5-carboxylate* ***(T5).***

Following the general **procedure B**, 3,4-dichlorobenzaldehyde (174.80 mg, 1.00 mmol, 1.00 eq), thiourea (76.00 mg, 1.00 mmol, 1.00 eq), ethyl acetoacetate (0.13 mL, 1.00 mmol, 1.00 eq) in 0.50 mL of DMF. Afterwards, 0.76 mL of TMSCl were added. Column chromatography (62.00 % Hex / 38.00 % AcOEt) gave comp **T5** (47.30 mg, 13.70 % yield) as a white solid, mp 197-198 ºC. IR: 3315, 3173, 1666, 1576, 1466, 1199, 1181, 1112, 1030, 821, 747 cm^-1^. ^1^H NMR (400 MHz, CDCl_3_) δ 1.22 (t, *J* = 7.1 Hz, 3H, C*H*_3_), 2.37 (d, *J* = 0.7 Hz, 3H, C*H*_3_), 4.08 – 4.18 (m, 2H, C*H*_2_), 5.38 (d, *J* = 3.4 Hz, 1H, C*H*), 6.96 (bs, 1H, -N*H*), 7.15 (dd, *J* = 2.2, 8.3 Hz, 1H, Ar*H*), 7.38 (d, *J* = 2.2 Hz, 1H, Ar*H*), 7.42 (d, *J* = 8.3 Hz, 1H, Ar*H*), 7.52 (bs, 1H, -N*H*). ^13^C NMR (101 MHz, DMSO) δ 174.36, 164.86, 145.86, 144.33, 131.05, 131.03, 130.30, 128.51, 126.70, 99.74, 59.75, 53.14, 17.23, 13.98. HRMS (ESI-TOF (-)) calcd for [C_14_H_14_Cl_2_N_2_O_2_S, -H]^-^, 343.0150; found 343.0073.

6. *Ethyl 6-methyl-2-thioxo-4-(4-(trifluoromethyl)phenyl)-1,2,3,4-tetrahydropyrimidine-5-carboxylate* ***(T6)****.*

Following the general **procedure B**, 4-(trifluoromethyl)benzaldehyde (173.90 mg, 1.00 mmol, 1.00 eq), thiourea (76.00 mg, 1.00 mmol, 1.00 eq), ethyl acetoacetate (0.13 mL, 1.0 mmol, 1.0 eq) in 0.5 mL of DMF. Afterwards, 0.76 mL of TMSCl were added. Reverse phase column chromatography (49.00 % CH_3_CN in 0.1 % aq. HCO_2_H) gave comp **T6** (47.70 mg, 13.80 % yield) as a white solid, mp 209-210 ºC. IR: 3320, 3170, 3103, 2993, 1673, 1574, 1467, 1372, 1327, 1285, 1200, 5 1170, 1113, 1066, 1017, 857, 816, 760. 746, 609, 640 cm^-1^. ^1^H NMR (400 MHz, CDCl_3_) δ 1.20 (t, *J* = 7.1 Hz, 3H, C*H*_3_), 2.38 (s, 3H, C*H*_3_), 4.13 (qd, *J* = 3.0, 7.1 Hz, 2H, C*H*_2_), 5.48 (d, *J* = 3.2 Hz, 1H, C*H*), 7.02 (bs, 1H, -N*H*), 7.43 (d, *J* = 8.2 Hz, 2H, Ar*H*), 7.56 (bs, 1H, -N*H*), 7.61 (d, *J* = 8.2 Hz, 2H, Ar*H*). ^13^C NMR (101 MHz, DMSO) δ 13.98, 17.21, 53.73, 59.72, 100.02, 125.59, 125.63, 125.67, 125.70, 127.28, 127.28, 145.68, 147.80, 164.94, 174.42. HRMS (ESI-TOF (+)) calcd for [C_15_H_15_F_3_N_2_O_2_S +H]^+^, 343.0810; found 343.0738.

9. *Ethyl 4-(2-chlorophenyl)-6-methyl-2-thioxo-1,2,3,4-tetrahydropyrimidine-5-carboxylate* ***(T9)***.

Following the general **procedure D,** 2-chlorobenzaldehyde (1.0 g, 7.11 mmol, 1.00 eq), ethyl acetoacetate (0.99 mL, 7.83 mmol, 1.10 eq), thiourea (0.70 g, 9.25 mmol, 1.30 eq) and citric acid (0.68 g, 3.56 mmol, 0.50 eq) gave after recrystallization from ethanol comp **T9** (1.95 g, 88.00 % yield) as a white solid, mp 165 °C. IR: 3663, 3205, 3106, 2979, 1707, 1650, 1573, 1473, 1425, 1377, 1365, 1330, 1284, 1313, 1255, 1203, 1181, 1134, 1121, 1090, 1051, 1037, 1026, 895, 890, 876, 862, 828, 784, 765, 747, 727, 693, 665, 644. ^1^H-NMR (400 MHz, DMSO-*d*_6_): δ 10.37 (s, 1H), 9.06 (s, 1H, -N*H*), 7.42-7.31 (m, 4H, Ar*H*), 5.64 (s, 1H, C*H*-Ph), 3.93-3.92 (m, 2H, OC*H*_2_CH_3_), 2.33 (s, 3H, C*H*_3_), 1.01 (t, 3H, CH_2_C*H*_3_). HRMS (ESI-TOF (+)) calcd for [C_14_H_15_ClN_2_O_2_S +H]^+^, 311.0500; found 311.0616.

10. *Ethyl 4-(4-ethoxyphenyl)-6-methyl-2-thioxo-1,2,3,4-tetrahydropyrimidine-5-carboxylate* ***(T10)***.

Following the general **procedure D**, 4-ethoxybenzaldehyde (1.0 g, 6.66 mmol, 1.00 eq), ethyl acetoacetate (0.93 mL, 7.33 mmol, 1.10 eq), thiourea (0.66 g, 8.66 mmol, 1.30 eq) and citric acid (0.64 g, 3.33 mmol, 0.50 eq) gave after recrystallization from ethanol comp **T10 (**1.90 g, 89.00 % yield) as a white solid, mp 212-214 °C. IR: 3312, 3167, 3102, 2976, 2936, 2903, 1667, 1610, 1574, 1509, 1453, 1426, 1392, 1372, 1330, 1305, 1284, 1265, 1251, 1194, 1181, 1169, 1120, 1111, 1046, 1026, 1000, 954, 925, 889, 873, 853, 836, 819, 794, 767, 740, 720, 694, 655. ^1^H-NMR (400 MHz, DMSO-*d*_6_): δ 10.29 (s, 1H, -N*H*), 9.60 (s,1H, -N*H*), 7.12 (d, 2H, *J* = 8.8 Hz, Ar*H*), 6.89 (d, 2H, *J* = 8.8 Hz, Ar*H*), 5.12 (d, 1H, *J* = 3.2 Hz, C*H*-Ph), 4.03-3.96 (m, 2H, OC*H*_2_CH_3,_ 2H, COOC*H*_2_CH_3_), 2.29 (s, 3H, C*H*_3_), 1.30 (t, 3H, *J* = 7.2 Hz, CH_2_C*H*_3_), 1.10 (t, 3H, *J* = 7.2 Hz, CH_2_C*H*_3_). HRMS (ESI-TOF (+)) calcd for [C_16_H_20_N_2_O_3_S +H]^+^, 321.1200; found 321.1272.

11. *Methyl 4-(2-methoxyphenyl)-6-methyl-2-thioxo-1,2,3,4-tetrahydropyrimidine-5-carboxylate* ***(T11)***.

Following the general **procedure D**, 2-methoxybenzaldehyde (1.0 g, 7.34 mmol, 1.00 eq), methyl acetoacetate (0.87 mL, 8.07 mmol, 1.10 eq), thiourea (0.57 g, 9.55 mmol, 1.30 eq) and citric acid (0.71 g, 3.67 mmol, 0.50 eq) gave after recrystallization from ethanol comp **T11** (1.85 g, 86.00 % yield) as a white solid, mp 284-286 °C. IR: 3311, 3167, 3102, 2976, 2936, 1697, 1655, 1610, 1573, 1509, 1474, 1452, 1427, 1391, 1372, 1330, 1306, 1284, 1265, 1251, 1241, 1193, 1180, 1169, 1120, 1103, 1090, 1047, 1025, 954, 925, 889, 873, 853, 836, 818, 794, 764, 720, 693, 655. ^1^H-NMR (400 MHz, DMSO-*d*_6_): δ 10.27 (s, 1H, -N*H*), 9.24 (s,1H, -N*H*), 7.29-7.25 (m, 1H, Ar*H*), 7.05-7.00 (m, 2H, Ar*H*), 6.92-6.88 (m, 1H, Ar*H*), 5.50 (d, 1H, *J* =3.6 Hz, C*H*-Ph), 3.80 (s, 3H, -OC*H*_3_), 3.51 (s, 3H, COOC*H*_3_), 2.31 (s, 3H, C*H*_3_). HRMS (ESI-TOF (+)) calculated for C_14_H_16_N_2_O_3_S [M+H] = 293.0900, found 293.0950.

12. *Ethyl 4-(3-hydroxyphenyl)-6-methyl-2-thioxo-1,2,3,4-tetrahydropyrimidine-5-carboxylate* ***(T12)***.

Following the general **procedure D**, 3-hydroxybenzaldehyde (1.0 g, 8.19 mmol, 1.00 eq), ethyl acetoacetate (1.15 mL, 9.00 mmol, 1.10 eq), thiourea (0.81 g, 10.64 mmol, 1.30 eq) and citric acid (0.79 g, 4.10 mmol, 0.50 eq) gave after recrystallization from ethanol comp **T12** (2.03 g, 85% yield) as a white solid, mp 183-184 ºC. IR: 3513, 3236, 2977, 1698, 1670, 1644, 1599, 1508, 1450, 1370, 1340, 1316, 1279, 1182, 1091, 1024, 956, 929, 871, 789, 765, 701, 655, 615. ^1^H-NMR (400 MHz, DMSO-*d*_6_): 10.30 (s, 1H, -N*H*), 9.60 (s, 1H, -N*H*), 9.44 (s, 1H, -O*H*), 7.14-7.10 (m, 1H, Ar*H*),  6.64-6.62 (m, 3H, Ar*H*),  5.09 (d, 1H, *J* = 3.6 Hz, C*H*-Ph), 4.05-3.99 (m, 2H, C*H*_2_CH_3_), 2.28 (s, 3H, C*H*_3_), 1.13 (t, 3H, CH_2_C*H*_3_). HRMS (ESI-TOF (+)) calcd for [C_14_H_16_N_2_O_3_S +H]^+^, 293.0900; found 293.0954.

13. *Ethyl 4-(4-fluorophenyl)-6-methyl-2-thioxo-1,2,3,4-tetrahydropyrimidine-5-carboxylate* ***(T13).***

Following the general **procedure D,** 4-fluorobenzaldehyde (1.0 g, 8.06 mmol, 1.00 eq), ethyl acetoacetate (1.13 mL, 8.86 mmol, 1.10 eq), thiourea (0.80 g, 10.47 mmol, 1.30 eq) and citric acid (0.77 g, 4.03 mmol, 0.50 eq) gave after recrystallization from ethanol comp **T13** (2.02 g, 85% yield) as a white solid, mp 210 ºC. IR: 3325, 3172, 2975, 1680, 1604, 1574, 1506, 1464, 1393, 1370, 1327, 1300, 1283, 1230, 1194, 1177, 1158, 1118, 1030, 955, 923, 873, 854, 837, 824, 810, 791, 758, 741, 707, 682, 650. ^1^H-NMR (400 MHz, DMSO-*d*_6_): δ 10.37 (s, 1H, -N*H*), 9.66 (s, 1H, -N*H*), 7.27-7.16 (m, 4H, Ar*H*), 5.18 (d, 1H, *J* = 3.6 Hz, C*H*-Ph), 4.04-3.98 (m, 2H, C*H*_2_CH_3_), 2.30 (s, 3H, C*H*_3_), 1.10 (t, 3H, *J* = 7.2 Hz, CH_2_C*H*_3_). HRMS (ESI-TOF (+)) calcd for [C_14_H_15_FN_2_O_2_S +H]^+^, 295.0800; found 295.0911.

14. *Methyl 4-(3-methoxyphenyl)-6-methyl-2-thioxo-1,2,3,4-tetrahydropyrimidine-5-carboxylate* ***(T14).***

Following the general **procedure D,** 3-methoxybenzaldehyde (1.00 g, 7.34 mmol, 1.00 eq), methyl acetoacetate (0.87 mL, 8.08 mmol, 1.10 eq), thiourea (0.73 g, 9.55 mmol, 1.30 eq) and citric acid (0.71 g, 3.67 mmol, 0.50 eq) gave after recrystallization from ethanol comp **T14** (1.87 g, 87.00 % yield) as a white solid, mp 171-173 °C. IR: 3317, 3175, 1658, 1598, 1571, 1488, 1433, 1382, 1345, 1284, 1269, 1251, 1182, 1114, 1040, 994, 946, 901, 876, 800, 757, 724, 700, 656, 624. ^1^H-NMR (400 MHz, DMSO-*d*_6_): δ 10.36 (s, 1H, -N*H*), 9.66 (s, 1H, -N*H*), 7.29-7.25 (t, 1H, Ar*H*), 6.87-6.77 (m, 3H, Ar*H*), 5.16 (d, 1H, *J* = 4 Hz, C*H*-Ph), 3.74 (s, 3H, OC*H*_3_), 3.58 (s, 3H, COOC*H*_3_), 2.29 (s, 3H, C*H*_3_). HRMS (ESI-TOF (+)) calcd for [C_14_H_16_N_2_O_3_S +H]^+^, 293.0900; found 293.0954.

15. ***Ethyl 6-methyl-2-oxo-4-phenyl-1,2,3,4-tetrahydropyrimidine-5-carboxylate (U1)***

Product purchased from Fischer Scientific (TCI).

16. ***Ethyl 4-(4-chlorophenyl)-6-methyl-2-oxo-1,2,3,4-tetrahydropyrimidine-5-carboxylate (U2)***

Product purchased from Sigma-Aldrich.

17. *Ethyl 6-methyl-2-oxo-4-(p-tolyl)-1,2,3,4-tetrahydropyrimidine-5-carboxylate* ***(U3).***

Following the general **procedure A,** *p*-anisaldehyde (0.24 mL, 2.00 mmol, 1.00 eq), urea (360.30 mg, 6.00 mmol, 3.00 eq), ethyl acetoacetate (0.25 mL, 2.0 0mmol, 1.00 eq). Afterwards, 0.25 mL of TMSCl were added, and subsequently the mixture was irradiated for 20 min at 120 ºC under microwave conditions Column chromatography (5.00 % MeOH / 95.00 % DCM) gave comp **U3** (194.00 mg, 33.40 % yield) as a white solid, mp 213-214 ºC. IR: 3236, 3113, 1724, 1704, 1651, 1514, 1455, 1278, 1257, 1222, 1177, 1086, 1031, 781, 659 cm^-1^. ^1^H NMR (400 MHz, CDCl_3_) δ 1.17 (t, *J* = 7.1 Hz, 3H, C*H*_33_), 2.34 (d, *J* = 0.7 Hz, 3H, C*H*_3_), 3.79 (s, 3H, Ar- C*H*_3_), 4.03 – 4.13 (m, 2H, C*H*_2_), 5.36 (d, *J* = 2.7 Hz, 1H, C*H*), 5.38 (bs, 1H, -N*H*), 6.81 – 6.86 (m, 2H, Ar*H*), 7.19 (bs, 1H, -N*H*), 7.23 (d, *J* = 2.1 Hz, 1H, Ar*H*), 7.25 (d, *J* = 2.1 Hz, 1H, Ar*H*).^13^C NMR (101 MHz, DMSO) δ 165.37, 158.44, 152.15, 148.01, 137.06, 127.39, 127.39, 113.70, 113.70, 99.56, 59.15, 55.06, 53.33, 17.76, 14.11. HRMS (ESI-TOF (-)) calcd for [C_15_H_18_N_2_O_4_ -H]^-^, 289.1270; found 289.1189.

18. ***Ethyl 6-methyl-2-oxo-4-(p-tolyl)-1,2,3,4-tetrahydropyrimidine-5-carboxylate (U4).***

Product purchased from Sigma-Aldrich.

19. *Ethyl 4-(3,4-dichlorophenyl)-6-methyl-2-oxo-1,2,3,4-tetrahydropyrimidine-5-carboxylate* ***(U5).***

Following the general **procedure A**, 3,4-dichlorobenzaldehyde (350.00 mg, 2.00 mmol, 1.00 eq), urea (360.30 mg, 6.00 mmol, 3.00 eq), ethyl acetoacetate (0.25 mL, 2.00 mmol, 1.00 eq). Afterwards, 0.25 mL of TMSCl were added, and subsequently the mixture was irradiated for 20 min at 120 ºC under microwave conditions. Column chromatography (4.00 % MeOH / 96.00 % DCM) gave comp **U5** (197.90 mg, 30.10 % yield) as a white solid, mp 213-214 ºC. IR (ATR) 3350, 3107, 2974, 1694, 1645, 1461, 1320, 1297, 1227, 1099, 872, 799, 721, 663 cm^-1^. ^1^H NMR (400 MHz, CDCl_3_) δ 1.21 (t, *J* = 7.1 Hz, 3H, C*H*_3_), 2.37 (d, *J* = 0.7 Hz, 3H, C*H*_3_), 4.06 – 4.15 (m, 2H, C*H*_2_), 5.38 (d, *J* = 3.1 Hz, 1H, C*H*), 5.44 (bs, 1H, -N*H*), 7.17 (dd, *J* = 2.2, 2.4 Hz, 1H, Ar*H*), 7.38 (s, 1H, Ar*H*), 7.41 (d, *J* = 1.9 Hz, 1H, Ar*H*). ^13^C NMR (101 MHz, DMSO) δ 165.08, 151.74, 149.27, 145.82, 130.87, 130.82, 129.80, 128.44, 126.59, 98.20, 59.35, 53.18, 17.85, 14.05. HRMS (ESI-TOF (-)) calcd for [C_14_H_14_Cl_2_N_2_O_3_ -H]^-^, 327.0380; found 327.0302.

20. *Ethyl 6-methyl-2-oxo-4-(4-(trifluoromethyl)phenyl)-1,2,3,4-tetrahydropyrimidine-5-carboxylate (****U6).***

Following the general **procedure A**, 4-(trifluoromethyl)benzaldehyde (348.20 mg, 2.00 mmol, 1.00 eq), urea (360.30 mg, 6.00 mmol, 3.00 eq), ethyl acetoacetate (0.25 mL, 2.00 mmol, 1.00 eq). Afterwards, 0.25 mL of TMSCl were added, and subsequently the mixture was irradiated for 20 min at 120 ºC under microwave conditions. Reverse phase column chromatography (45.00 CH_3_CN in 0.1 % aq. HCO_2_H)) gave comp **U6** (275.60 mg, 42.00 % yield) as a white solid, mp 189-190 ºC. IR: 3242, 3113, 1701, 1644, 1333, 1289, 1221, 1126, 1091, 1069, 1018, 790, 778, 715, 665 cm-1. ^1^H NMR (400 MHz, CDCl3) δ 1.18 (t, *J* = 7.1 Hz, 3H, C*H*_3_), 2.36 (s, 3H, C*H*_3_), 4.09 (qd, *J* = 1.3, 7.1 Hz, 2H, C*H*_2_), 5.47 (d, *J* = 3.0 Hz, 1H, C*H*), 5.65 (bs, 1H, -N*H*), 7.45 (d, *J* = 8.1 Hz, 2H, Ar*H*), 7.54 (bs, 1H, -N*H*), 7.58 (d, *J* = 8.2 Hz, 2H, Ar*H*). ^13^C NMR (101 MHz, DMSO) δ 14.04, 17.84, 53.74, 59.32, 98.51, 125.40, 125.44, 125.48, 125.52, 127.14, 127.14, 149.09, 149.28, 151.88, 165.16. HRMS (ESI-TOF (-)) calcd for [C_15_H_15_F_3_N_2_O_3_, -H]^-^, 327.1030; found 327.0964.

21. *Ethyl 4-(2,4-dichlorophenyl)-6-methyl-2-oxo-1,2,3,4-tetrahydropyrimidine-5-carboxylate* ***(U7).***

Following the general **procedure A,** 2,4-dichlorobenzaldehyde (350.00 mg, 2.00 mmol, 1.00 eq), urea (360.30 mg, 6.00 mmol, 3.00 eq), ethyl acetoacetate (0.25 mL, 2.00 mmol, 1.00 eq). Afterwards, 0.25 mL of TMSCl were added, and subsequently the mixture was irradiated for 20 min at 120 ºC under microwave conditions. Reverse phase column chromatography (42.00 % ACN / 58.00 % H2O: 0.1% FA) gave comp **U7** (32.30 mg, 4.90 % yield) as a white solid, mp 248-249 ºC. ^1^H NMR (400 MHz, CDCl_3_) δ 1.09 (t, *J* = 7.1 Hz, 3H, C*H*_3_), 2.45 (s, 3H, C*H*_3_), 4.02 (qd, *J* = 2.0, 7.1 Hz, 2H, C*H*_2_), 5.56 (bs, 1H, -N*H*), 5.83 (d, *J* = 3.0 Hz, 1H, C*H*), 7.17 (d, *J* = 8.4 Hz, 1H, Ar*H*), 7.22 (dd, *J* = 2.0, 8.4 Hz, 1H, Ar*H*), 7.34 (s, 1H, -N*H*), 7.41 (d, *J* = 2.0 Hz, 1H, Ar*H*). ^13^C NMR (101 MHz, DMSO) δ 13.92, 17.69, 51.17, 59.14, 97.47, 127.98, 128.69, 130.28, 132.55, 132.65, 140.96, 149.58, 151.13, 164.83. HRMS (ESI-TOF (-)) calcd for [C_14_H_14_Cl_2_N_2_O_3_ -H]^-^, 327.0380; found 327.0323.

22. *Ethyl 6-methyl-4-(4-nitrophenyl)-2-oxo-1,2,3,4-tetrahydropyrimidine-5-carboxylate* ***(U8).***

Following the general **procedure D,** 4-nitrobenzaldehyde (1.00 g, 6.61 mmol, 1.00 eq), ethyl acetoacetate (0.93 mL, 7.28 mmol, 1.10 eq), urea (0.52 g, 8.60 mmol, 1.30 eq) and citric acid (0.64 g, 3.31 mmol, 0.50 eq) gave after recrystallization from ethanol comp **U8** (1.90 g, 94.00 % yield) as a white solid, mp 206-208 °C. . IR: 3226, 3112, 2976, 1725, 1697, 1640, 1606, 1595, 1517, 1492, 1424, 1463, 1391, 1376, 1347, 1317, 1304, 1290, 1209, 1181, 1123, 1094, 1083, 1021, 1011, 960, 881, 867, 855, 771, 740, 696, 678, 656, 627. ^1^H-NMR (400 MHz, DMSO-*d*_6_): δ 9.36 (s, 1H, -N*H*), 8.22 (d, 2H, *J* = 8.8 Hz, Ar*H*), 7.89 (s, 1H, -N*H*), 7.51 (d, 2H, *J* = 8.8 Hz, Ar*H*), 5.28 (d, 1H, *J* = 3.2 Hz, C*H*-Ph), 3.99 (q, 2H, *J* = 7.2 Hz, C*H*_2_CH_3_), 2.27 (s, 3H, C*H*_3_), 1.10 (t, 3H, *J* = 7.2 Hz, CH_2_C*H*_3_). HRMS (ESI-TOF (+)) calcd for [C_14_H_15_N_3_O_5_ +H]^+^, 306.1000; found 306.1084.

23. *Ethyl 4-(2-bromophenyl)-6-methyl-2-oxo-1,2,3,4-tetrahydropyrimidine-5-carboxylate* ***(U9).***

Following the general **procedure D,** 2-bromobenzaldehyde (1.00 g, 5.40 mmol, 1.00 eq), ethyl acetoacetate (0.76 mL, 5.95 mmol, 1.10 eq), urea (0.42 g, 7.03 mmol, 1.30 eq) and citric acid (0.64 g, 3.31 mmol, 0.50 eq) gave after recrystallization from ethanol comp **U9** (1.70 g, 93.00 % yield) as a white solid, mp 207-209 °C. . IR: 3342, 3220, 3108, 2977, 1688, 1637, 1567, 1439, 1369, 1337, 1317, 1299, 1250, 1226, 1161, 1142, 1096, 1023, 958, 876, 859, 845, 827, 808, 785, 745, 722, 648, 613. ^1^H-NMR (400 MHz, DMSO-*d*_6_): δ 9.28 (s, 1H, -N*H*), 7.69 (s,1H, -N*H*), 7.59-7.17 (m, 4H, Ar*H*), 5.61 (d, 1H, *J* = 2.8 Hz, C*H*-Ph), 3.89 (q, 2H, *J* = 6.8 Hz, C*H*_2_CH_3_), 2.31 (s, 3H, C*H*_3_), 1.00 (t, 3H, *J* = 6.8 Hz, CH_2_C*H*_3_). HRMS (ESI-TOF (+)) calcd for [C_14_H_15_BrN_2_O_3_ +H]^+^, 339.0300; found 339.0339.

24. *Ethyl 4-(3-bromophenyl)-6-methyl-2-oxo-1,2,3,4-tetrahydropyrimidine-5-carboxylate* ***(U10).***

Following the general **procedure D,** 3-bromobenzaldehyde (1.00 g, 5.40 mmol, 1.0 eq), ethyl acetoacetate (0.76 mL, 5.95 mmol, 1.10 eq), urea (0.42 g, 7.03 mmol, 1.30 eq) and citric acid (0.64 g, 3.31 mmol, 0.50 eq). gave after recrystallization from ethanol comp **U10** *(*1.66 g, 91.00 % yield) s a white solid, mp 185-186 °C IR: 3219, 3096, 2977, 1702, 1651, 1590, 1570, 1474, 1426, 1389, 1378, 1365, 1330, 1308, 1284, 1266, 1223, 1183, 1150, 1120, 1089, 1024, 953, 924, 889, 875, 862, 797, 783, 767, 750, 693, 665, 603. ^1^H-NMR (400 MHz, DMSO-*d*_6_): δ 9.28 (s, 1H, -N*H*), 7.80 (s,1H, -N*H*), 7.47-7.23 (m, 4H, Ar*H*), 5.15 (d, 1H, *J* = 3.2Hz, C*H*-Ph), 4.05-3.96 (m, 2H, C*H*_2_CH_3_), 2.26 (s, 3H, C*H*_3_), 1.11 (t, 3H, *J* = 7.2 Hz, CH_2_C*H*_3_). HRMS (ESI-TOF (+)) calcd for [C_14_H_15_BrN_2_O_3_ +H]^+^, 339.0300; found 339.0339.

25. *Ethyl 4-(3-bromo-4-fluorophenyl)-6-methyl-2-oxo-1,2,3,4-tetrahydropyrimidine-5-carboxylate* ***(U11).***

Following the general **procedure D,** 3-bromo-4-fluorobenzaldehyde (1.00 g, 4.93 mmol, 1.00 eq), ethyl acetoacetate (0.69 mL, 5.42 mmol, 1.10 eq), urea (0.38 g, 6.40 mmol, 1.30 eq) and citric acid (0.47 g, 2.46 mmol, 0.50 eq) gave after recrystallization from ethanol comp **U11** (1.60 g, 91.00 % yield) as a white solid, mp 193-195 °C. IR: 3343, 3202, 3094, 2977, 1698, 1687, 1655, 1636, 1491, 1429, 1391, 1369, 1329, 1300, 1275, 1230, 1193, 1171, 1095, 1047, 1023, 1004, 952, 896, 878, 866, 827, 819, 803, 783, 768, 752, 713, 657, 640, 628. ^1^H-NMR (400 MHz, DMSO-*d*_6_): δ 9.29 (s, 1H, -N*H*), 7.79 (s,1H, -N*H*), 7.51-7.25 (m, 4H, Ar*H*), 5.16 (d, 1H, *J* = 3.2 Hz, C*H*-Ph), 4.03-3.95 (m, 2H, C*H*_2_CH_3_), 2.26 (s, 3H, C*H*_3_), 1.10 (t, 3H, *J* = 7.2 Hz, CH_2_C*H*_3_). HRMS (ESI-TOF (+)) calcd for [C_14_H_14_BrFN_2_O_3_ +H]^+^, 357.0200; found 357.0245.

26. *Methyl 4-(3-bromophenyl)-6-methyl-2-oxo-1,2,3,4-tetrahydropyrimidine-5-carboxylate* ***(U12).***

Following the general **procedure D,** 3-bromobenzaldehyde (1.00 g, 5.40 mmol, 1.00 eq), methyl acetoacetate (0.42 mL, 5.94 mmol, 1.10 eq), urea (0.42 g, 7.03 mmol, 1.30 eq) and citric acid (0.52 g, 2.70 mmol, 0.50 eq) gave after recrystallization from ethanol comp **U12** (1.62 g, 92.00 % yield) as a white solid, mp  228 °C. IR: 3310, 1711, 1673, 1652, 1571, 1509, 1435, 1330, 1274, 1234, 1190, 1091, 1025, 940, 851, 802, 786, 768, 757, 726, 703, 682, 654, 621. ^1^H-NMR (400 MHz, DMSO-*d*_6_): δ 9.30 (s, 1H, -N*H*), 7.82 (s, 1H, -N*H*), 7.47-7.22 (m, 4H, Ar*H*), 5.15 (d, 1H, *J* = 3.2 Hz, C*H*-Ph), 3.55 (s, 3H, COOC*H*_3_), 2.27 (s, 3H, C*H*_3_). HRMS (ESI-TOF (+)) calcd for [C_13_H_13_BrN_2_O_3_ +H]^+^, 325.0100; found 325.0182.

27. *Ethyl 4-(3-hydroxyphenyl)-6-methyl-2-oxo-1,2,3,4-tetrahydropyrimidine-5-carboxylate* ***(U13).***

Following the general **procedure D,** 3-hydroxybenzaldehyde (1.00 g, 8.19 mmol, 1.00 eq), ethyl acetoacetate (1.15 mL, 9.00 mmol, 1.10 eq), urea (0.64 g, 10.64 mmol, 1.30 eq) and citric acid (0.79 g, 4.10 mmol, 0.50 eq) gave after recrystallization from ethanol comp **U13** (1.95 g, 86.00 % yield) as a white solid, mp 180-183 °C. IR: 3513, 3343, 3236, 3117, 2977, 1723, 1708, 1698, 1674, 1643, 1634, 1600, 1452, 1423, 1372, 1315, 1296, 1279, 1219, 1182, 1163, 1120, 1089, 1025, 997, 953, 929, 872, 854, 793, 775, 703, 655, 615, 604. ^1^H-NMR (400 MHz, DMSO-*d*_6_): 9.35 (s, 1H, -N*H*), 9.15 (s, 1H, -N*H*), 7.68 (s,1H, -O*H*), 7.12-7.08 (m, 1H, Ar*H*), 6.68-6.61 (m, 3H, Ar*H*),  5.06 (d, 1H, *J* = 3.6 Hz, C*H*-Ph), 4.05 (q, 2H, *J* = 7.2 Hz, C*H*_2_CH_3_), 2.24 (s, 3H, C*H*_3_), 1.12 (t, 3H, *J* = 7.2 Hz, CH_2_C*H*_3_). HRMS (ESI-TOF (+)) calcd for [C_14_H_16_N_2_O_4_ +H]^+^, 277.1100; found 277.1183.

28. *Ethyl 4-(4-fluorophenyl)-6-methyl-2-oxo-1,2,3,4-tetrahydropyrimidine-5-carboxylate* ***(U14).***

Following the general **procedure D,** 4-fluorobenzaldehyde (1.00 g, 8.06 mmol, 1.00 eq), ethyl acetoacetate (1.13 mL, 8.86 mmol, 1.10 eq), urea (0.63 g, 10.47 mmol, 1.30 eq) and citric acid (0.77 g, 4.03 mmol, 0.50 eq) gave after recrystallization from ethanol comp **U14** (2.06 g, 92.00 % yield) as a white solid, mp 171-173 °C. IR: 3235, 3112, 2978, 1699, 1673, 1645, 1599, 1508, 1461, 1384, 1368, 1324, 1286, 1220, 1161, 1088, 1046, 1025, 1011, 954, 868, 839, 791, 776, 694, 676, 655, 631, 605. ^1^H-NMR (400 MHz, DMSO-*d*_6_): δ 9.23 (s, 1H, -N*H*), 7.75 (s, 1H, -N*H*), 7.29-7.25 (m, 2H, Ar*H*), 7.18-7.13 (m, 2H, Ar*H*), 5.15 (d, 1H, *J* = 3.2 Hz, C*H*-Ph), 4.01-3.96 (m, 2H, C*H*_2_CH_3_), 2.26 (s, 3H, C*H*_3_), 1.09 (t, 3H, *J* = 7.2 Hz, CH_2_C*H*_3_). HRMS (ESI-TOF (+)) calcd for [C_14_H_15_FN_2_O_3_ +H]^+^, 279.1100; found 279.1139.

29. *Methyl 6-methyl-2-oxo-4-phenyl-1,2,3,4-tetrahydropyrimidine-5-carboxylate* ***(U15).***

Following the general **procedure D,** benzaldehyde (1.0 g, 9.42 mmol, 1.00 eq), methyl acetoacetate (1.12 mL, 10.37 mmol, 1.10 eq), urea (0.74 g, 12.25 mmol, 1.30 eq) and citric acid (0.91 g, 4.71 mmol, 0.50 eq) gave after recrystallization from ethanol comp **U15** (2.13 g, 92.00 % yield) as a white solid, mp 210-211 °C. IR: 3328, 3103, 2950, 1692, 1663, 1600, 1433, 1412, 1338, 1273, 1236, 1188, 1092, 1029, 937, 855, 823, 791, 755, 747, 697, 668, 638, 623, 615. ^1^H-NMR (400 MHz, DMSO-*d*_6_): δ 9.23 (s, 1H, -N*H*), 7.76 (s, 1H, -N*H*), 7.35-7.31 (m, 2H, Ar*H*), 7.26-7.23 (m, 3H, Ar*H*), 5.15 (d, 1H, *J* = 3.2 Hz, C*H*-Ph), 3.54 (s, 3H, COOC*H*_3_), 2.26 (s, 3H, C*H*_3_).HRMS (ESI-TOF (+)) calcd for [C_13_H_14_N_2_O_3_ [M+H]^+^, 247.1000; found 247.1077.

**Series 2: Amide bond**

30. *N*-(4-chlorophenyl)-6-methyl-2-oxo-4-phenyl-1,2,3,4-tetrahydropyrimidine-5-*carboxamide* ***(CU1).***

Following the general **procedure C**, benzaldehyde (0.07 mL, 0.71 mmol, 1.00 eq), urea (63.80 mg, 1.06 mmol, 1.50 eq), 1.2, *N*-(4-chlorophenyl)-3-oxobutanamide (150.00 mg, 0.71 mmol, 1.00 eq), Cu(OTf)_2_ (7.20 mg, 0.02 mmol, 0.02 eq) in 1.40 mL of EtOH. Column chromatography (95.00 % DCM / 5.00 % MeOH) gave comp **CU1** (133.20 mg, 55.00 % yield) as a white solid, mp 266-267 ºC. IR: 3401, 3271, 1711, 1669, 1629, 1508, 1492, 1396, 1321, 1243, 1089, 1013, 819, 748, 697, 682, 656, 624 cm^-1^. ^1^H NMR (400 MHz, MeOD) δ 2.08 (d, *J* = 1.0 Hz, 3H, C*H*_3_), 5.47 (d, *J* = 1.0 Hz, 1H, C*H*), 7.23 – 7.28 (m, 3H, Ar*H*), 7.31 – 7.36 (m, 4H, Ar*H*), 7.39 – 7.43 (m, 2H, Ar*H*). ^13^C NMR (101 MHz, DMSO) δ 165.39, 152.54, 144.29, 139.00, 138.20, 128.49, 128.49, 128.40, 128.40, 127.34, 126.60, 126.20, 126.20, 121.06, 121.06, 105.14, 54.97, 17.08. HRMS (ESI-TOF (-)) calcd for [C_18_H_16_ClN_3_O_2_ -H]^-^, 340.0930; found 340.0857.

31. *N, 4-bis(4-chlorophenyl)-6-methyl-2-oxo-1,2,3,4-tetrahydropyrimidine-5-carboxamide* ***(CU2).***

Following the general **procedure C**, 4-chlorobenzaldehyde (99.60 mg, 0.71 mmol, 1.00 eq), urea (63.80 mg, 1.06 mmol, 1.50 eq), 1.2, *N*-(4-chlorophenyl)-3-oxobutanamide (150.00 mg, 0.71 mmol, 1.00 eq), Cu(OTf)_2_ (7.20 mg, 0.02 mmol, 0.02 eq) in 1.40 mL of EtOH. Column chromatography (95.00 % DCM / 5.00 % MeOH) comp **CU2** (69.70 mg, 26.10 % yield) as a white solid, mp 273-274 ºC. IR: 3404, 3268, 3108, 1707, 1669, 1626, 1510, 1490, 1397, 1320, 1241, 1089, 1012, 817, 756, 681, 639, 625 cm^-1^. ^1^H NMR (400 MHz, MeOD) δ 2.08 (d, *J* = 1.0 Hz, 3H, C*H*_3_), 5.45 (d, *J* = 1.0 Hz, 1H, C*H*), 7.24 – 7.28 (m, 2H, Ar*H*), 7.33 (d, *J* = 1.0 Hz, 4H, Ar*H*), 7.41 – 7.44 (m, 2H, Ar*H*). ^13^C NMR (101 MHz, DMSO) δ 165.25, 152.38, 143.18, 139.34, 138.11, 131.89, 128.48, 128.48, 128.42, 128.42, 128.14, 128.14, 126.68, 121.10, 121.10, 104.72, 54.40, 17.10. HRMS (ESI-TOF (-)) calcd for C_18_H_15_Cl_2_N_3_O_2_ -H]^-^, 374.0540; found 374.0468.

32. *N-(4-chlorophenyl)-4-(4-methoxyphenyl)-6-methyl-2-oxo-1,2,3,4-tetrahydropyrimidine-5-carboxamide* ***(CU3).***

Following the general **procedure C**, *p*-anisaldehyde (0.09 mL, 0.71 mmol, 1.00 eq), urea (63.80 mg, 1.06 mmol, 1.5 eq), 1.2, *N*-(4-chlorophenyl)-3-oxobutanamide (150.00 mg, 0.71 mmol, 1.00 eq), Cu(OTf)_2_ (7.20 mg, 0.02 mmol, 0.02 eq) in 1.40 mL of EtOH. Column chromatography (95.00 % DCM / 5.00 % MeOH) comp **CU3** (87.10 mg, 33.10 % yield) as a white solid, mp 256-257 ºC. IR: 3401, 3271, 3113, 2929, 1707, 1670, 1628, 1508, 1395, 1322, 1243, 1173, 1088, 1032, 820, 755, 725, 682, 648 cm^-1^. ^1^H NMR (400 MHz, MeOD) δ 2.08 (d, *J* = 1.1 Hz, 3H, C*H*_3_), 3.75 (s, 3H, ArC*H*_3_), 5.41 (d, *J* = 1.2 Hz, 1H, CH), 6.85 – 6.90 (m, 2H, Ar*H*), 7.23 – 7.28 (m, 4H, Ar*H*), 7.38 – 7.44 (m, 2H, Ar*H*). ^13^C NMR (101 MHz, DMSO) δ 165.42, 158.55, 152.46, 138.84, 138.24, 136.43, 128.40, 128.40, 127.48, 127.48, 126.58, 126.58, 121.05, 121.05, 113.82, 105.32, 55.07, 54.43, 17.07. HRMS (ESI-TOF (+)) calcd for [C_19_H_18_ClN_3_O_3_ +H]^+^, 372.1040; found 372.1108.

33. *N*-(4-chlorophenyl)-6-methyl-2-oxo-4-(p-tolyl)-1,2,3,4-tetrahydropyrimidine-5-*carboxamide* ***(CU4).***

Following the general **procedure C**, *p*-toluadlehyde (0.08 mL, 0.71 mmol, 1.00 eq), urea (63.80 mg, 1.06 mmol, 1.50 eq), 1.2, *N*-(4-chlorophenyl)-3-oxobutanamide (150.00 mg, 0.71 mmol, 1.00 eq), Cu(OTf)_2_ (7.20 mg, 0.02 mmol, 0.02 eq) in 1.40 mL of EtOH. Column chromatography (95.00 % DCM / 5.00 % MeOH) comp **CU4** (157.20 mg, 62.30 % yield) as a white solid, mp 263-264 ºC. IR 3397, 3278, 1707, 1668, 1628, 1592, 1508, 1397, 1244, 1231, 1090, 1014, 820, 764, 717, 682, 638, 648, 626 cm^-1^. ^1^H NMR (400 MHz, MeOD) δ 2.07 (d, *J* = 1.1 Hz, 3H, C*H*_3_), 2.28 (s, 3H, OC*H*_3_) 5.42 (d, *J* = 1.2 Hz, 1H, C*H*), 7.12 – 7.15 (m, 2H, Ar*H*), 7.20 – 7.22 (m, 2H, Ar*H*), 7.23 – 7.26 (m, 2H, Ar*H*), 7.38 – 7.42 (m, 2H, Ar*H*). ^13^C NMR (101 MHz, DMSO) δ 165.40, 152.51, 141.37, 138.82, 138.23, 136.47, 128.99, 128.99, 128.39, 128.39, 126.56, 126.14, 126.14, 121.02, 121.02, 105.24, 54.71, 20.63, 17.06. HRMS (ESI-TOF (+)) calcd for [C_19_H_18_ClN_3_O_2_ [M+H]^+^, 356.1090; found 356.1165.

34. *N-(4-chlorophenyl)-4-(3,4-dichlorophenyl)-6-methyl-2-oxo-1,2,3,4-tetrahydropyrimidine-5-carboxamide* ***(CU5).***

Following the general **procedure C**, 3,4-dichlorobenzaldehyde (124.00 mg, 0.71 mmol, 1.00 eq), urea (63.80 mg, 1.06 mmol, 1.50 eq), 1.2, *N*-(4-chlorophenyl)-3-oxobutanamide (150.00 mg, 0.71 mmol, 1.00 eq), Cu(OTf)_2_ (7.20 mg, 0.02 mmol, 0.02 eq) in 1.40 mL of EtOH. Column chromatography (93.00 % DCM / 7.00 % MeOH) gave comp **CU5** (112.00 mg, 20.00 % yield) as a white solid, mp 258 ºC. IR: 3266, 3113, 1711, 1668, 1593, 1512, 1493, 1467, 1397, 1312, 1245, 1133, 1089, 1029, 1013, 820, 755, 707, 683, 640 cm^-1^. ^1^H NMR (400 MHz, MeOD) δ 2.09 (d, *J* = 1.0 Hz, 3H, C*H*_3_), 5.44 (d, *J* = 1.2 Hz, 1H, C*H*), 7.24 – 7.29 (m, 3H, Ar*H*), 7.42 – 7.46 (m, 2H, Ar*H*), 7.47 – 7.50 (m, 2H, Ar*H*). ^13^C NMR (101 MHz, DMSO) δ 165.14, 152.24, 145.16, 139.85, 138.00, 131.01, 130.83, 129.93, 128.46, 128.46, 128.34, 126.80, 126.69, 121.15, 121.15, 104.14, 54.08, 17.17. HRMS (ESI-TOF (+)) calcd for [C_18_H_14_Cl_3_N_3_O_2_ +H]^+^, 410.0150; found 410.0238.

**CHROMATOGRAPHY AND NMR SPECTRA**

HPLC trace for blank:

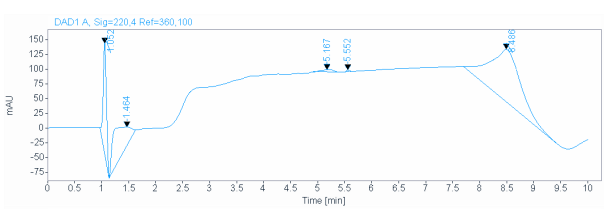

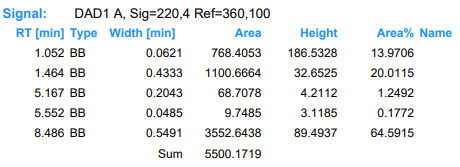

HPLC trace for compound **U1:**

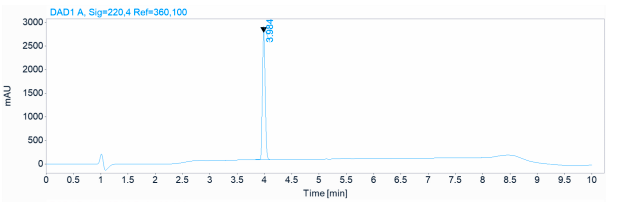

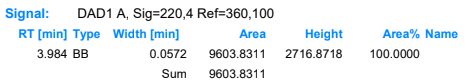

HPLC trace for compound **U2:**

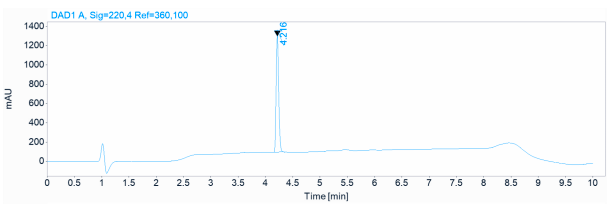

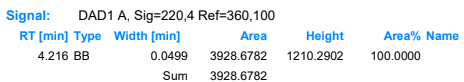

HPLC trace for compound **U3:**

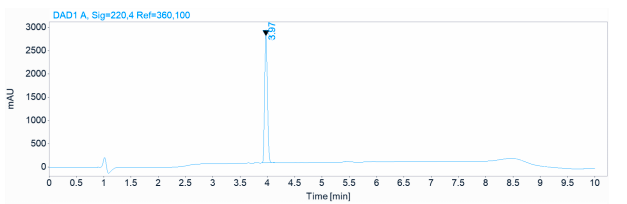

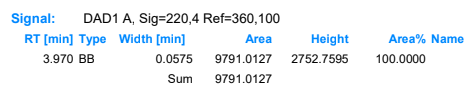

HPLC trace for compound **U4:**

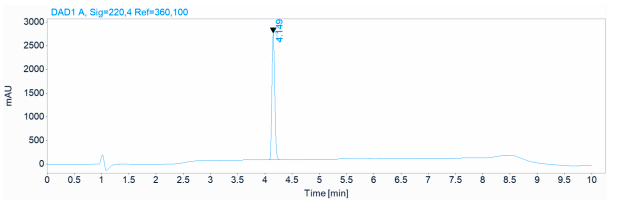

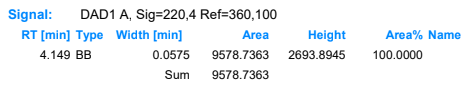

HPLC trace for compound **U5:**

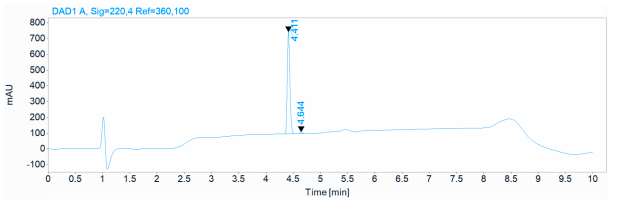

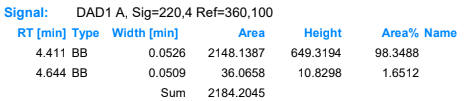

HPLC trace for compound **U6:**

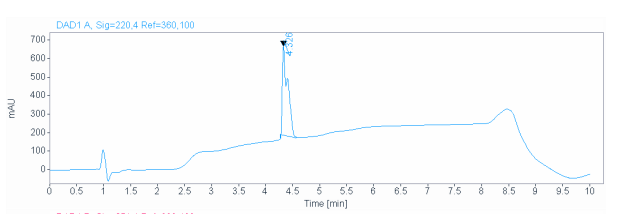

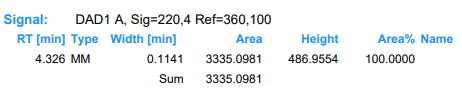

HPLC trace for compound **U7:**

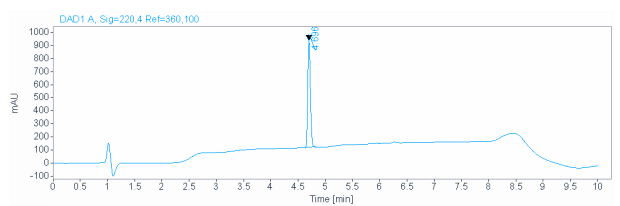

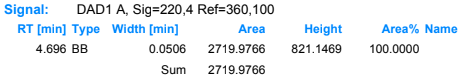

HPLC trace for compound **T1:**

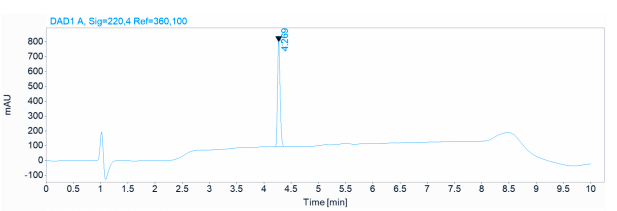

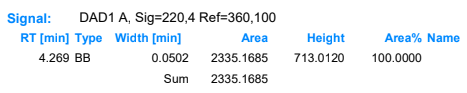

HPLC trace for compound **T2:**

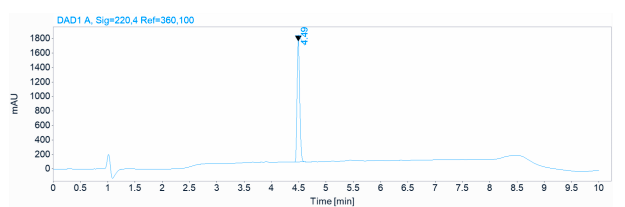

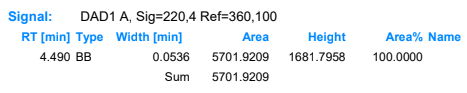

HPLC trace for compound **T3:**

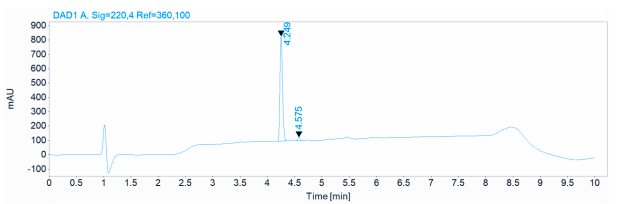

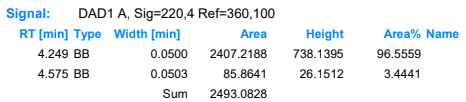

HPLC trace for compound **T4:**

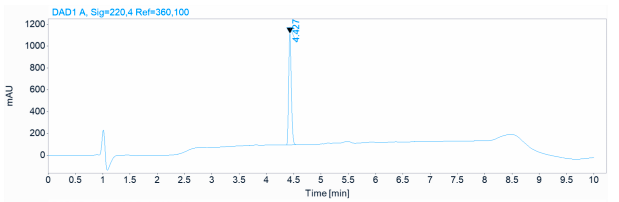

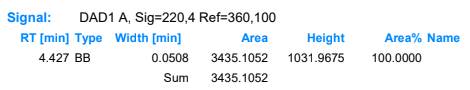

HPLC trace for compound **T5:**

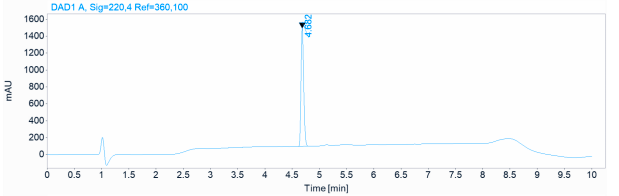

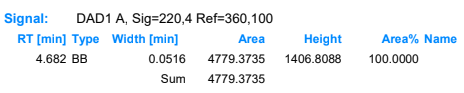

HPLC trace for compound **T6:**

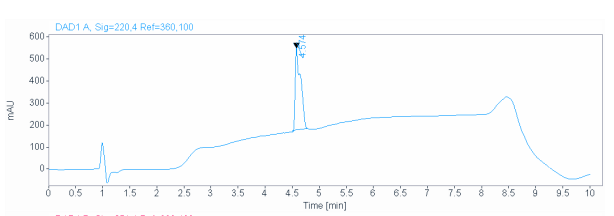

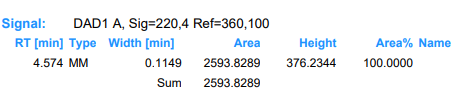

HPLC trace for compound **CU1:**

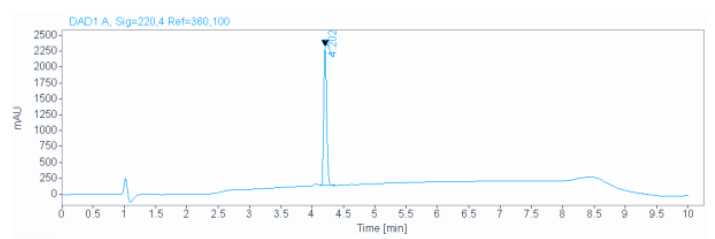

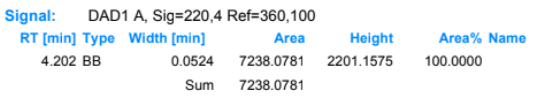

HPLC trace for compound **CU2:**

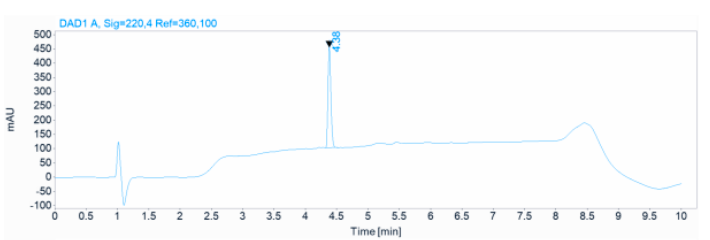

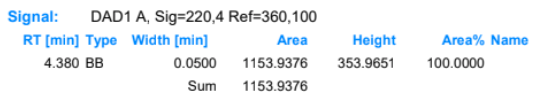

HPLC trace for compound **CU3:**

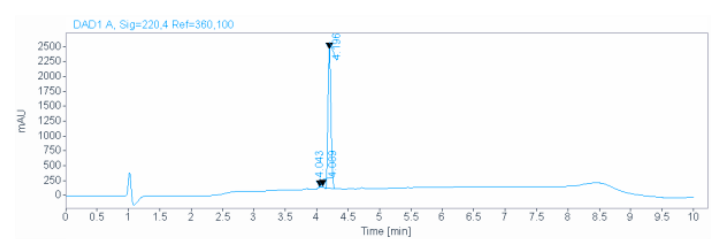

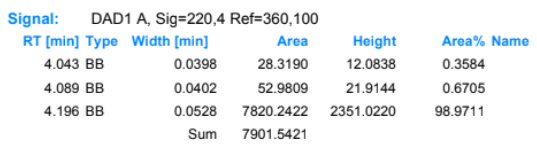

HPLC trace for compound **CU4:**

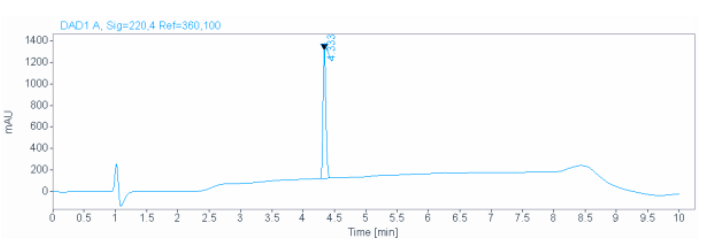

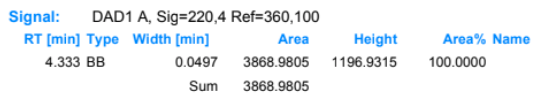

HPLC trace for compound **CU5:**

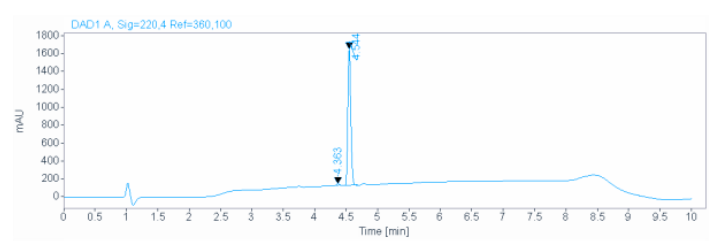

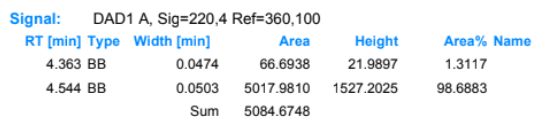
HPLC trace for compound **U8:**

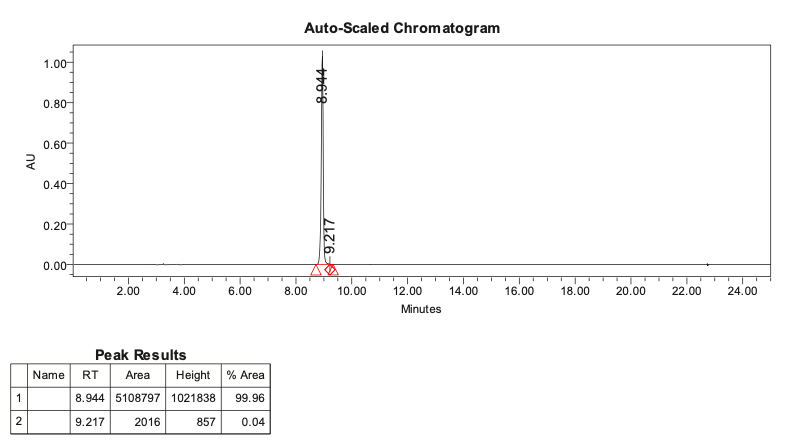

HPLC trace for compound **U9:**

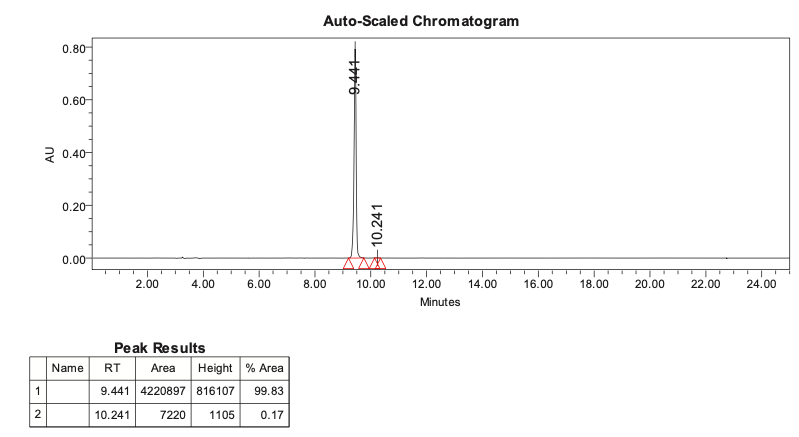

HPLC trace for compound **U10:**

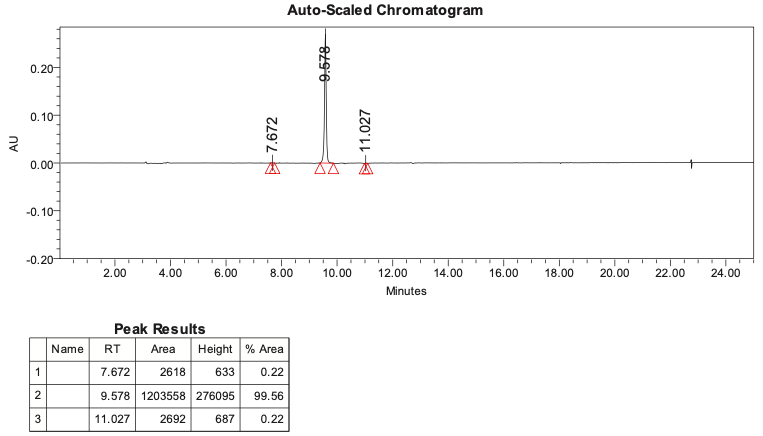

HPLC trace for compound **U11:**

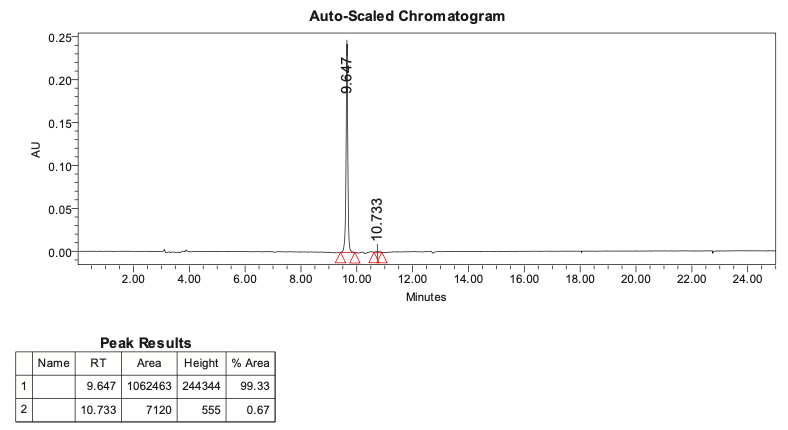

HPLC trace for compound **T9:**

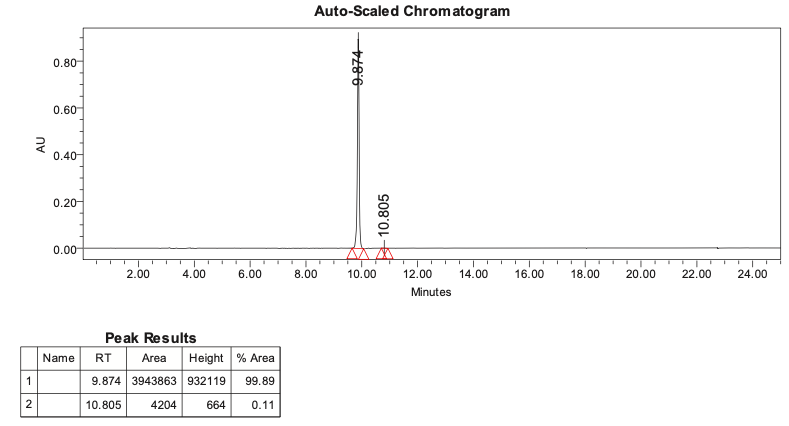

HPLC trace for compound **T10:**

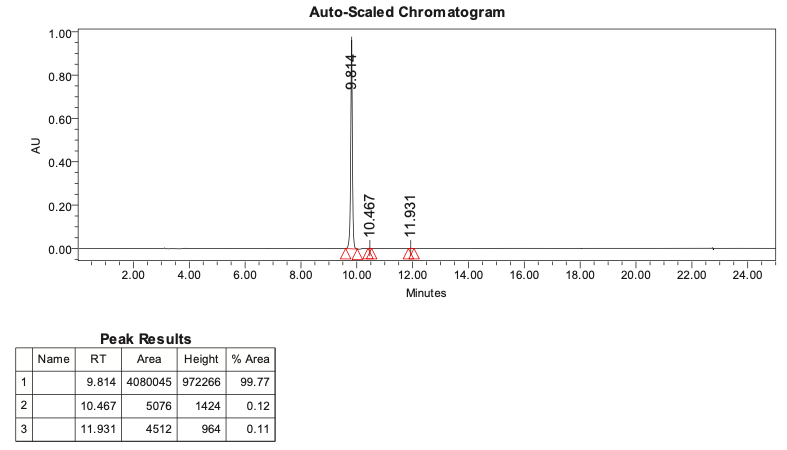

HPLC trace for compound **T11:**

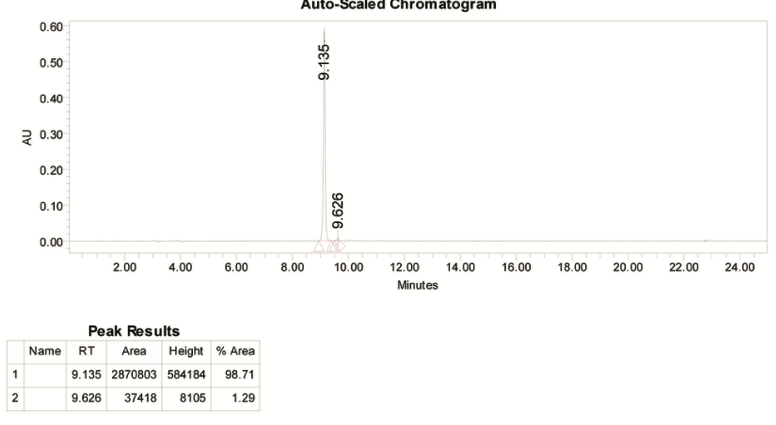

HPLC trace for compound **U12:**

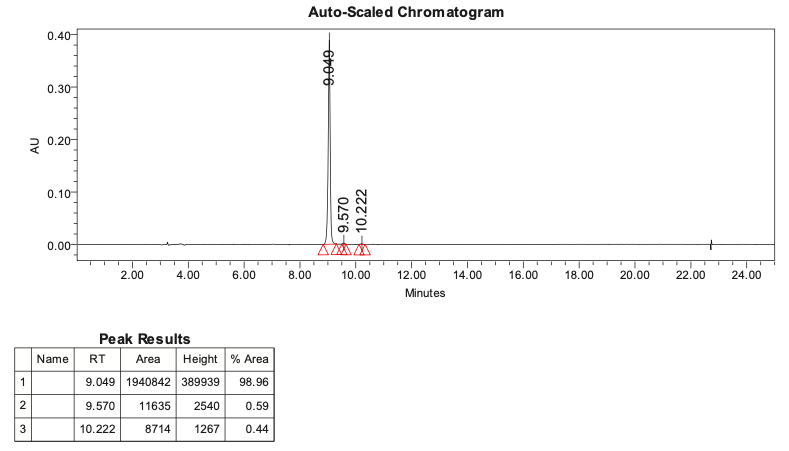

HPLC trace for compound **U13:**

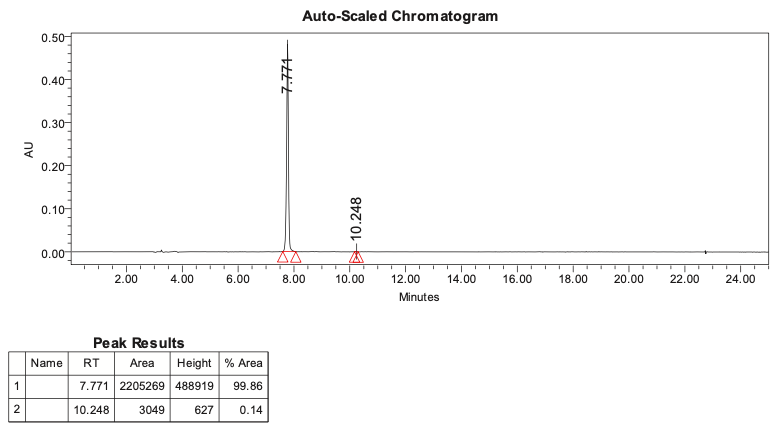

HPLC trace for compound **T12:**

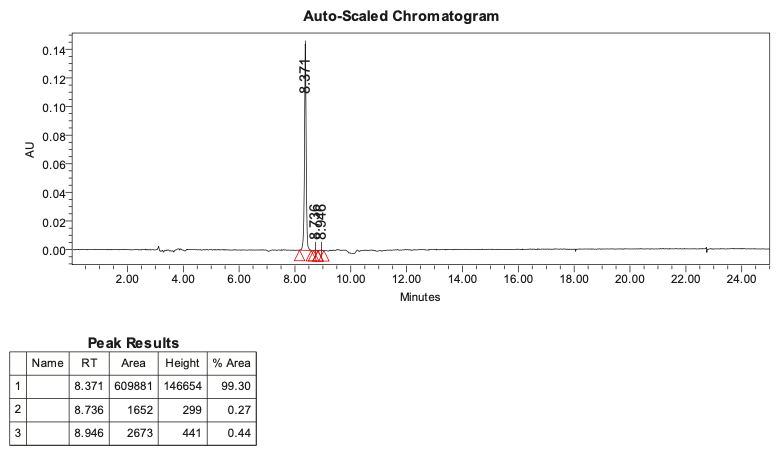

HPLC trace for compound **U14:**

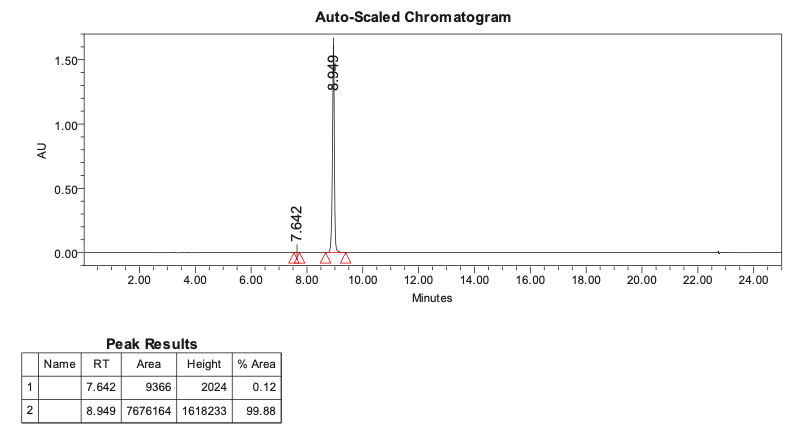

HPLC trace for compound **T13:**

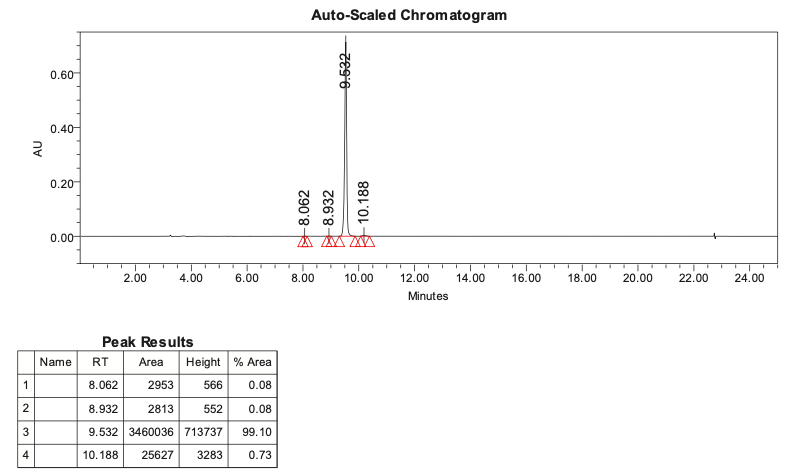

HPLC trace for compound **T14:**

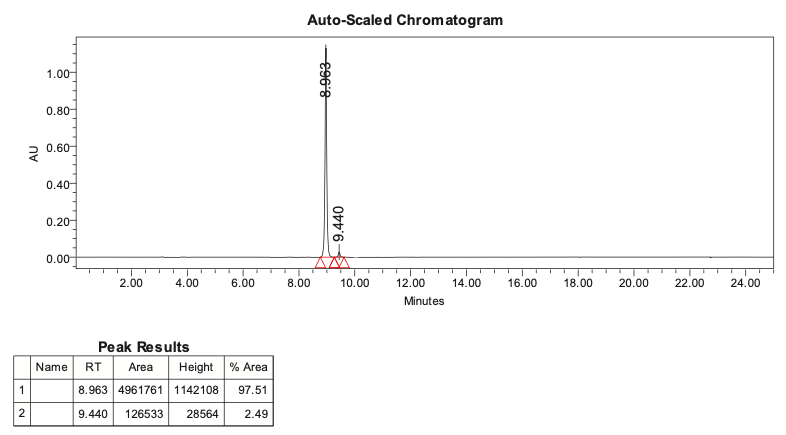

HPLC trace for compound **U15:**

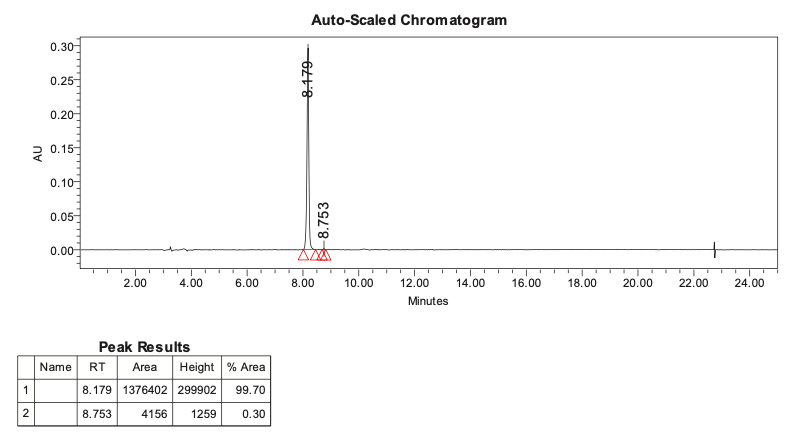

**^1^H-NMR AND ^13^C-NMR SPECTRA**

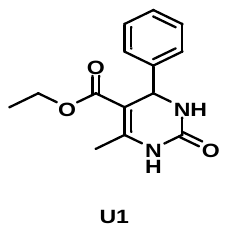

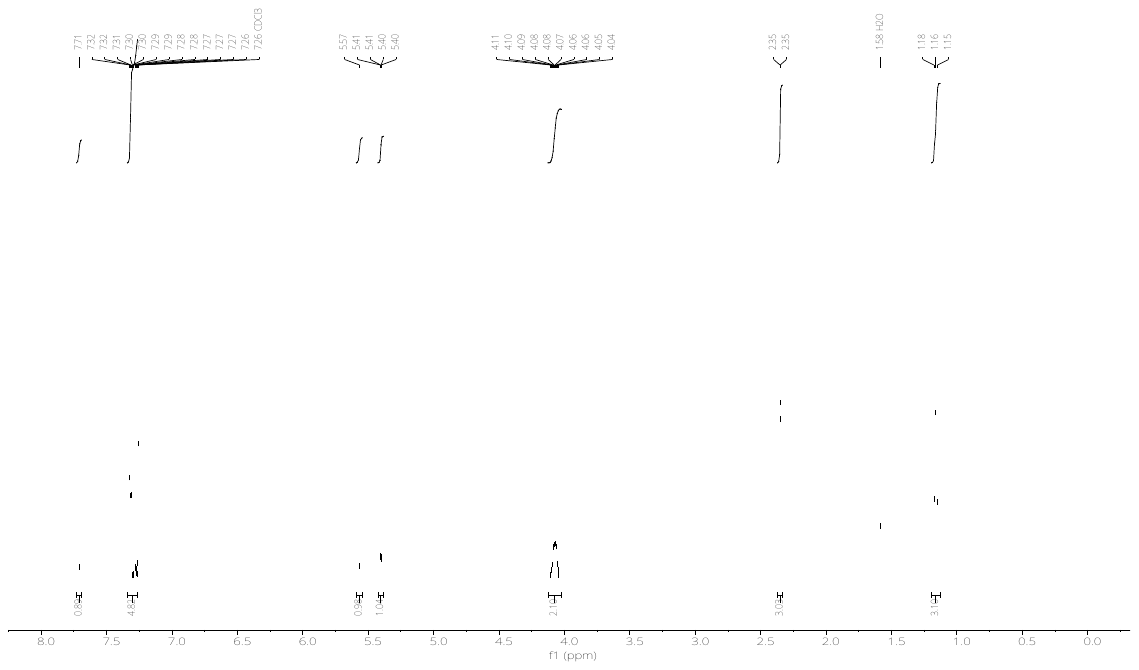

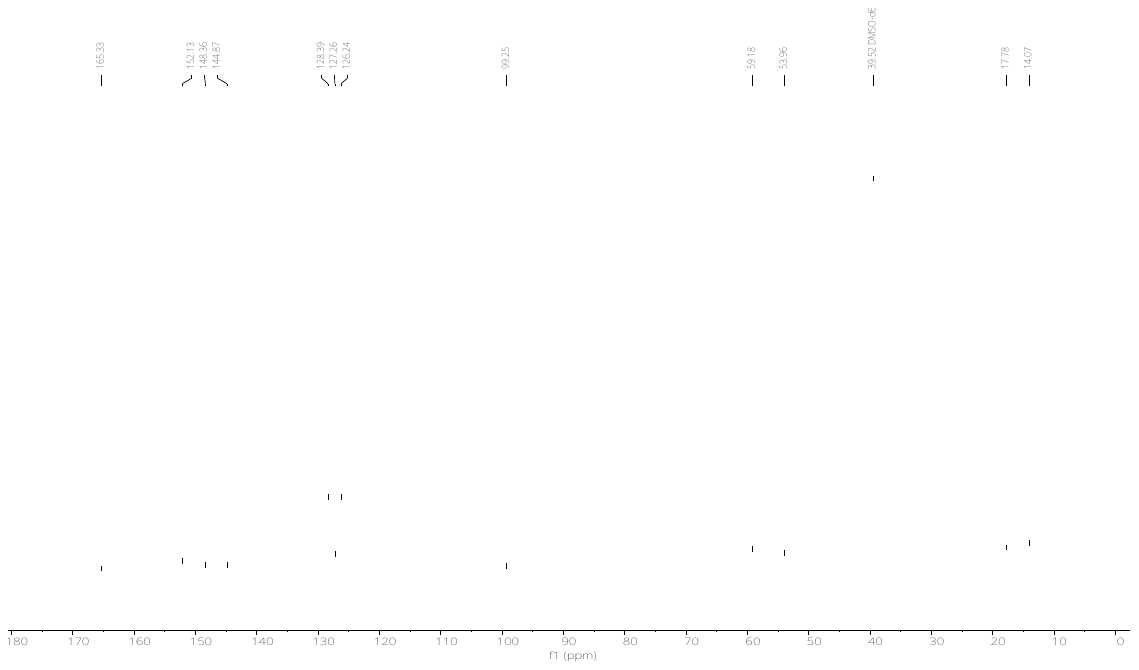

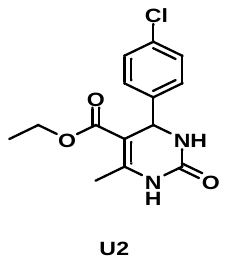

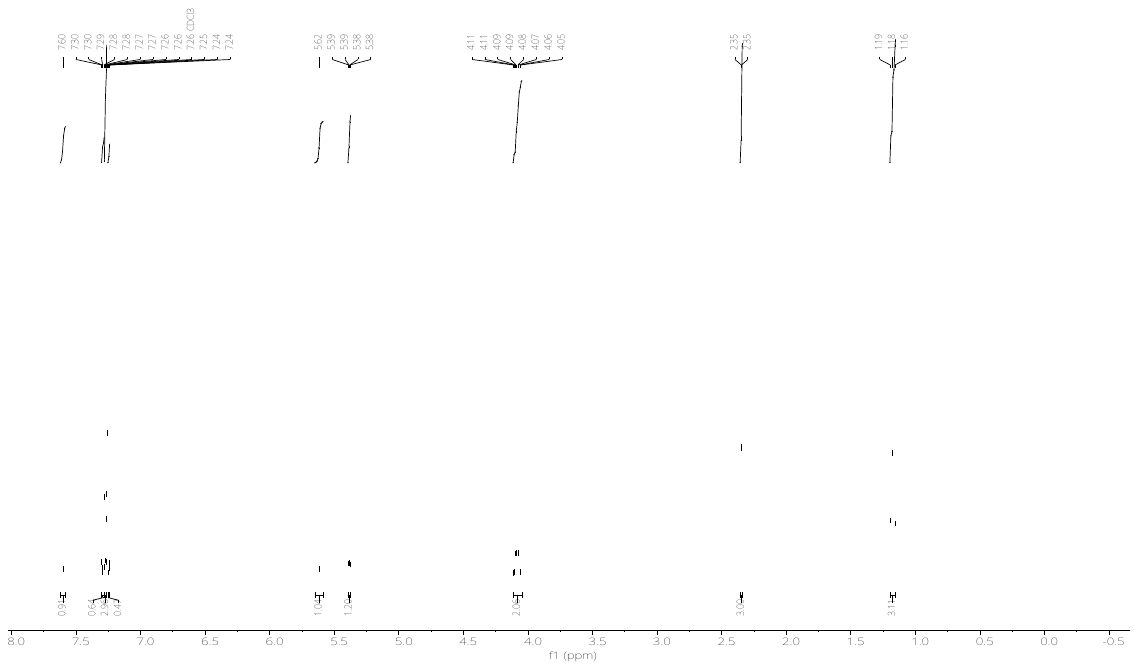

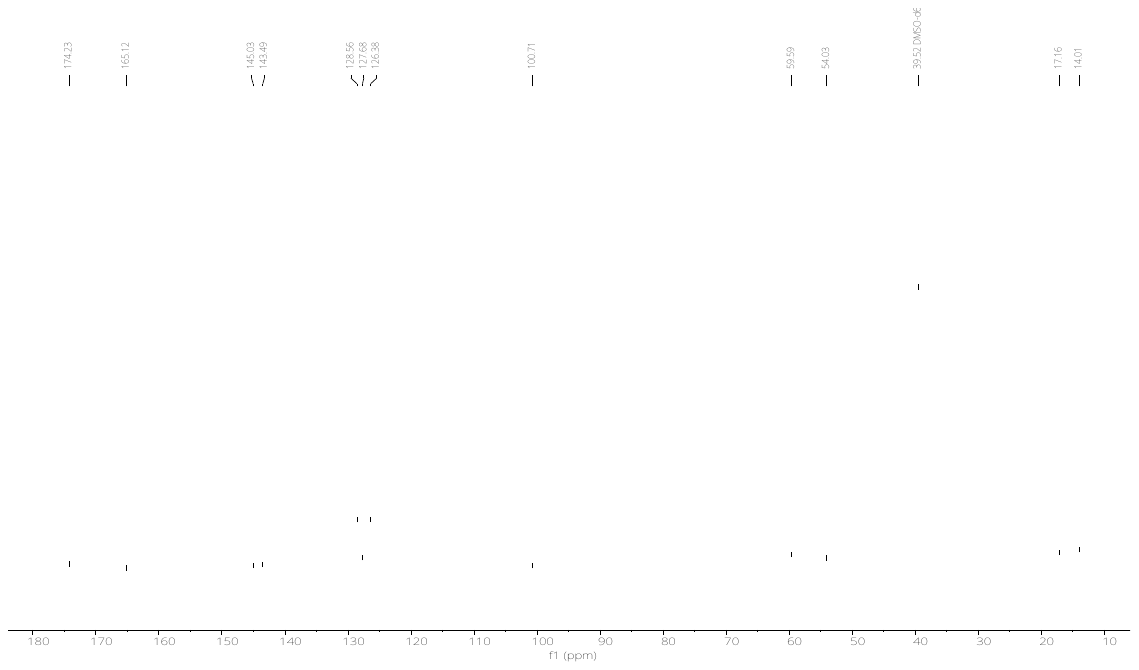

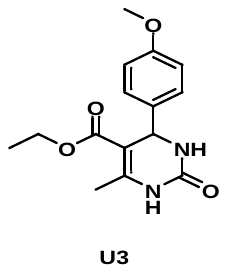

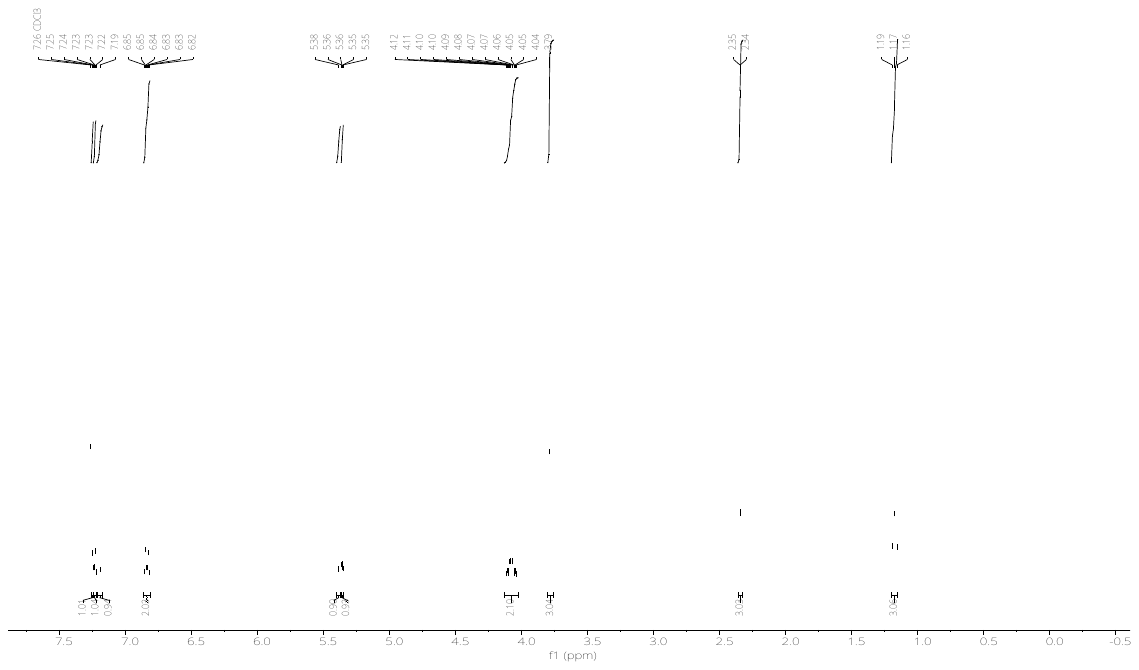

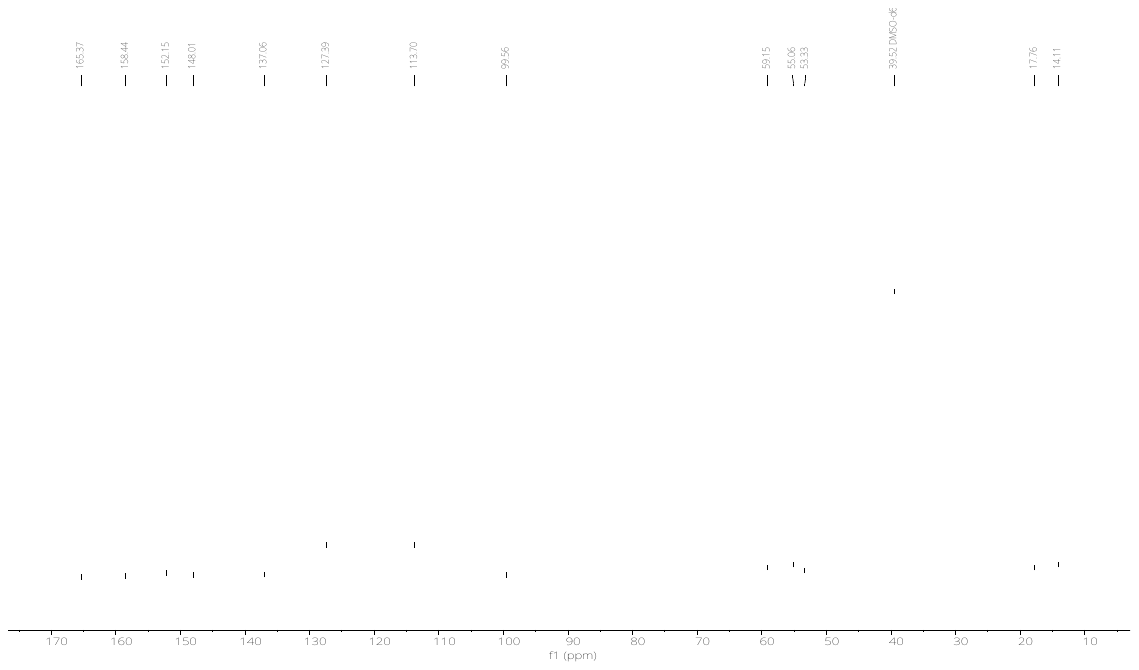

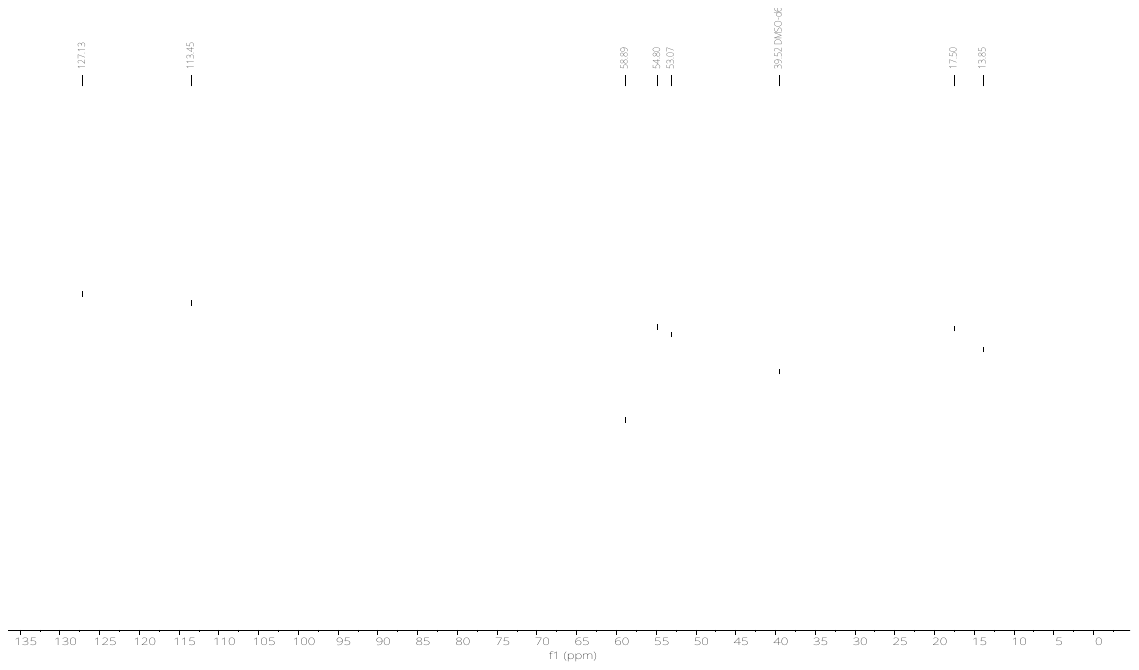

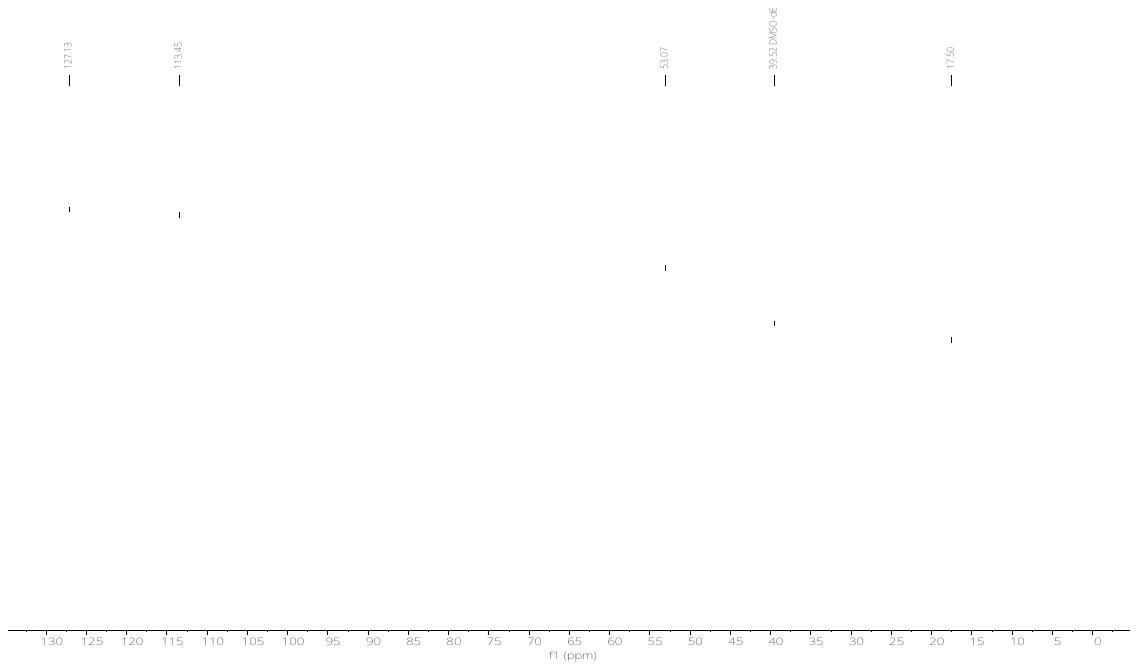

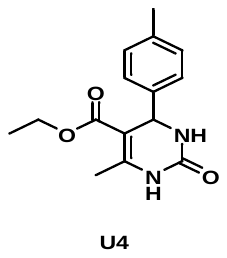

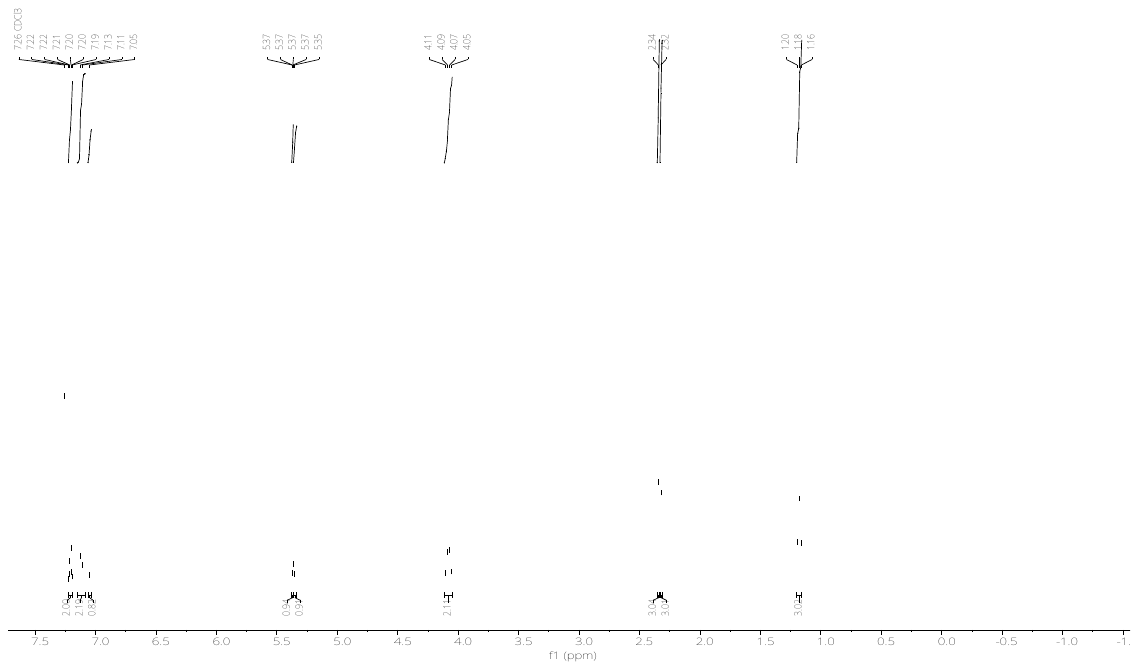

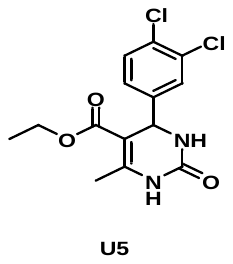

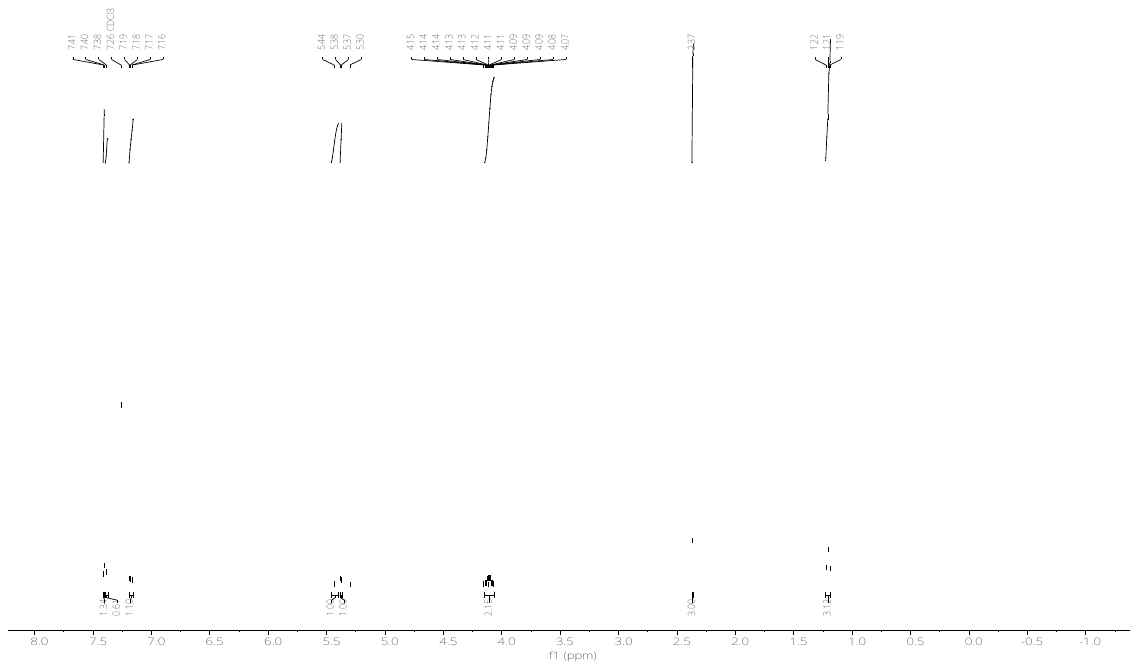

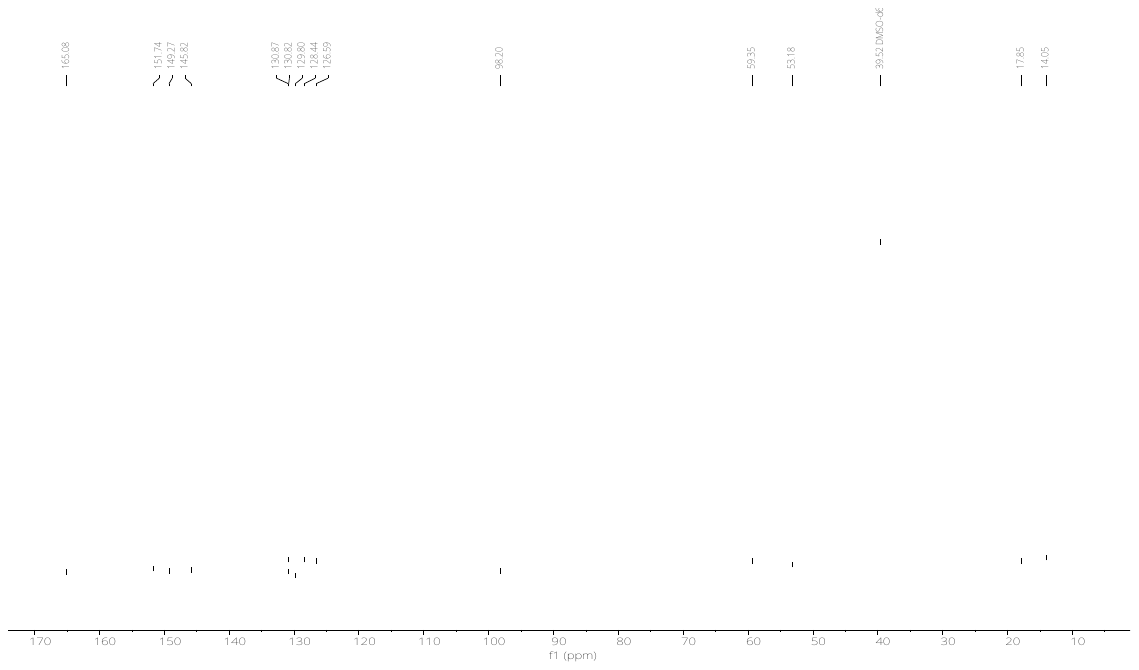

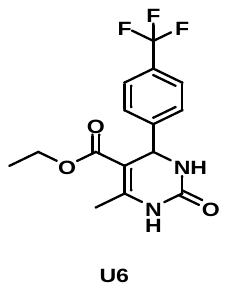

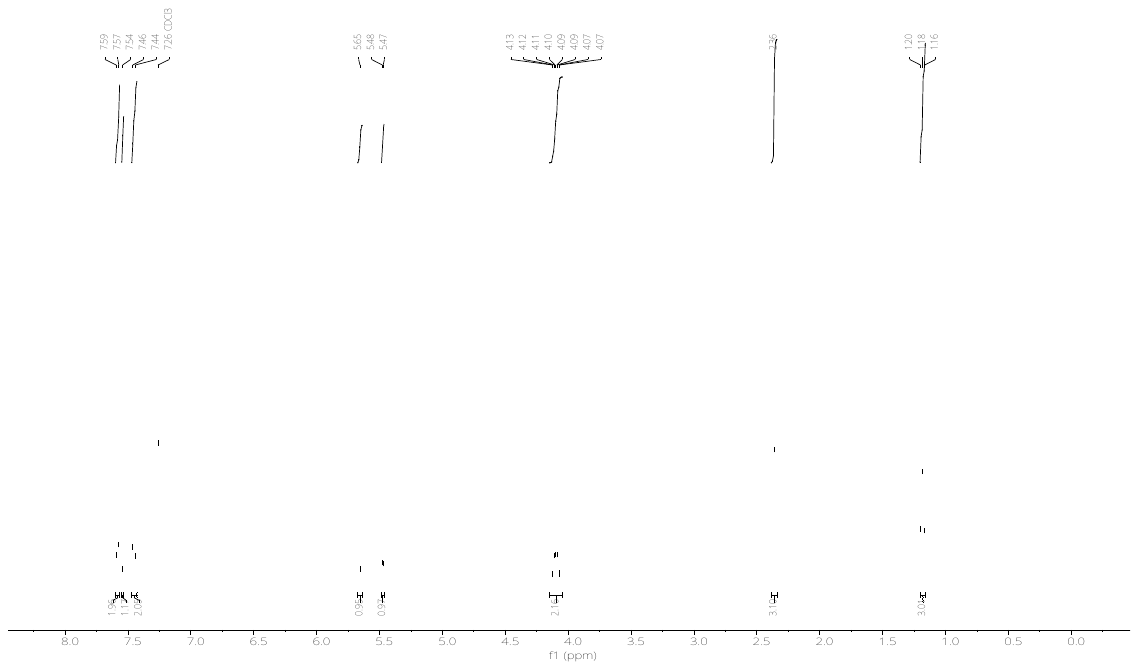

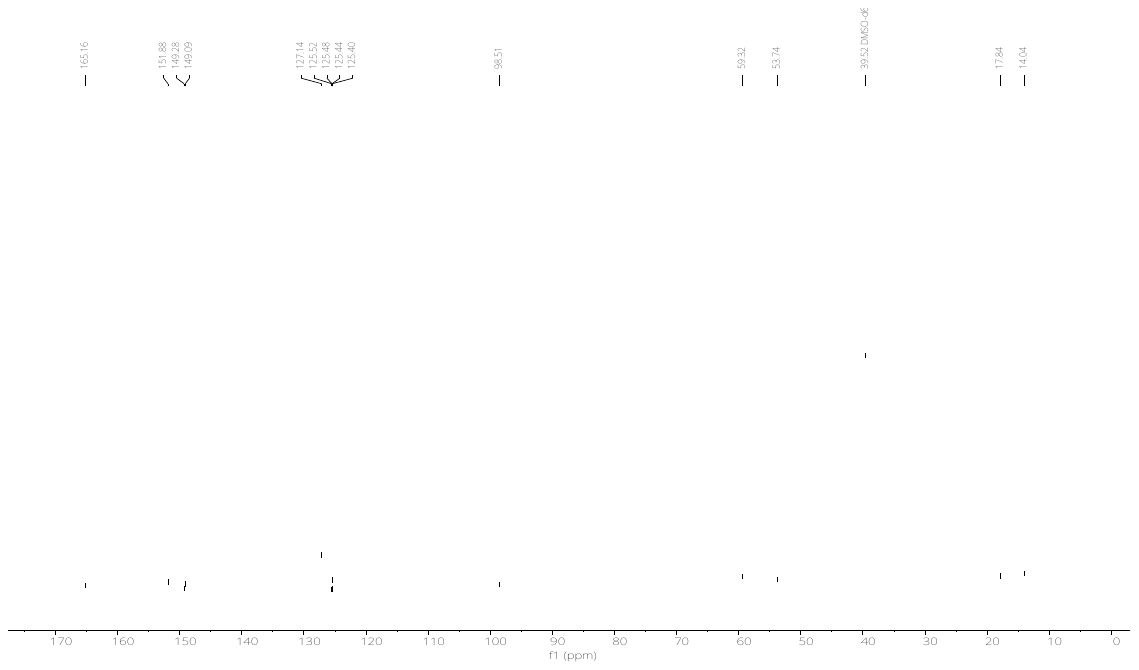

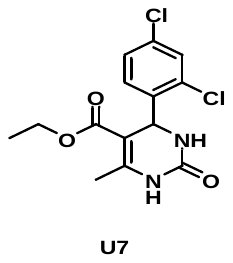

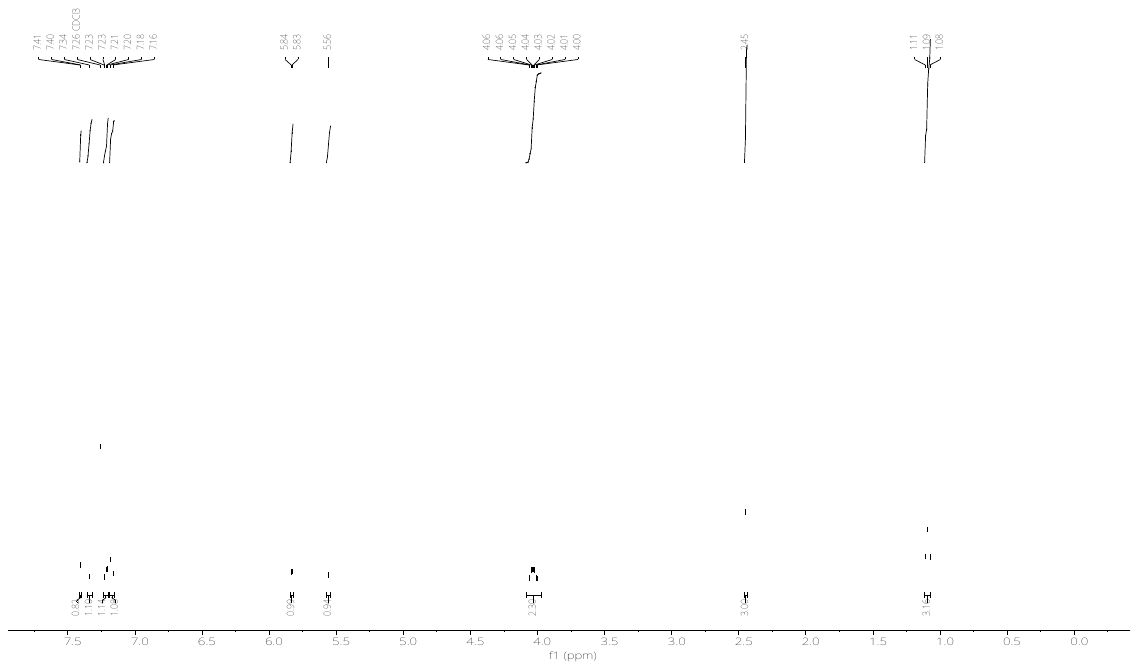

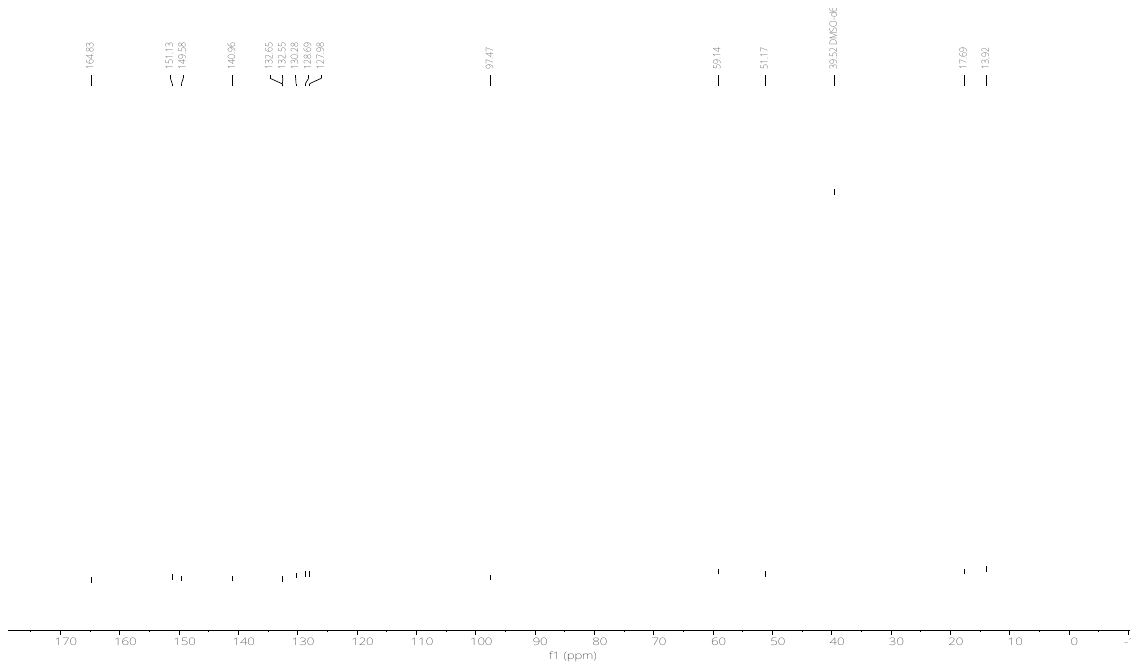

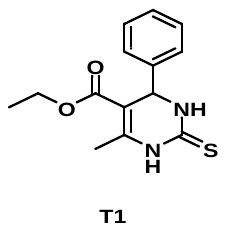

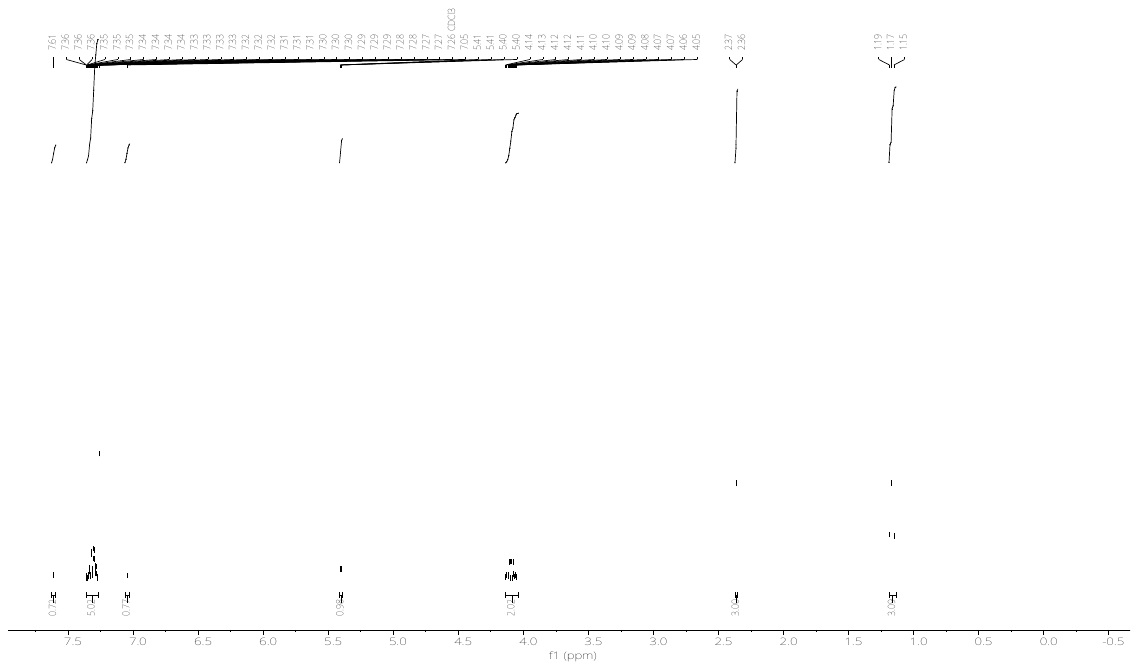

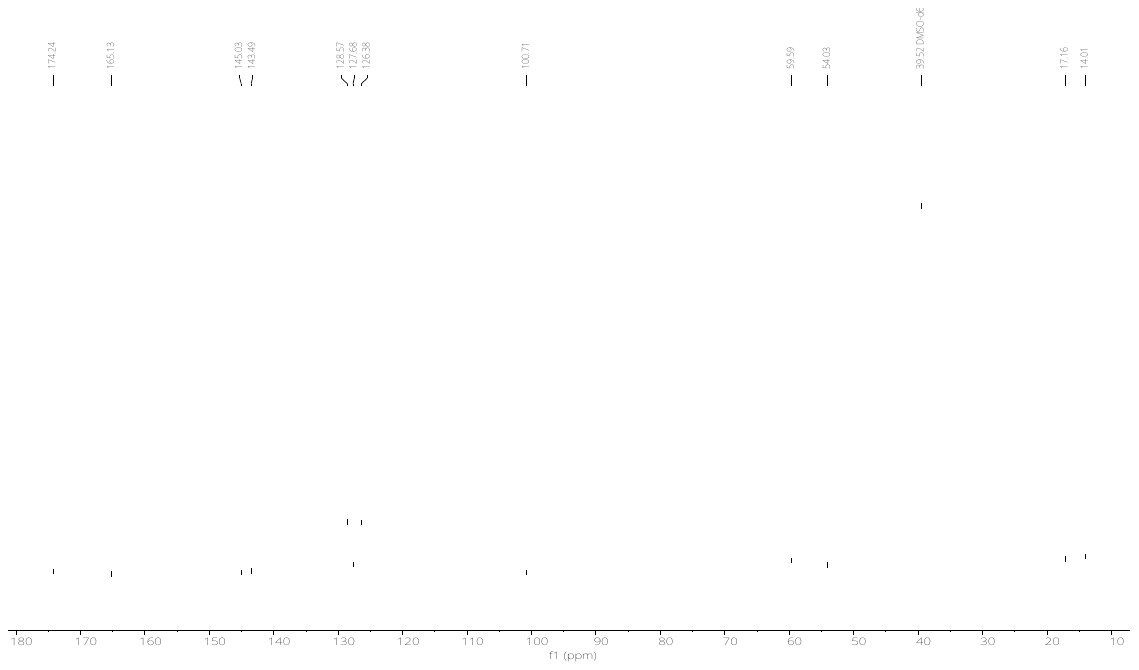

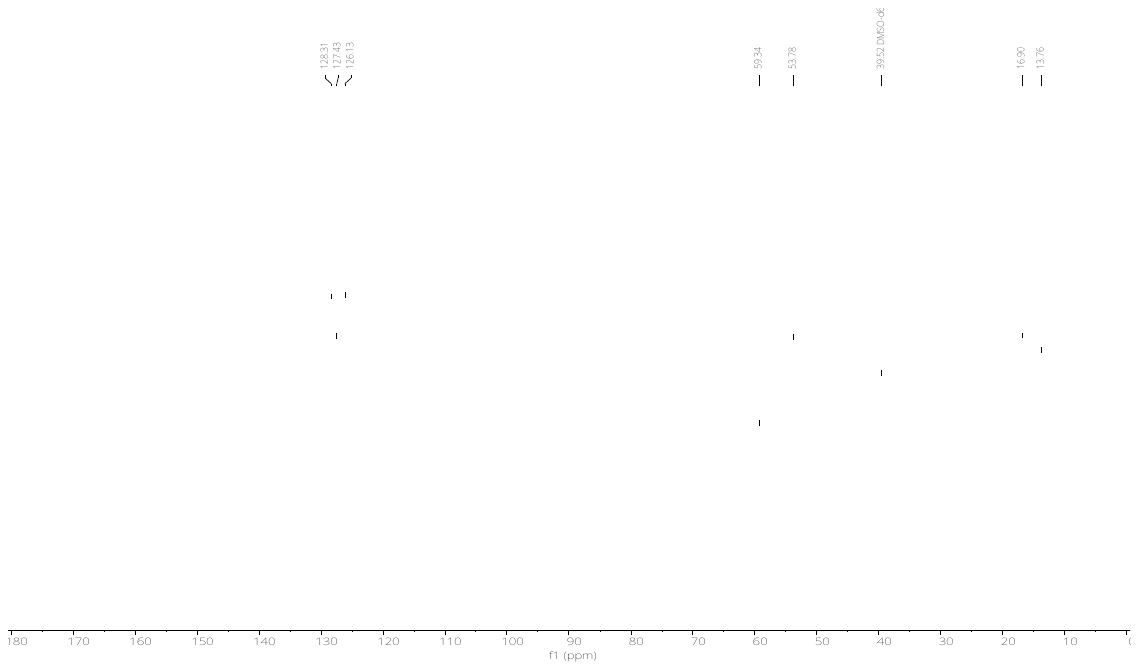

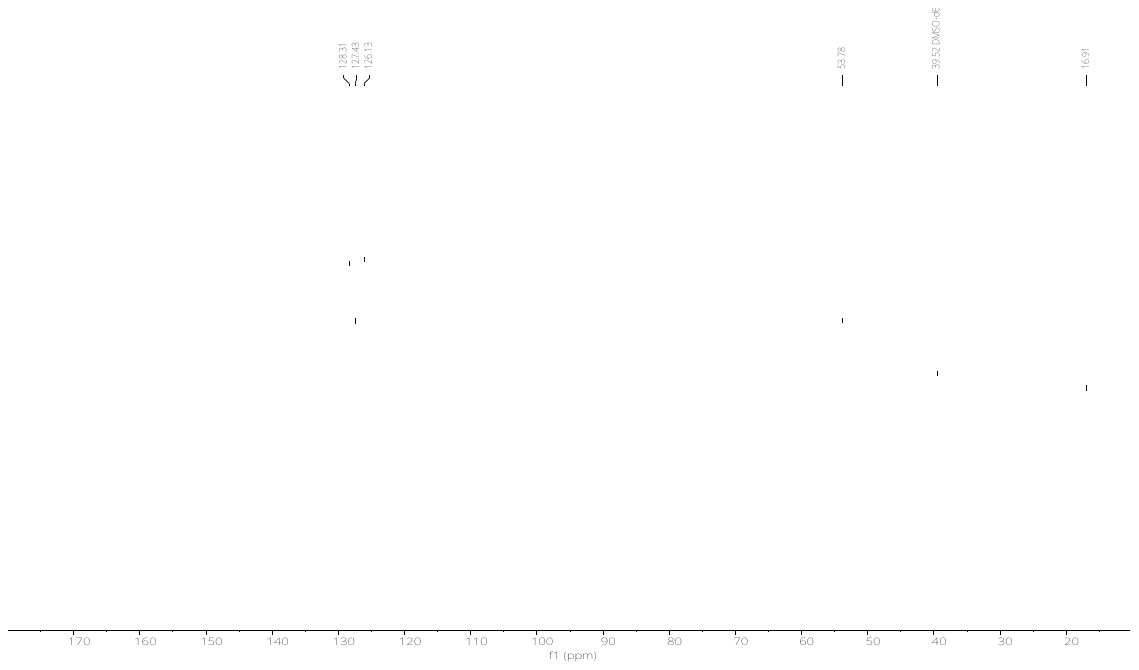

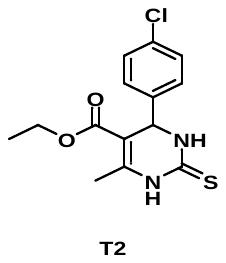

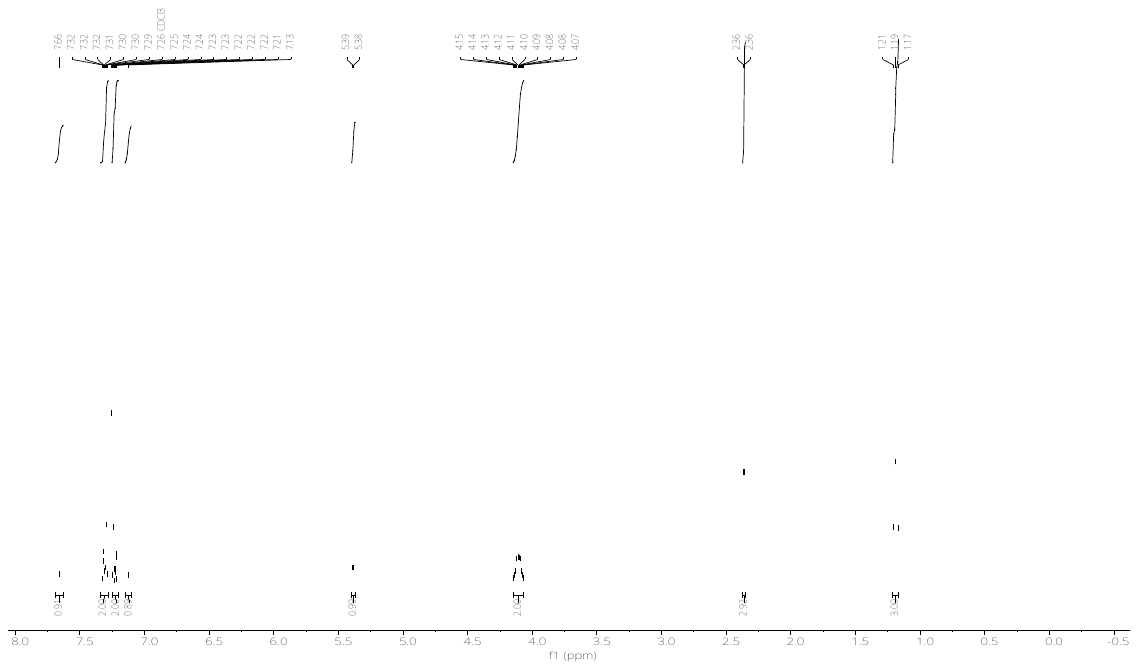

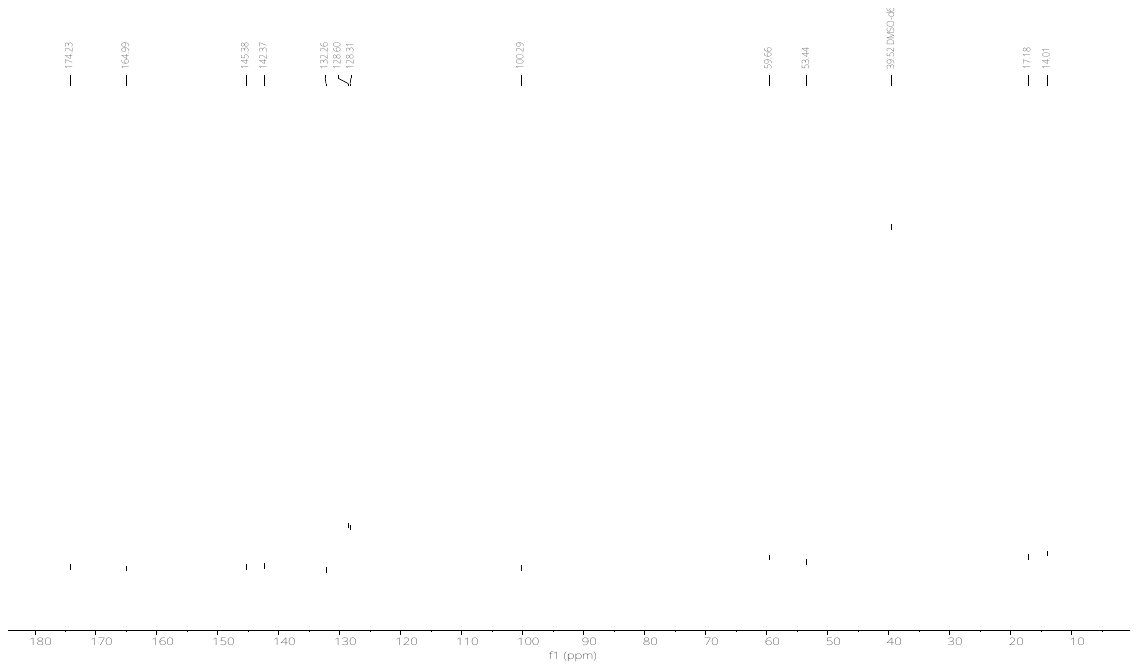

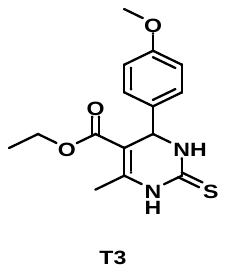

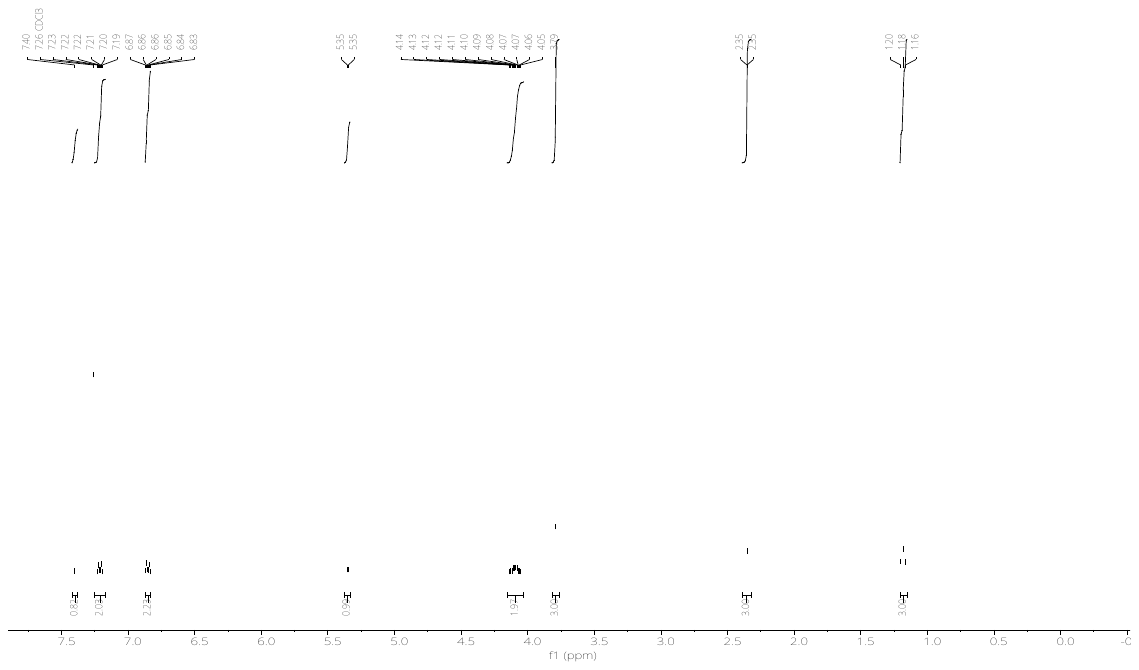

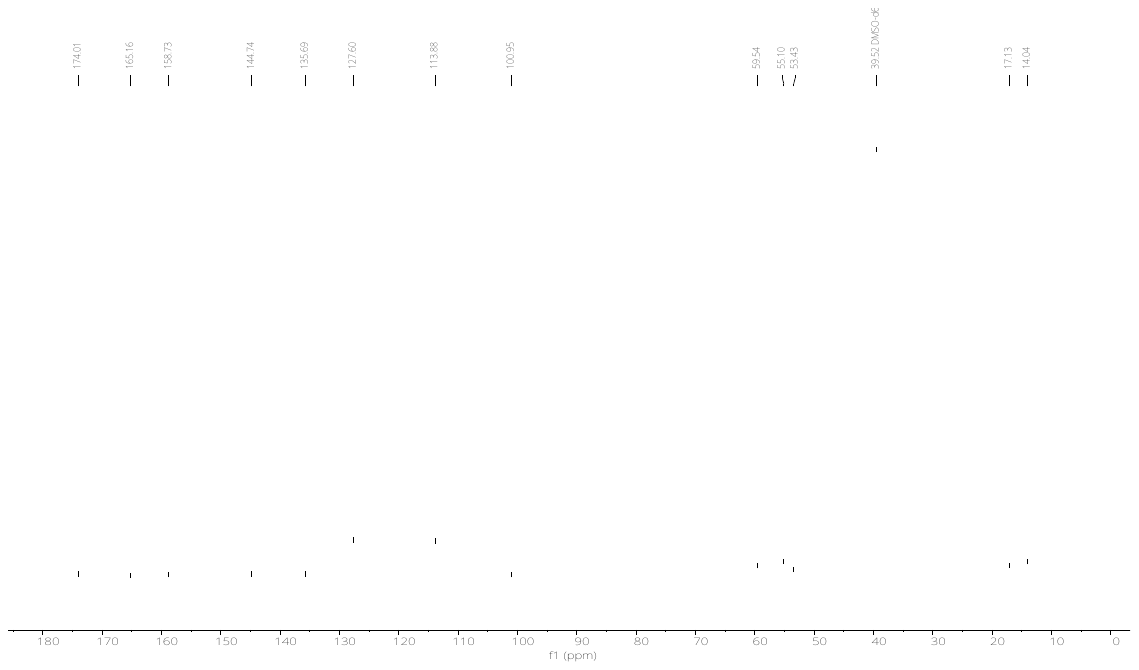

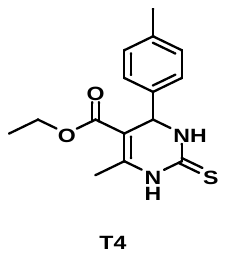

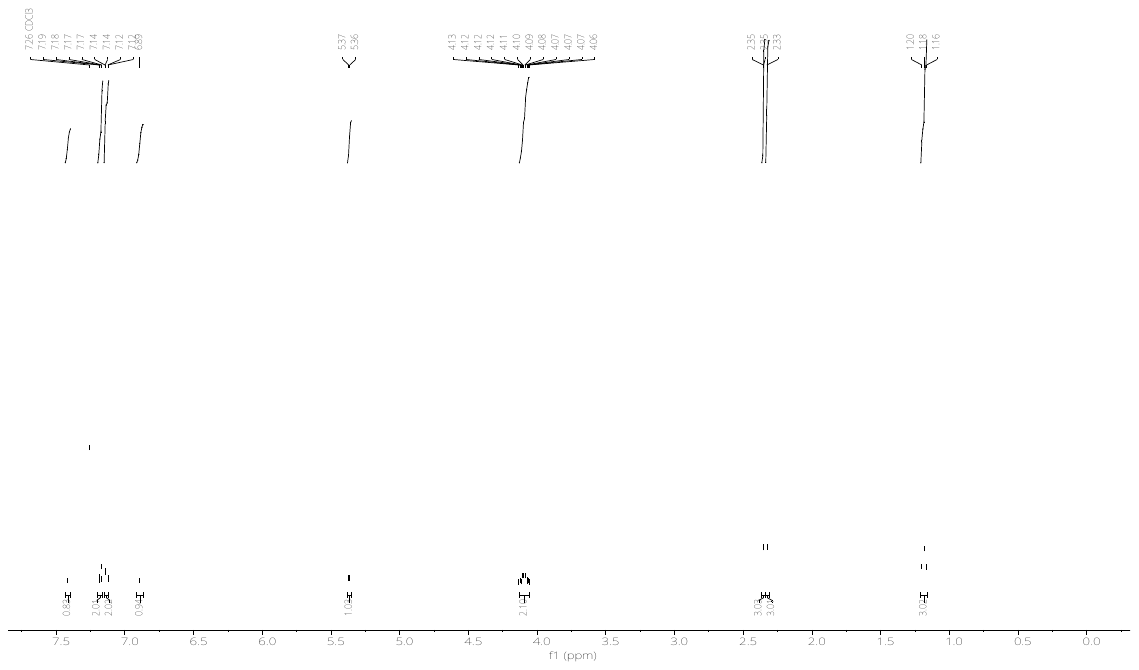

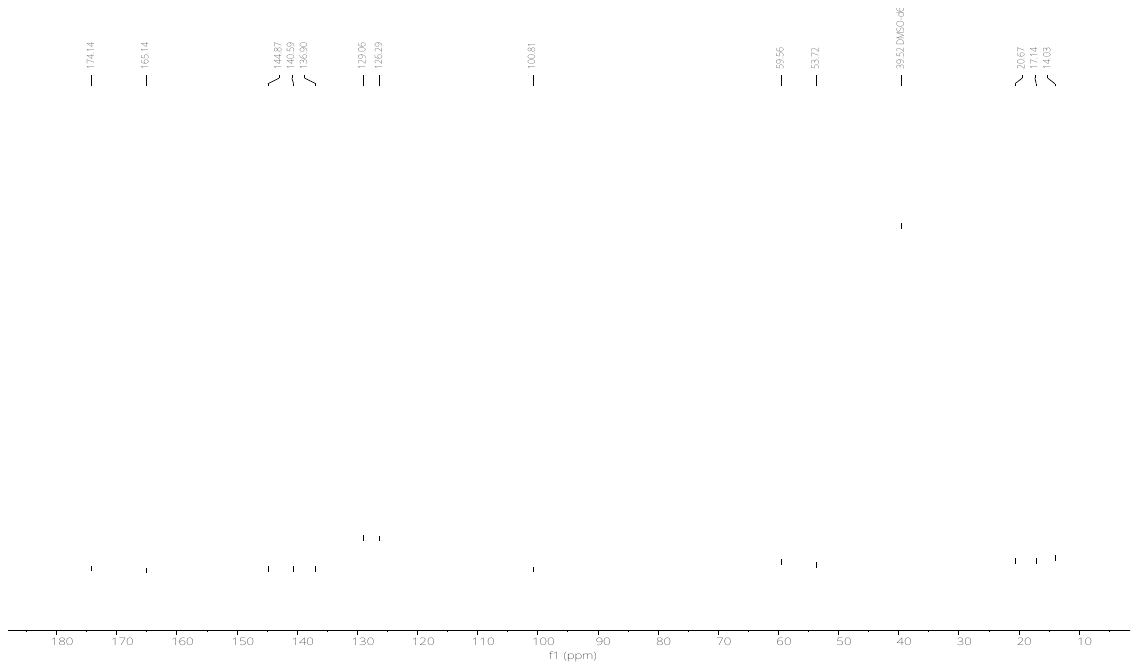

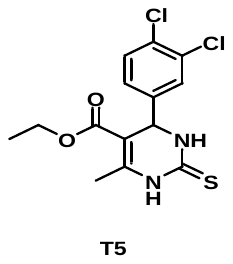

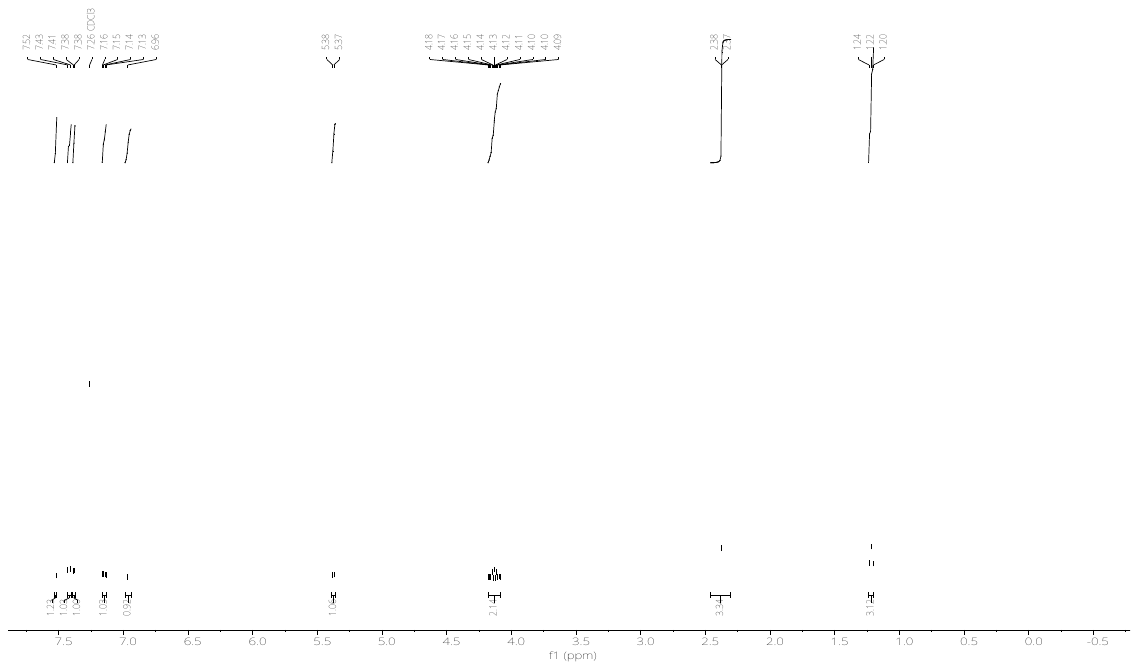

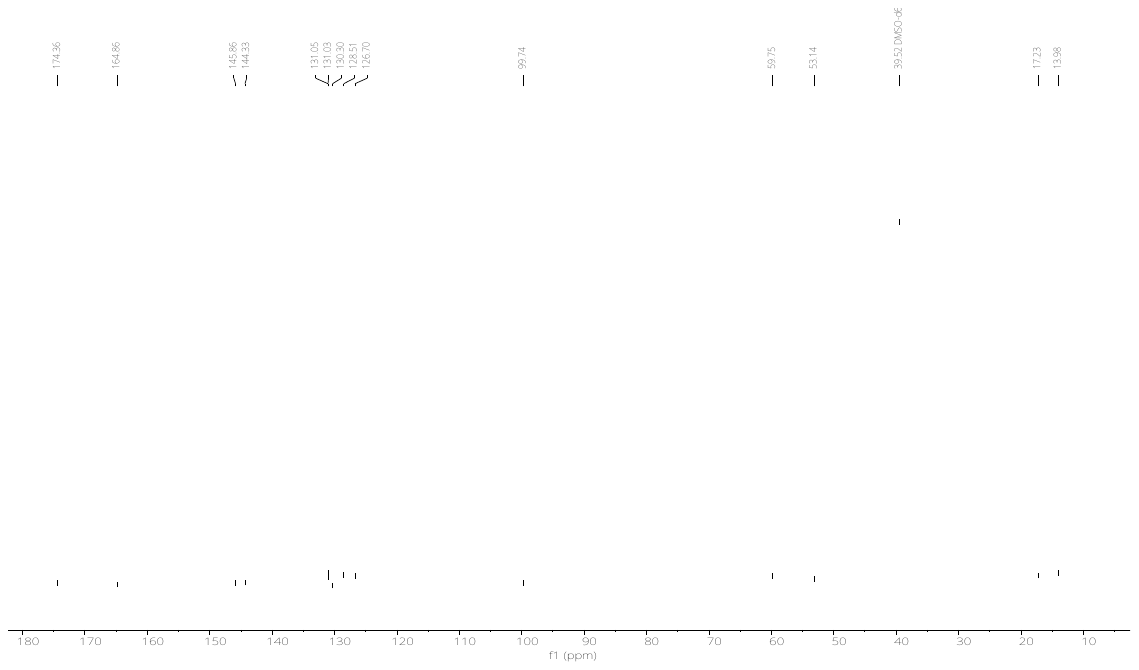

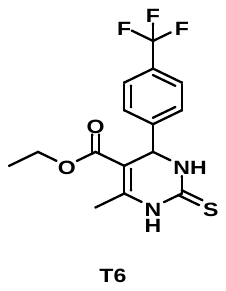

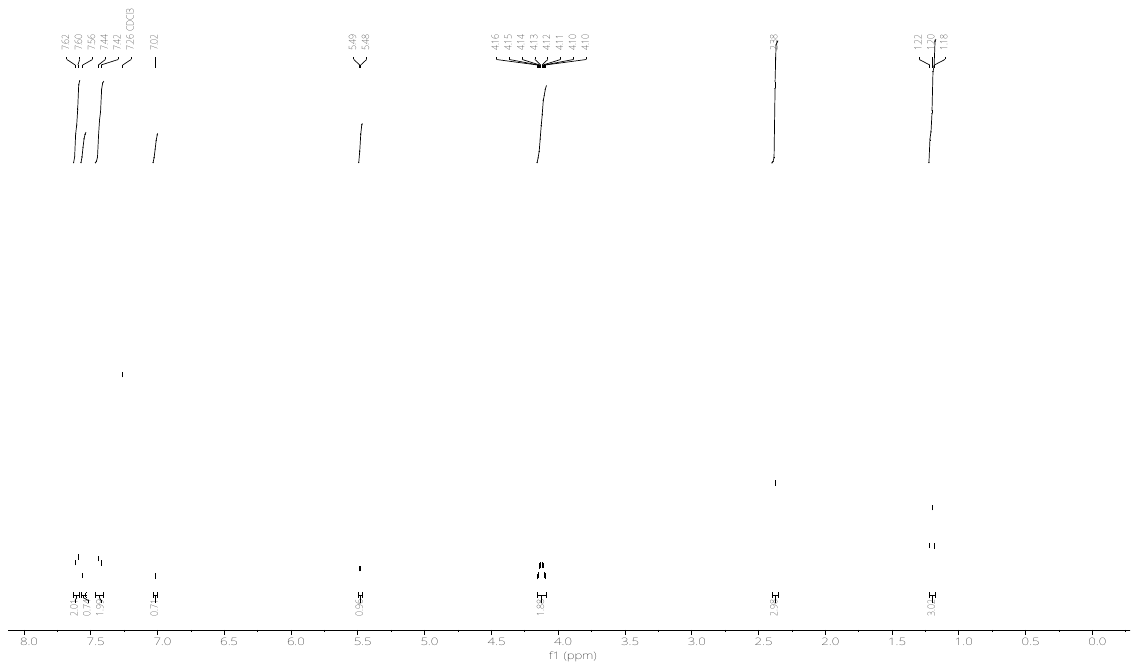

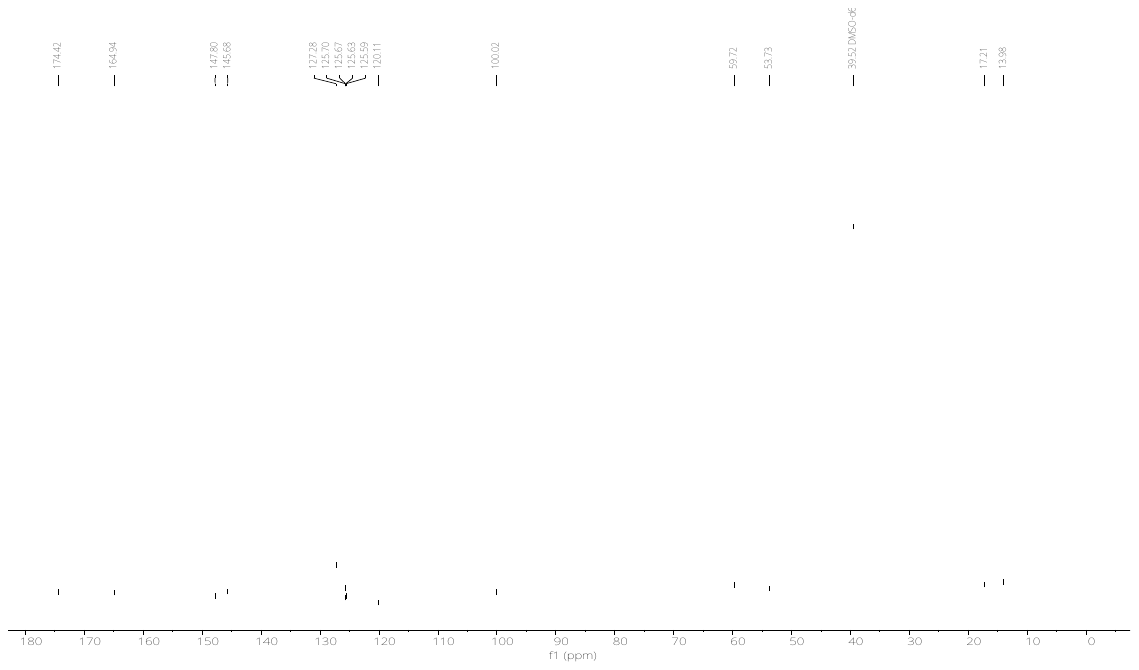

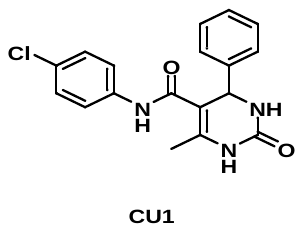

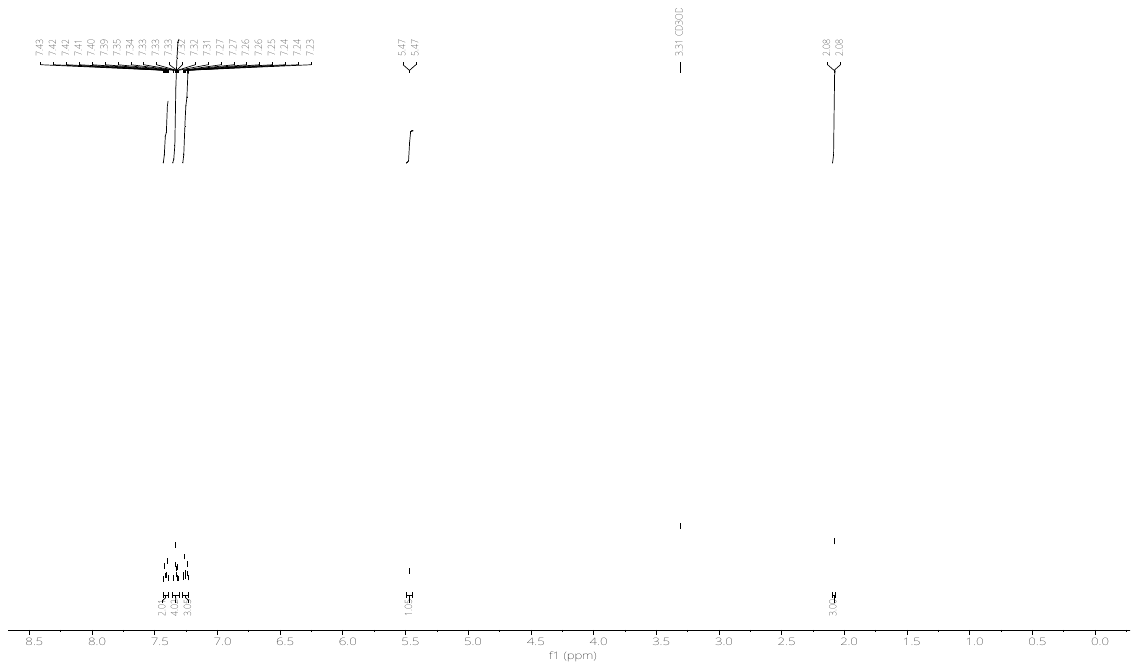

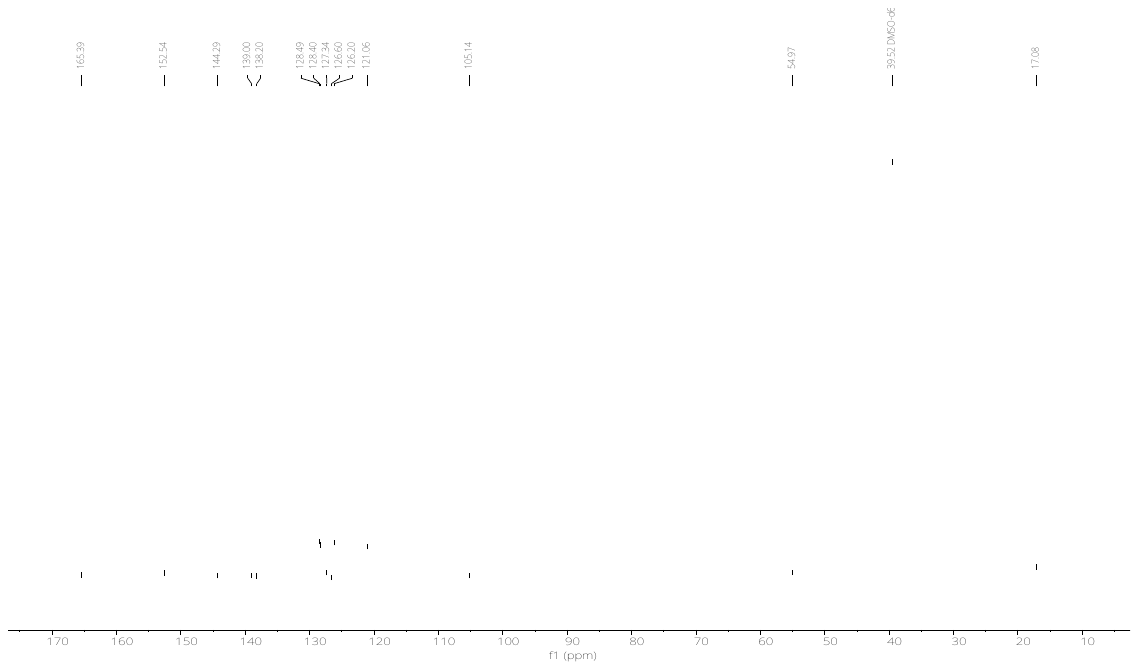

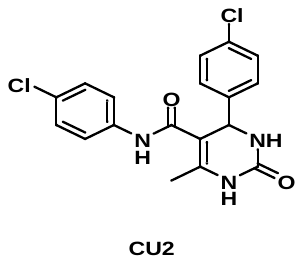

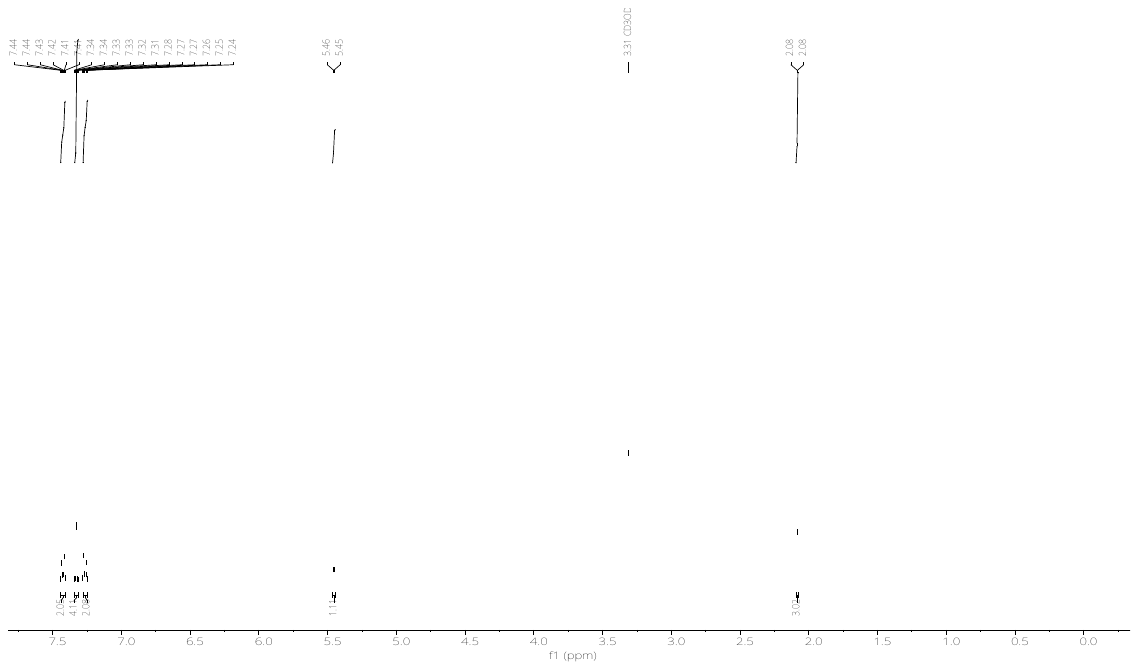

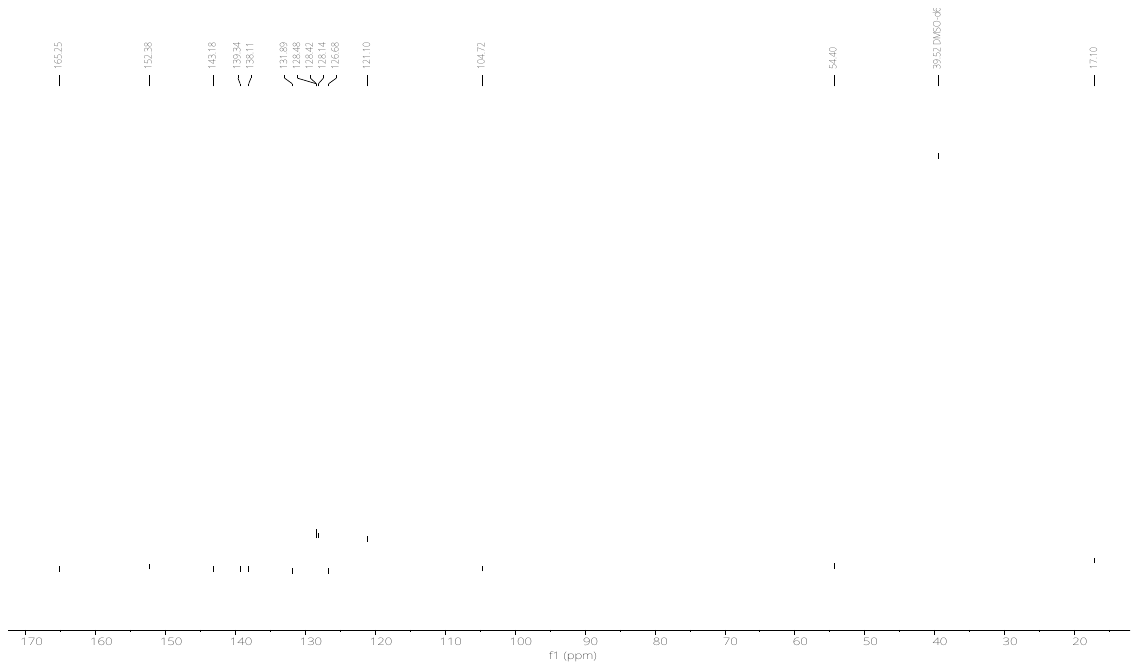

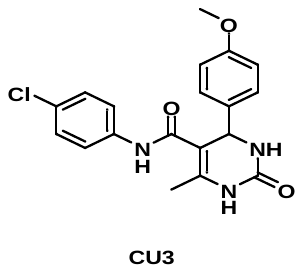

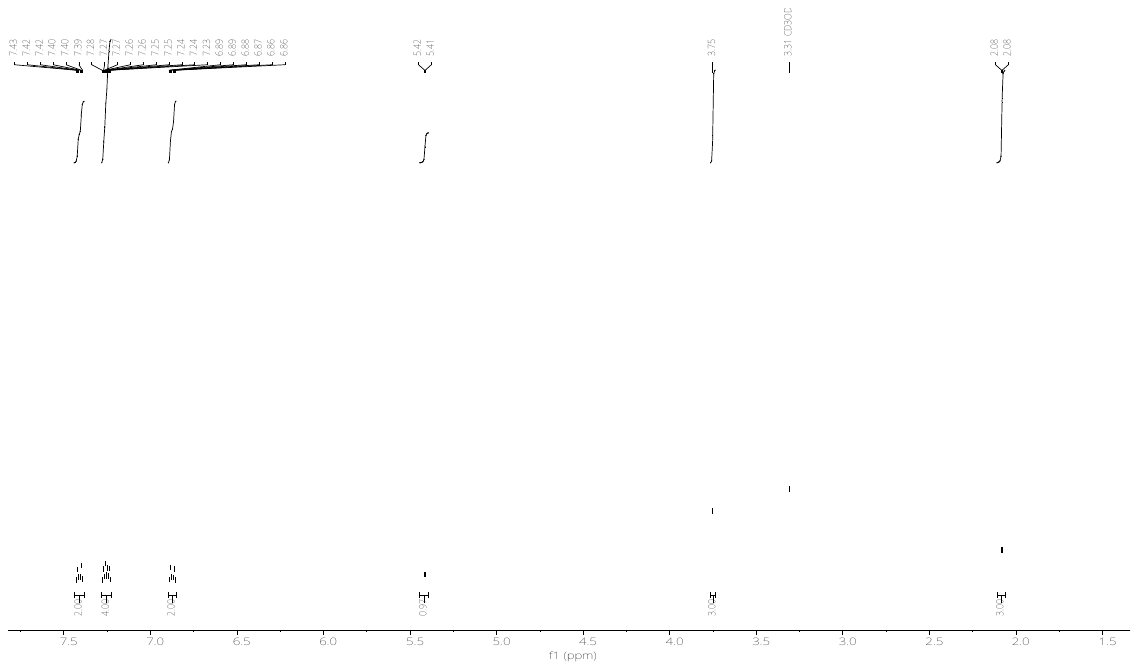

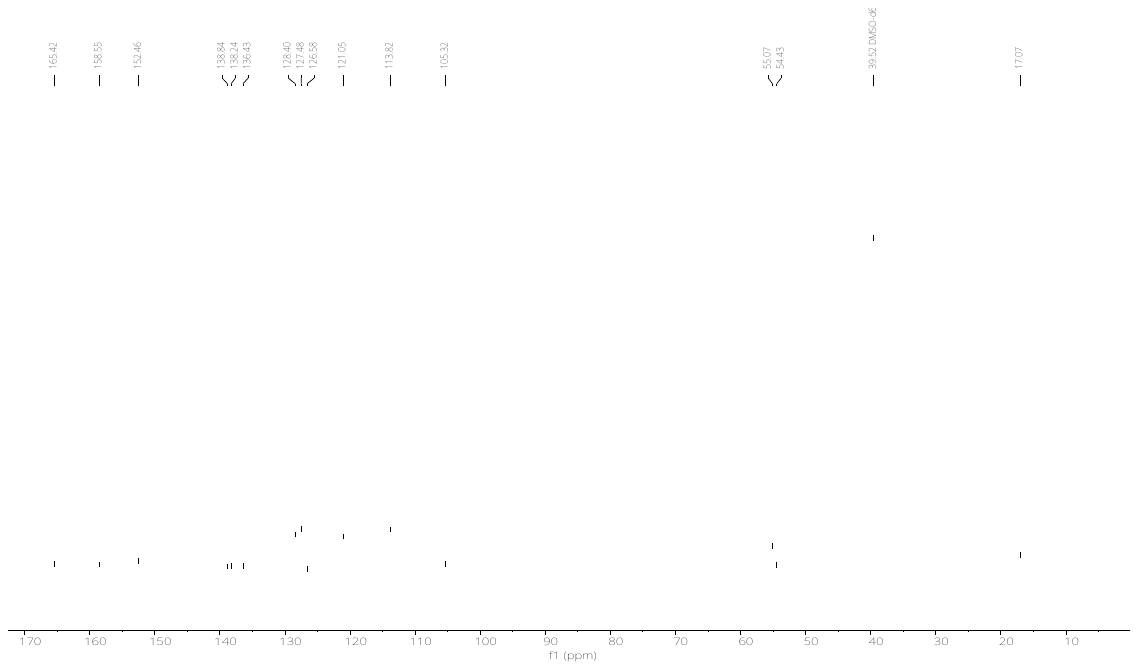

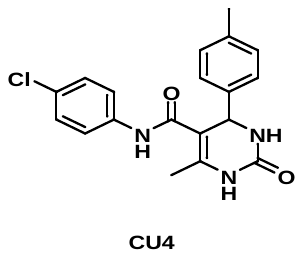

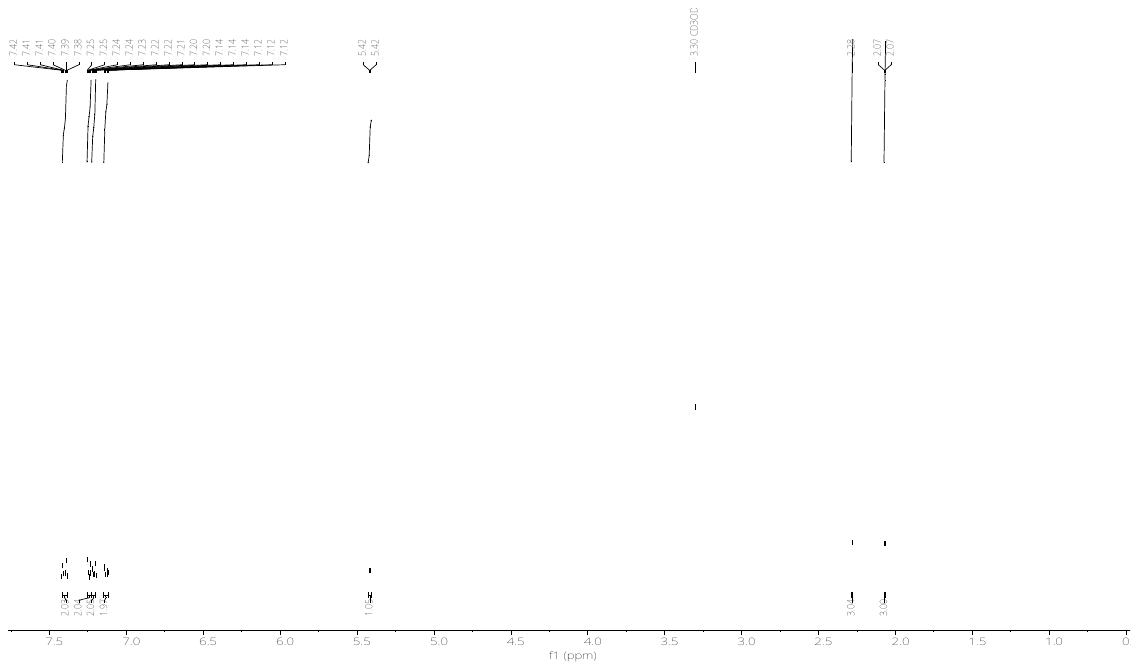

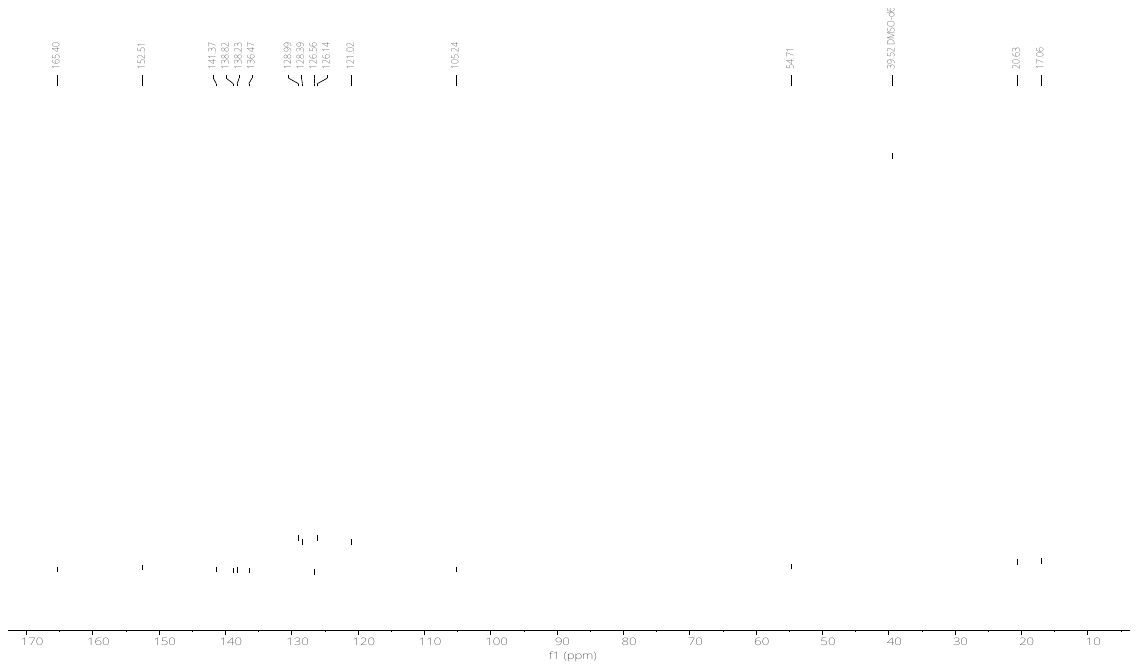

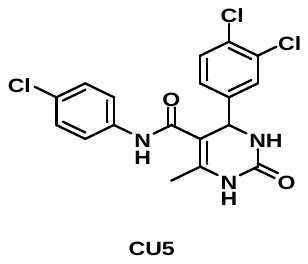

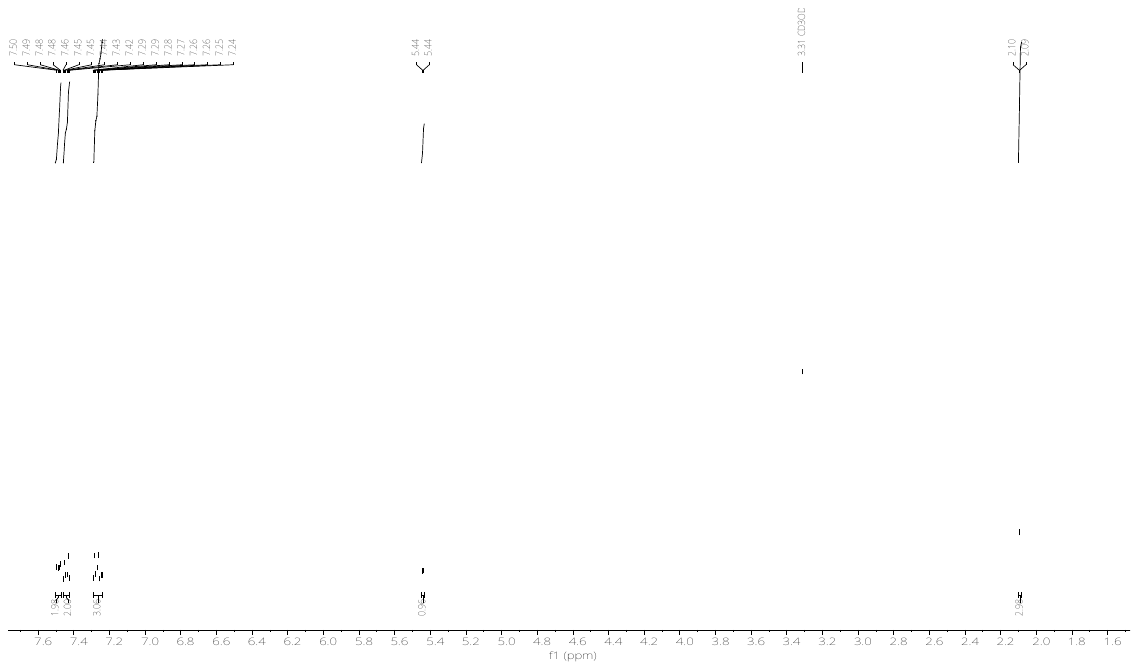

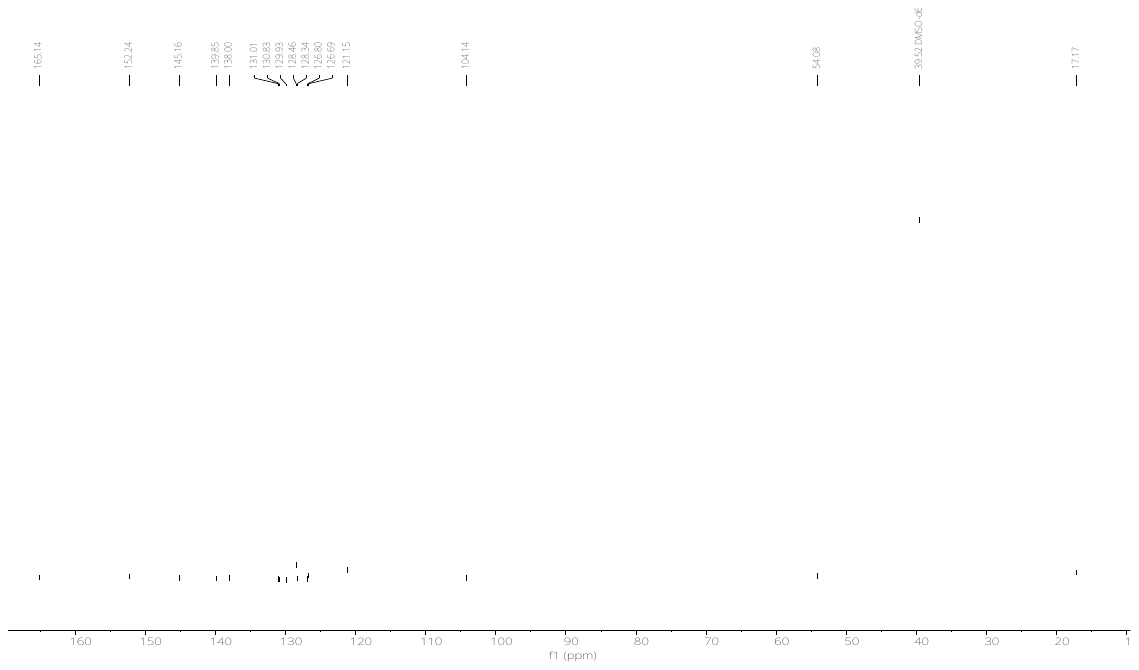

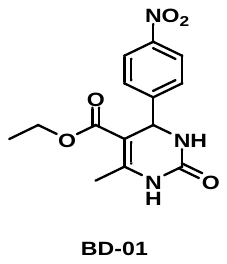

**U8**

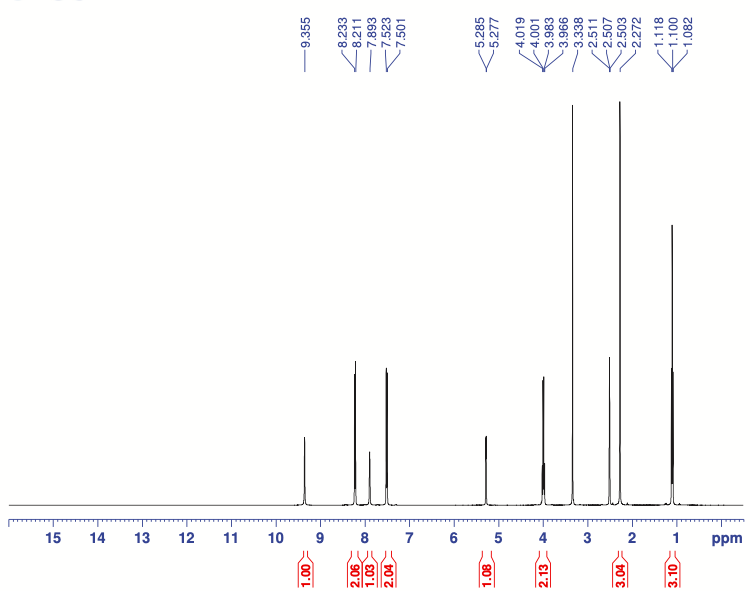

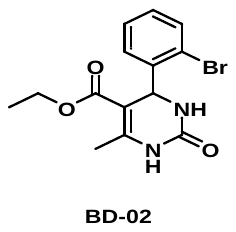

**U9**

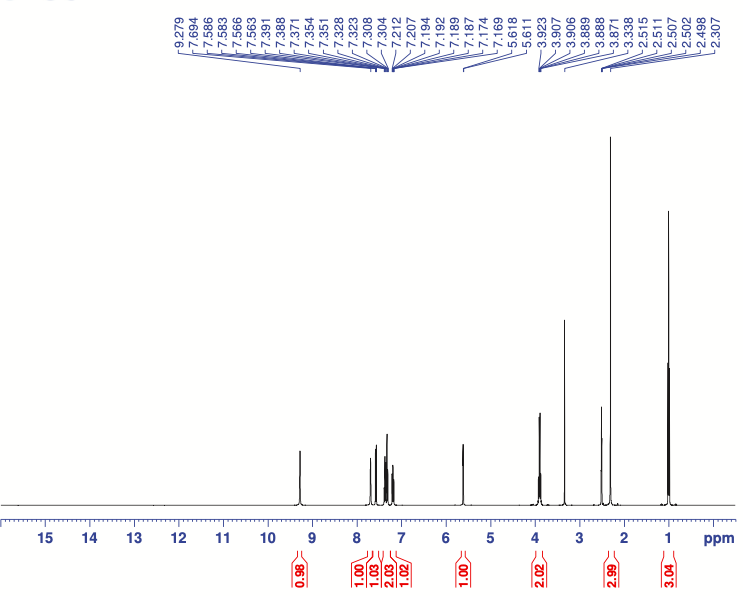

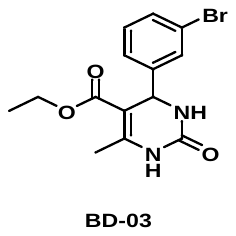

**U10**

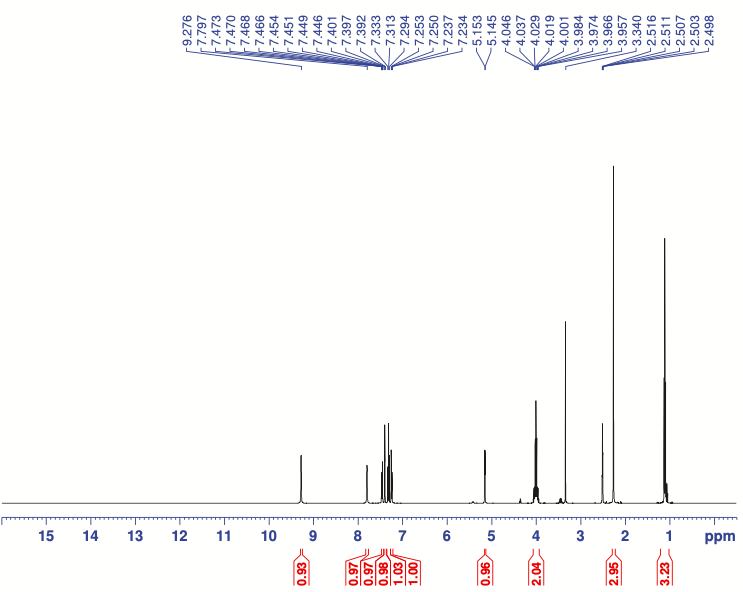

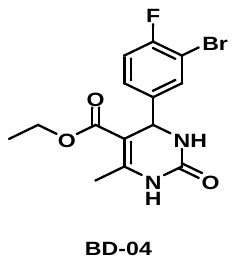

**U11**

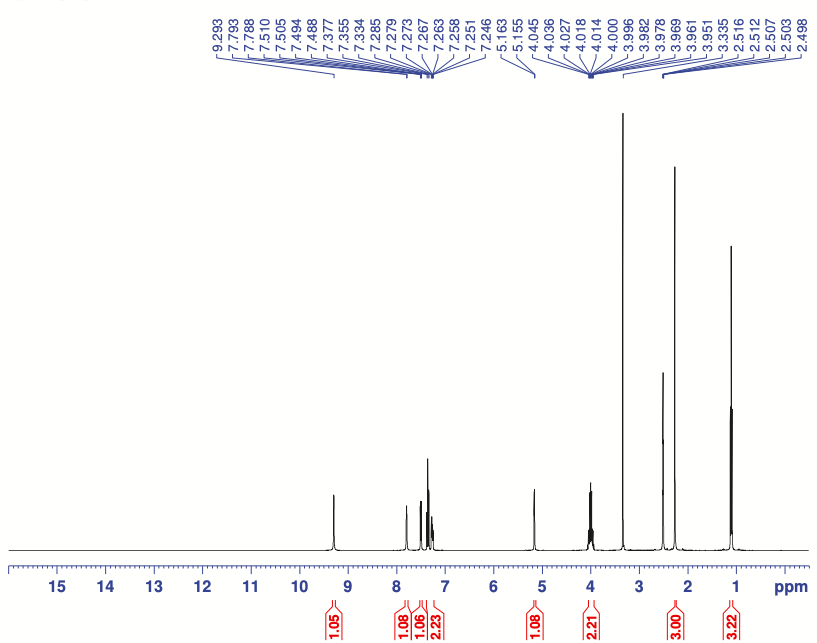

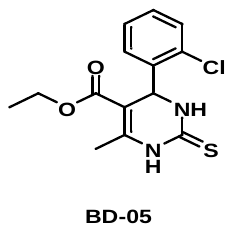

**T9**

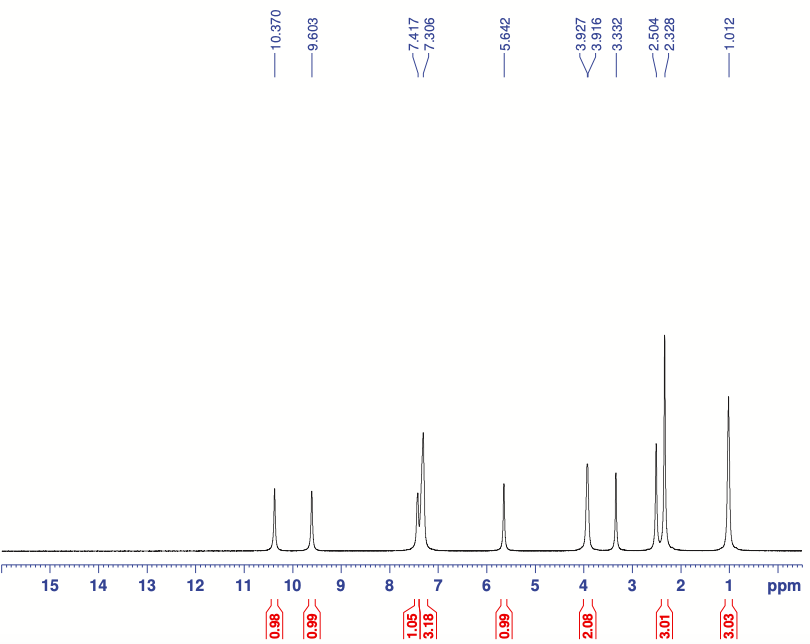

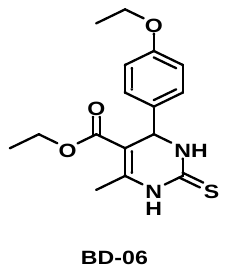

**T10**

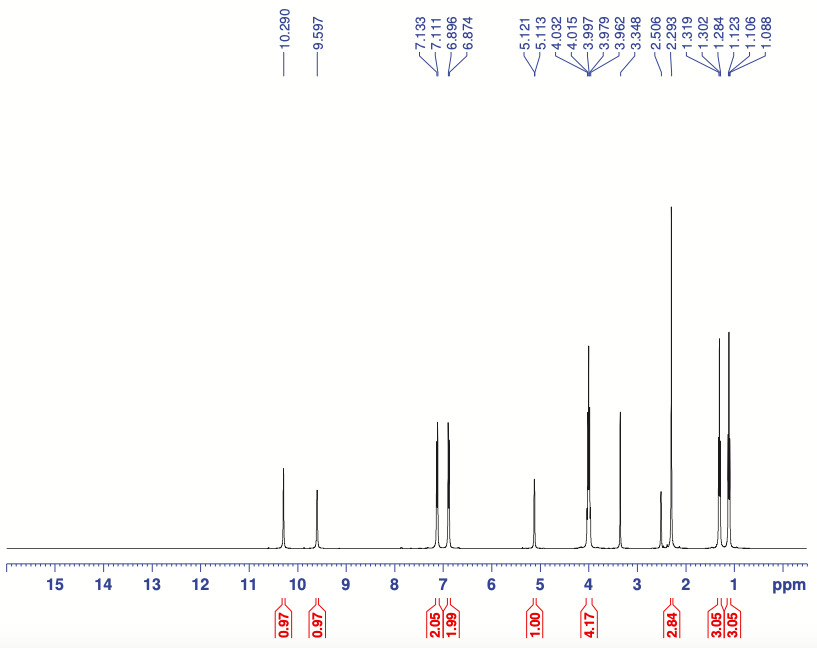

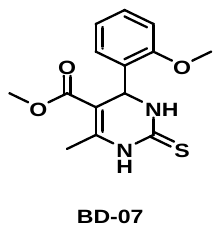

**T11**

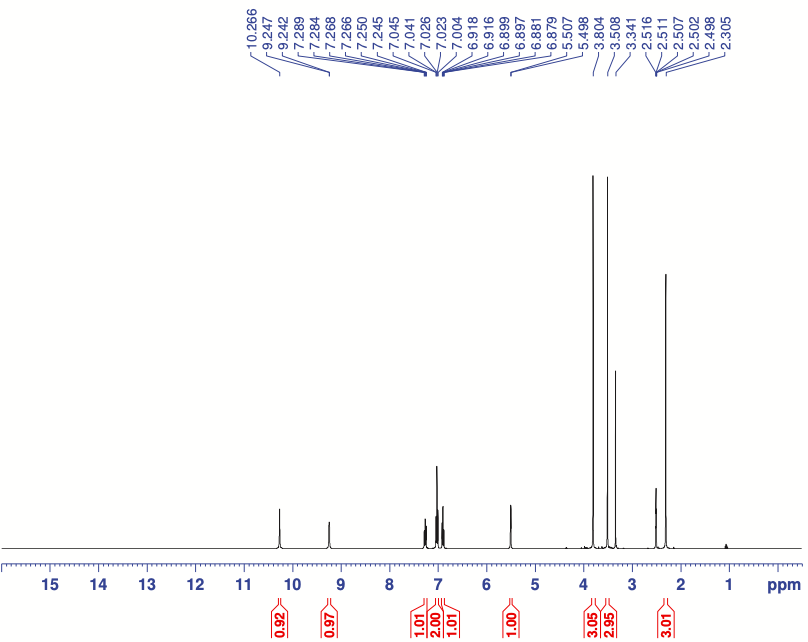

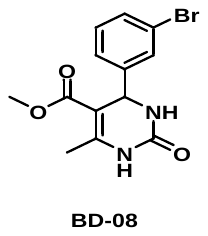

**U12**

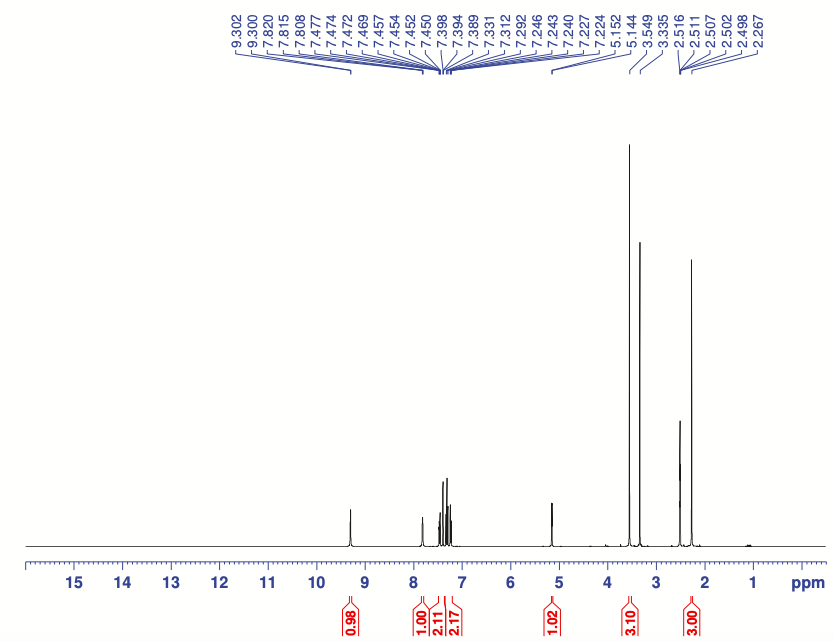

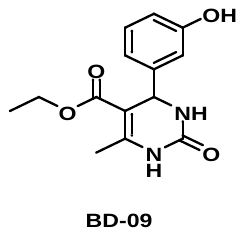

**U13**

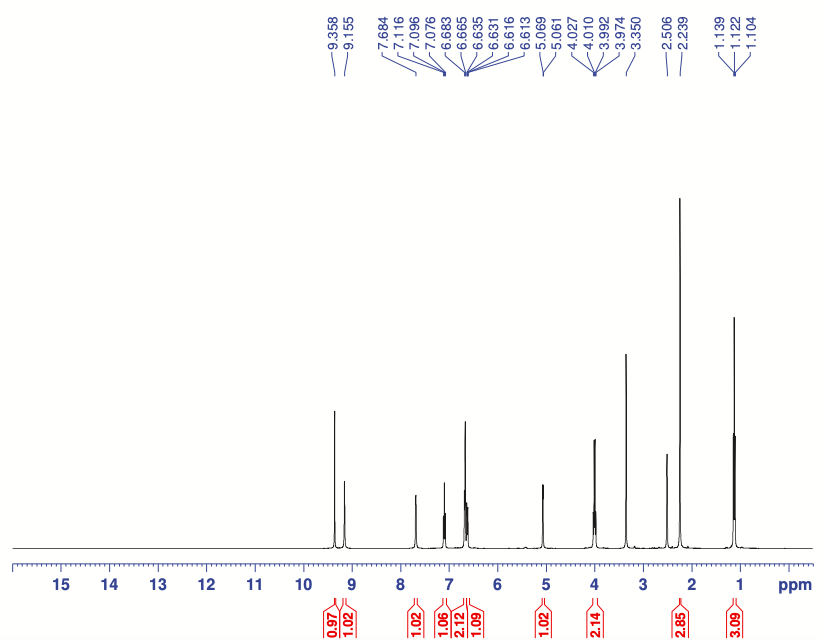

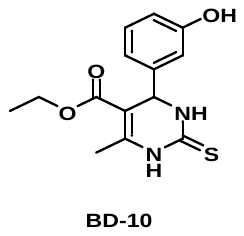

**T11**

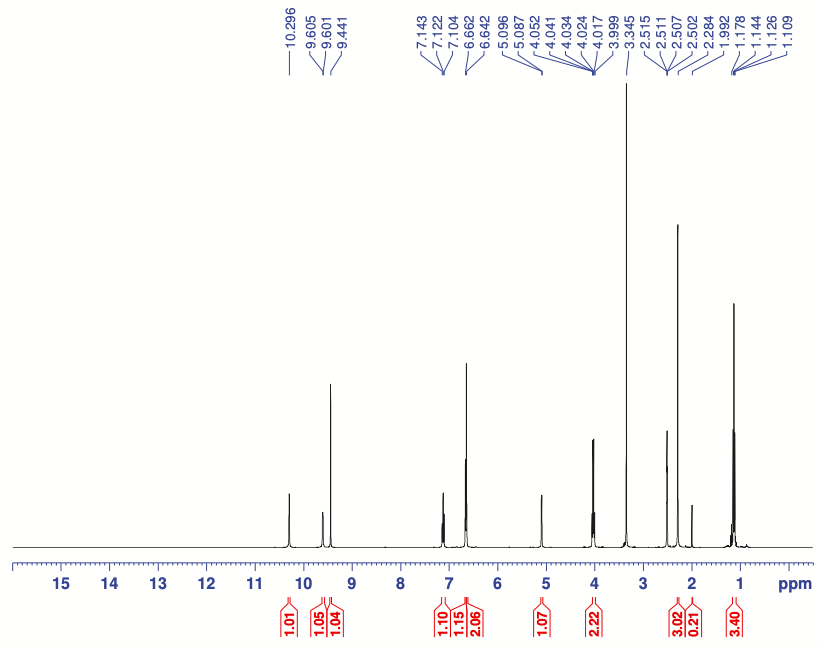

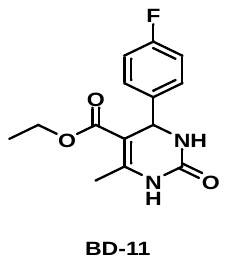

**U14**

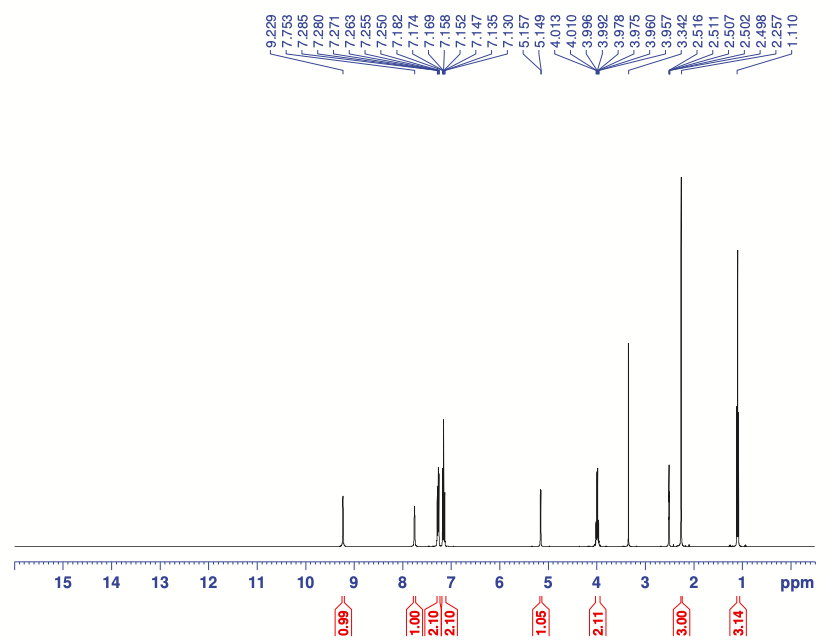

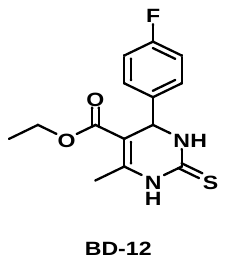

**T13**

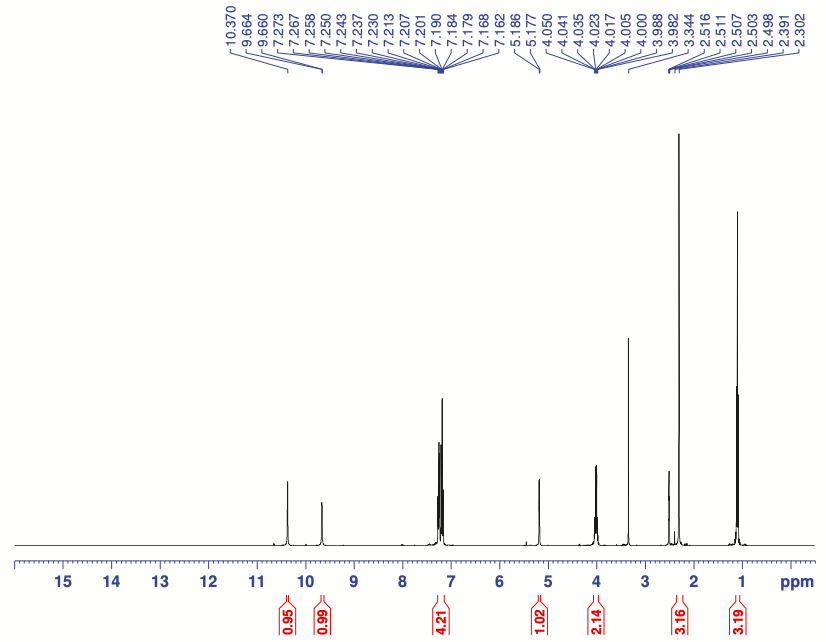

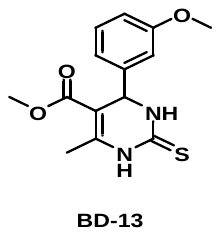

**T14**

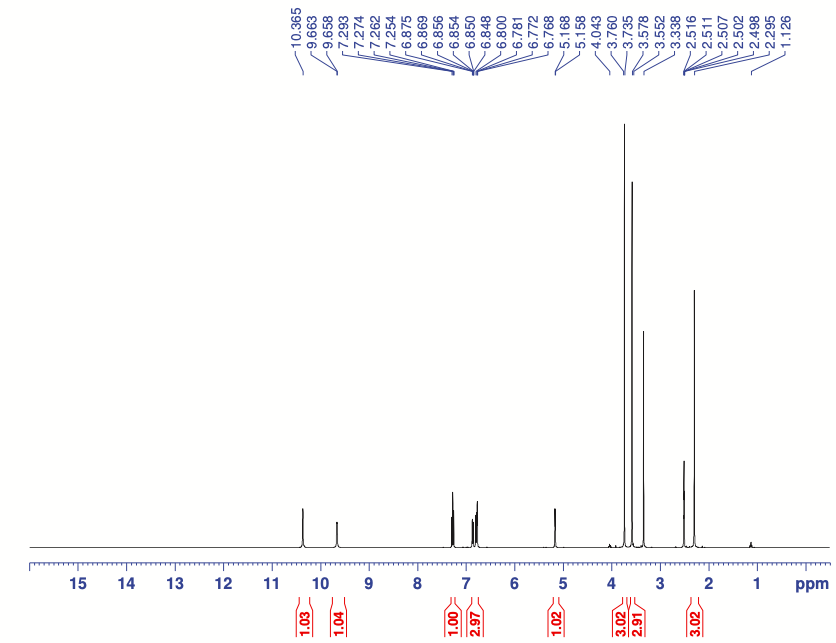

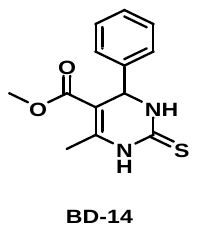

**U14**

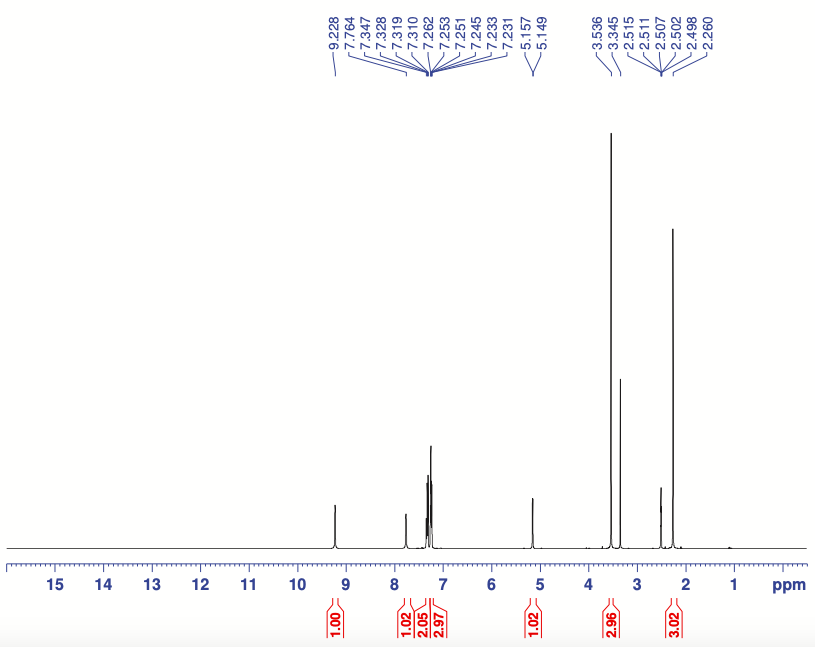
