## Supplementary 2 for "Dual Inhibitors Targeting G9a and GSK-3β: Translational Perspectives on Alzheimer’s Disease Treatment"

Compound preparation for *in vitro assays*...............................................................................................................S3

GSK-3β inhibition activity.......................................................................................................................................S3

G9a inhibition activity............................................................................................................................................S3

GLP inhibition activity............................................................................................................................................S3

Molecular dynamics..............................................................................................................................................S4

HMTs panel to evaluate inhibition activity...............................................................................................................S4

Kinase panel to evaluate inhibition activity..............................................................................................................S4

Blood-Brain Barrier Permeation Assay....................................................................................................................S4

Pharmacokinetics.................................................................................................................................................S5

Safety pharmacology profile..................................................................................................................................S6

In vitro determination of hERG activity....................................................................................................................S6

In vitro determination of microsomal stability at human microsomes......................................................................S6

In vitro determination of human plasma protein binding..........................................................................................S7

In vitro determination of human plasma protein binding..........................................................................................S7

Ames test………....................................................................................................................................................S7

Cell Micronuecleus test………................................................................................................................................S8

Acute oral toxicity……….........................................................................................................................................S9

*In vitro* efficacy studies in cells culture.................................................................................................................S10

*In vivo* efficacy studies in *Caenorhabditis elegans* (*C. elegans*)..............................................................................S13

*In vivo* efficacy studies in SAMP8..........................................................................................................................S14

ELISA determinations …………..............................................................................................................................S16

Protein Level Determination by Western Blotting (WB) ..........................................................................................S16

RNA extraction and gene expression determination...............................................................................................S17

MDA Levels.........................................................................................................................................................S17

ROS Levels .........................................................................................................................................................S17

Detection GSH/GSSG Levels................................................................................................................................S17

Determination Iron (II) Levels ...............................................................................................................................S17

Aβ plaques histology...........................................................................................................................................S18

Spine density and Golgi staining...........................................................................................................................S18

Transcriptomic analysis ......................................................................................................................................S18

Human cases …………….......................................................................................................................................S20

Statistical Analysis..............................................................................................................................................S21

References.........................................................................................................................................................S21

**Compound preparation for *in vitro* assays**

G9a/GSK-3β inhibitors were serially diluted between 5 nM and 0.001 nM in 100 % DMSO (Sigma, St. Louis, USA). Then, respective concentrations were diluted in MilliQ purified water to reach a final concentration ranging between 50 and 0.001 μM in 1 % DMSO in the well.

**GSK-3**β **inhibition activity**

GSK-3β activity was measured using the GSK3B Chemiluminescent Assay Kit (BPS Bioscience, #79700 San Diego, CA, USA), following the manufacturer’s instructions. Briefly, a master mix of 25 µL was prepared, consisting of 5 µL of 5x kinase assay buffer, 1 µL of ATP (500 µM), 5 µL of GSK substrate peptide (1 mg/mL), and 14 µL of nuclease-free water. Whole artery lysate (25 µL) was combined with 25 µL of this master mix in each well of a 96-well plate and incubated at 30°C for 45 minutes. After incubation, 50 µL of Kinase-Glo Max reagent (Promega) was added, and the mixture was incubated for 15 minutes at room temperature in the dark. Luminescence was then measured using a microplate reader (Biotek).

**G9a inhibition activity**

G9a enzyme activity was measured using the G9a Chemiluminescent Assay Kit (Catalog #52001L, BPS Bioscience, San Diego, CA, USA), following the manufacturer's protocol (REF). The assay included the histone H3 peptide substrate and the control compound UNC0642. The kit features wells precoated with histone H3 peptide substrate, an antibody specific to methylated lysine residues on histone H3, a secondary HRP-labeled antibody, S-adenosylmethionine (SAM), methyltransferase assay buffer, and purified G9a enzyme. In the assay, SAM was incubated with the sample containing assay buffer and methyltransferase enzyme for 1 hour. Following incubation, a primary antibody was added and incubated for 1 hour at room temperature. The wells were then treated with an HRP-labeled secondary antibody for 30 minutes, followed by the addition of an HRP substrate to generate chemiluminescence, which was measured using a chemiluminescence reader. The experiment was performed in triplicate. For IC50 determination, different concentrations of the compounds (10 µM, 1 µM, 0.1 µM, 0.01 µM, and 0.001 µM) were plotted on the X-axis, with percentage activity on the Y-axis. IC50 values were calculated by importing the dose-response data into GraphPad Prism.

**GLP inhibition activity**

Considering that G9a and GLP are members of the Suv39h subgroup of SET domain-containing molecules, together form a complex that methylates H3K9. However, the binding to GLP is undesirable because it gives rise to cytotoxicity. Then, to confirm the selectivity against GLP, we performed the *in vitro* GLP Chemiluminescent Assay Kit (Catalog # 53007, BPS Bioscience, San Diego, CA, USA). The kit is designed for convenience, featuring wells precoated with histone H3 peptide substrate and an anti-methylated lysine antibody, HRP-labeled secondary antibody, SAM, methyltransferase assay buffer, and purified GLP enzyme. In the assay, S-adenosylmethionine is incubated with a sample containing assay buffer and methyltransferase enzyme for 1h. After that, a primary antibody is added for 1h at RT. The wells are then treated with an HRP-labeled secondary antibody for 30 minutes, followed by the addition of an HRP substrate to generate chemiluminescence. The signal is measured using a chemiluminescence reader. The experiment was performed in triplicate.

**Molecular dynamics**

Molecular dynamic (MD) simulations help understand microscopic events like ligand binding and important macromolecular movements connected to it that take place in computationally simulated environment. Here we are performing MD simulation of GSK3β-T2 (5F95-T2) complex at ATP site as well as for G9a-T2 (5TTF-T2) complex at substrate site after adding missing Lys1092 and Asp1093 amino acid in crystal from AlphaFold 1 model of G9a (AF-Q96KQ7-F1-v4) using Protein Splice module of Maestro 2025-2 (Version 14.4) to understand the impact of missing amino acids in loop on stability of complex and future use of this complex for molecular modelling studies. Full atom MD simulations were performed for both protein ligand complex using Desmond 2025-2 (Maestro, version 14.4) in three steps: the system builder, minimization, and molecular dynamics simulation. A simple point charge (SPC) solvent model with an orthorhombic boundary box were used for building the model. The system was further neutralized by adding calculated amount of Na+ and 0.15 M of NaCl. The system was put through energy minimization for 100 ps to rule out any stearic conflicts. The simulation studies of 400 ns were performed post equilibration of a complex system finished with the constant-temperature constant-pressure (NPT) ensemble at 300 K temperature and pressure of 1 bar for both G9a-FLAV27 protein-ligand complex. The MD trajectory analysis, RMSD of protein, Protein RMSF, and Ligand RMSF were calculated after the MD simulations using md-out.cms file.^1^

**HMTs panel to evaluate inhibition activity**

We utilized an extensive Histone methyltransferases collection available at BPS Biosciences (<https://bpsbioscience.com>) in Luminescence assays. T2 compound was tested at a single dose.

**Kinase panel to evaluate inhibition activity**

We utilized an extensive kinase collection available at HD Biosciences (44 kinase panel, <https://www.hdbiosciences.com/EngKinase.htm>) in Luminescence assays. T2 compound was tested at single dose.

**Blood-Brain Barrier Permeation Assay**

To evaluate the brain penetration of the different compounds, a parallel artificial membrane permeation assay for blood-brain barrier was used, following the method described by Di, et al.^2^ The *in vitro* permeability (Pe) of fourteen commercial drugs through the lipid extract of porcine brain membrane together with the test compounds were determined. Commercial drugs and assayed compounds were tested using a mixture of PBS:EtOH (70:30). Assay validation was made by comparing the experimental permeability with the reported values of the commercial drugs by a bibliography and linear correlation between experimental and reported permeability of the fourteen commercial drugs using the parallel artificial membrane permeation assay was evaluated (Table S8). From this equation and taking into account the limits established by Di et al. for BBB permeation, we established the ranges of permeability as compounds of high BBB permeation (CNS+): *Pe* (10^-6^ cms^-1^) > 5.198; compounds of low BBB permeation(CNS-): *Pe* (10-6 cm s^-1^) < 2.054 and compounds of uncertain BBB permeation (CNS+/-): 5.198 > *Pe* (10-6 cm s^-1^) > 2.054. Results of the BBB-permeation for new compounds are in Table S1-S2.

**Pharmacokinetics**

Experimental procedure in mice

60 male mice (BW mean 40.2 ± 1.8 g) were randomly assigned (3 mice per time group) to a specific plasma and brain collection time and injected by i.p. with 10 mg/Kg of T2 base or by oral gavage (30 mice per route of administration and dose average administrated of 0.125 µg) dissolved in oil excipient. In anesthetised mice with isoflurane, blood and brain tissue were extracted by intracardiac punction and brain tissue removal at 2, 5, 10, 15, 30, 60, 120, 240 ,360 and 1440 min after administration.

T2 salt formulation dissolved in water has been oral administered by gavage (5 mg/kg) to 36 male mice (BW mean 38.7 ± 2 g). In anesthetized mice with isoflurane, blood, and brain tissue were extracted by intracardiac punction and brain removal at 5, 15, 30,120, 240, 360, 1440, 2160 and 2880 min after administration.

Chemical analysis and quantification of T2 concentrations in blood and plasma samples

To a 50 μL aliquot of brain homogenate was added 150 μL of cold acetonitrile containing 0.1% formic acid and the internal standard AC009. After vortex mixing for one minute, the samples were centrifuged for 15 minutes at 13300 rpm. The temperature of the centrifuge was set to 4 °C. An aliquot of 150.0 μL of the supernatant was transferred to a vial for HPLCMS/MS analysis.

Aliquots of 50 μL of plasma samples were mixed with 150 μL of ACN with 0.1% formic acid containing AC009 as internal standard, centrifuged at 13,300 rpm for 15 min and supernatant removed. Aliquot supernatant samples were kept at -20 ºC till HPLC-MS/MS analysis. The internal standard AC009 was obtained from the supplier and of T2.

HPLC-MS/MS analysis was performed using an Agilent 1290 HPLC System coupled to a mass spectrometer Api4000 SCIEX. Gradient chromatographic conditions were applied (A: water 0.1% formic acid /ACN (90/10), B: ACN/water 0.1% formic acid (90/10), with 0.4 mL flow rate, oven temperature 20ºC and autosampler temperature set at 4ºC. A volume of 5 μL of sample was injected to an analytical HPLC column C18 2.1 x5 mm, 3µm particle size (Supelco). MSMS detection of analyte and internal standard were carried out in electrospray positive ionization (m/z analyte: 381.316/91.0; 381.316/204.0 and m/z internal standard: 379.407/91.0).

Analytical method validation was carried out for the detection and quantitation of T2 in brain and plasma samples meet acceptance criteria for linearity (range 5-1280 ng/mL), precision and accuracy (CV ± 20% of theoretical concentration). The upper and lower limits of T2 quantitation in plasma and brain samples were stablished at 5 ng/mL and 1280 ng/mL respectively.

**Safety pharmacology profile**

Safety 44 panel testing was performed at <https://wuxibiology.com/resource/mini-safety-panel/>. T2 compound was tested at a single dose.

***In vitro* determination of hERG activity (safety test)**

Determining hERG activity was performed because blocking hERG channels is a major therapeutic challenge in drug discovery due to its importance in ensuring safety (Kalyaanamoorthy & Barakat, 2018).

CHO cells that stably express hERG channels (Millipore) were cultured in F12 HAM medium supplemented with 10% FBS and 400 μg/L Geneticin. The extracellular Ringer’s solution consisted of the following (in mM): 2 CaCl2, 1 MgCl2, 10 HEPES, 4 KCl, 145 NaCl, and 10 glucose, with a pH of 7.4 and 305 mOsm. The intracellular Ringer's solution was composed of the following (in mM): 5.37 CaCl2, 1.75 MgCl2, 31.25/10 KOH/EGTA, 10 HEPES, and 210 KCl, with a pH of 7.2 and 295 mOsm. Shortly before use, 4 mM Na2-ATP was added to the intracellular Ringer's solution. Whole-cell currents were measured using a QPatch system (Sophion) in response to voltage protocols executed continuously, as recommended by the manufacturer. At the onset of the voltage protocol, cells were held at a Vh of -80 mV. They were then briefly clamped to -50 mV (20 ms), depolarized to 20 mV for 4800 ms, and finally repolarized to -50 mV for 5000 ms, during which the peak outward tail current was measured. The voltage was then returned to Vh for 3100 ms. This voltage protocol was repeated every 15 seconds. For each cell, the extracellular solution was applied before increasing the concentrations of the tested compound. Then, we select 4 compounds to evaluate their hERG activity. Higher IC_50_ values indicate lower affinity, hence suggesting greater safety.

***In vitro* determination of microsomal stability at human microsomes**

The human microsomes employed were purchased from Tebu−Xenotech. The compound was incubated at 37 °C with the microsomes in a 50 mM phosphate buffer (pH = 7.4) containing 30 mM MgCl_2_, 10 mM NADP, 100 mM glucose-6-phosphate, and 40 U/mL glucose-6-phosphate-dehydro- genase. Samples (75 μL) were taken from each well at 0, 10, 20, 40, and 60 min and transferred to a plate containing 4 °C 75 μL of acetonitrile and 30 μL of 0.5% formic acid in water were added to improve the chromatographic conditions. The plate was centrifuged (46,000g, 30 min). Supernatants were taken and analyzed by an ultraperformance liquid chromatograph−tandem mass spectrometer (Xevo-TQD, Waters) by employing a BEH C18 column and an isocratic gradient of 0.1% formic acid in water: 0.1% formic acid acetonitrile (60:40). The metabolic stability of the compounds was calculated from the logarithm of the remaining compounds at each of the time points studied.

***In vitro* determination of human plasma protein binding (bioavailability test)**

In drug discovery, it is essential to note that low binding to plasma proteins is desirable as it allows more drugs to be available in plasma to exert their effect (Bohnert & Gan, 2013). However, if the binding is too low, the drug will be eliminated quickly, so a balance is sought. To assess bioavailability, we collaborated with Seralab.

The assay used Rapid Equilibrium Dialysis (RED) from Thermo Scientific. Compounds were dissolved at a concentration of 5 μM in plasma and added to the corresponding insert of the RED device. Dialysis buffer was added to the corresponding insert of the RED device. The plate was then incubated at 37ºC for 4 hours. After incubation, 50 μL aliquots were taken from each chamber and transferred to empty vials. For the plasma samples, 50 μL of dialysis buffer was added, while for the buffer samples, 50 μL of plasma was added. All samples were treated with 300 μL of acetonitrile and centrifuged at 4000 rpm. Next, 100 μL aliquots of the supernatants were transferred to an LC analysis plate and analyzed using a UPLC/MS/MS. The stationary phase consisted of ACQUITY BEH C18 1.7μm 2.1x50mm (Waters)/ ACQUITY HSST3 1.8μm 2.1x100mm (Waters). The gradient was A (H2O + 0.1% Formic acid) and B (ACN + 0.1% Formic acid), with a flow rate of 0.6mL/min. The ACQUITY UPLC/Xevo TQD System was used for chromatography. Compound concentrations were determined based on the MS peak areas.

***In vitro* determination of human plasma protein binding**

The assay used Rapid Equilibrium Dialysis (RED) from Thermo Scientific. Compounds were dissolved at a concentration of 5 μM in plasma and added to the corresponding insert of the RED device. Dialysis buffer was added to the corresponding insert of the RED device. The plate was then incubated at 37ºC for 4 hours. After incubation, 50 μL aliquots were taken from each chamber and transferred to empty vials. For the plasma samples, 50 μL of dialysis buffer was added, while for the buffer samples, 50 μL of plasma was added. All samples were treated with 300 μL of acetonitrile and centrifuged at 4000 rpm. Next, 100 μL aliquots of the supernatants were transferred to an LC analysis plate and analyzed using a UPLC/MS/MS. The stationary phase consisted of ACQUITY BEH C18 1.7μm 2.1x50mm (Waters)/ ACQUITY HSST3 1.8μm 2.1x100mm (Waters). The gradient was A (H2O + 0.1% Formic acid) and B (ACN + 0.1% Formic acid), with a flow rate of 0.6mL/min. The ACQUITY UPLC/Xevo TQD System was used for chromatography. Compound concentrations were determined based on the MS peak areas.

**Ames test**

The mutagenicity of T2 was examined in a bacterial reverse mutation assay conducted with and without S9 mix, using amino-acid requiring *S. typhimurium* strains TA100 and TA98. The results were compared with three different positive controls which are mutagenic and suspected to be carcinogenic agents: 2-nitrofluorene (2-NF) and sodium azide (NaN3) for TA98 without S9 mix, 4-nitroquinoline-N-oxide (4-NQO) for TA100 without S9 mix, and 2- aminoanthracene (2-AA) for both TA98 and TA100 with S9 mix. TA98 strain is meant to detect frameshift mutations while TA100 is meant to detect base-pair substitutions. Also, both TA98 and TA100 S. typhimurium strains have rfa mutations, which result in a defective lipopolysaccharide layer that makes their cell wall more permeable to larger molecules, uvrB mutations, which eliminate excision repair of DNA damage, and the pKM101 plasmid which increases error-prone repair of DNA damage.

A commercial test kit, Ames MPFTM 98/100 Microplate Format Mutagenicity Assay, from Xenometrix AG (Allschwill, Switzerland) was used to evaluate the mutagenicity of T2. This kit is based on the most common bacterial reverse-mutation test, known as the “Ames Test”,^5^ and on an assay performed entirely in liquid culture, which is the “Fluctuation test”, originally devised by Luria and Delbruck (1943) and modified by Hubbard et al.^6^ This assay provides a few benefits over traditional agar plates, and it is cited in the guidelines of the OECD and the FDA^7,8^.

Bacteria (*S. typhimurium* strains TA100 and TA98) were exposed to 4 concentrations of T2 along with a positive and a negative control for 90 minutes in medium containing enough histidine to support approximately two cell divisions. After incubation, the cultures were diluted in pH indicator medium lacking histidine, and aliquoted into 48 wells of a 384-well plate. Within two days, cells that had undergone reversion to histidine prototrophy grew into colonies. Bacterial metabolism reduces the pH of the medium, changing the color from purple to yellow. Accordingly, all yellow, partially yellow or turbid wells were scored as positive, while all purple wells were scored as negative. The number of wells containing revertant colonies are counted for each dose and compared to a negative control (blank with solvent). A dose-dependent increase in the number of revertant colonies upon exposure to the chemical compared to the controls indicates that the sample is mutagenic in the Ames MPF™ 98/100 assay. The mutagenic potential of substances is assessed directly and in the presence of metabolic activation.

Test bacterial strains

Two mutant strains, *S. typhimurium* TA98 and *S. typhimurium* TA100 were used to detect frameshift and base-pair mutation, respectively. The bacteria were kept frozen and stored in total darkness until used. Both bacteria were inoculated in nutrient broth and incubated at 37ºC, 250 rpm for 14-16 hours prior to the test.

**Cell Micronucleus Test**

The protocol follows the recommendations of Test Guideline 487 (TG-487) of OECD guideline for the testing of chemicals^9^. The test was performed on rodent CHO cells (ECACC Ref.: 85050302). The cells are seeded at a density of 2000/well in a black 96-well plate with clear bottom and are incubated in a humidified atmosphere at 37ºC with 5% CO_2_. To estimate the micronuclei frequency, the cells scored must have completed one mitosis during the treatment or the post-treatment incubation period. Compound assayed was T2, stock wase prepared at 20 mM in DMSO 100 and was assayed at 200, 100, 50, 25 and 12.5 μM, during 24 h in six replicates. DMSO should not exceed 1% according to TG-487. Mitomycin C (MitC, Sigma Aldrich), a known inducer of micronuclei formation, was the positive control used to demonstrate the sensitivity of the test, and cells untreated are negative control. After treatment, Cytochalasin B (cytoB) is used as cytokinesis-blocker of cultures during 28 h. Cells are then fixed with 3.7% formaldehyde and 1% Triton X-100 and nuclei are stained with bisbenzimide (Hoechst dye no. 33258) for 30 min at room temperature. Imaging acquisition is performed by using Operetta CLS High-Content Analysis System (Perkin Elmer). Analysis is performed using Harmony software of Perkin Elmer and the in-house App NucleusFinder^10^, based on an open source processing image program, ImageJ.

*NucleusFinder* identifies regularly shaped mononuclear, binuclear and multinuclear cells, excludes irregular, small and isolated nuclei (odd nuclei) and detects valid micronuclei following very conservative conditions as cytoplasmic location without connection with the main nuclei and proper size. NucleusFinder chooses the best analysis algorithm for each image capture from six implemented filters, using a smart fit choice calculation. It allows to perform a classifying and a counting of the different elements. The cytokinesis-block proliferation index (CBPI), which indicates the average number of cell cycles per cell during the period of exposure to cytoB, is used to estimate the cytostatic activity of a treatment by comparing values in the treated and control cultures. Cytostasis percentage should not to exceed 60% because higher levels may induce micronuclei as a secondary effect of cytotoxicity and is calculated as follows:

% Cytostasis = 100 - 100 {(CBPIT - 1) ÷ (CBPIC -1)}

Where:

T = test chemical treatment culture

C = vehicle control culture

And:

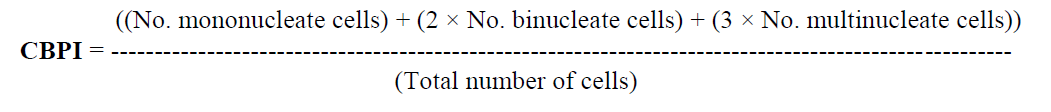

**Acute oral toxicity**

Compound T2 was dissolved in PEG-400 (Merck) at 100, 320, 1000, and 2000mg/kg doses.

The acute oral toxicity test, known as the Up-and-Down Procedure (UDP), follows the OECD Test Guideline 425 with minor modifications. This test involves administering a single dose to mice at intervals of at least 48 hours. The initial dose is set below the estimated lethal dose (LD50). If the first animal survives, the subsequent dose is increased by a factor of 3.2; if it dies, the next dose is decreased similarly. Mice were monitored for two weeks after dosing. The maximal tolerated dose (MTD) is defined as the highest dose that does not result in death or significant adverse effects in the animals.

The experimental groups were composed of three female mice, according to Test Guideline OECD 425. The initial dosing for the first three mice per compound was set at 100 mg/kg, with each mouse receiving 200 μl of an 18 mg/ml preparation. After a 48-hour observation period, no toxicity was noted in these animals. Subsequently, a second group was administered 320 mg/kg, using 200 μl of a 56 mg/ml preparation per mouse. Again, after 48 hours, no signs of toxicity were observed in either the first or second groups. Following this, a third group received a dose of 1 g/kg, with each mouse receiving 200 μl of a 174 mg/ml preparation. After another 48-hour period, toxicity was still absent. The fourth group was treated with T2 at the maximum recommended dose of 2g/kg, using 200 μl of a preparation that was intended to be 350 mg/ml. However, this concentration proved insoluble.

Mice from the groups were subjected to a comprehensive observation period lasting 14 days, with assessments occurring every 48 hours. Observations focused on alterations in skin and fur, as well as the condition of the eyes and mucous membranes. Evaluations covered the respiratory, circulatory, autonomic, and central nervous systems, alongside somatomotor activity and behavioral patterns. Specific attention was given to signs of tremors, convulsions, salivation, diarrhea, lethargy, sleep disturbances, and coma. During this period, any mice found in a severely ill state or exhibiting significant pain or distress were euthanized humanely. Euthanasia was implemented to alleviate unrelieved pain and distress that could not be controlled by other means.

***In vitro* efficacy studies in cells culture**

Primary Glial Cultures

Primary glial cells were isolated from the cerebral cortex of mice as previously described ^11^. Briefly, the brain was dissected, and the cerebral cortex was isolated, dissociated, and incubated with 0.25% trypsin/EDTA at 37 °C for 1 hour. After centrifugation, the tissue underwent several washes with HBSS (Gibco), and the cells were plated in uncoated flasks, maintained in a 1:1 HAMS/DMEM medium containing 10 % FBS + 10 % horse serum + 1 % penicillin/streptomycin. After 7 days in culture under standard conditions, the flasks were agitated on an orbital shaker at 240 rpm for 4 hours at 37 °C. The supernatant was collected, centrifuged, and the resulting pellet, containing microglial cells, was resuspended in complete medium (HAMS/DMEM (1:1) with 10% FBS) and seeded into uncoated 96-well plates. After 2 hours of adhesion, the medium was changed to remove non-adherent oligodendrocytes, and fresh medium containing 10 ng/mL GM-CSF was added. Astroglial cells remaining adherent in the flasks were trypsinized, collected, centrifuged, and plated into 96-well plates with complete medium. Culture purity, confirmed by immunofluorescence using Iba-1 (microglial marker) and GFAP (astrocyte marker) antibodies, was greater than 98%. After 2 days in culture, cells were pretreated for 1 hour with the compounds (1 μM), followed by treatment with bacterial lipopolysaccharide (LPS; 10 μg/mL) for 24 hours. Nitrite production in the cultures was then assessed.

Cell Viability Assay

Cell viability was assessed using the MTT assay (Roche Diagnostic, GmbH, Basel, Switzerland), which relies on the ability of living cells to convert yellow MTT into blue formazan. In brief, cells were cultured in 96-well plates and exposed to the specified compounds for 18 hours. After treatment, cells were incubated with MTT (0.5 mg/mL) for 1 hour in the dark, followed by solubilization with 10% dimethyl sulfoxide (DMSO). The reduction of MTT was quantified by measuring absorbance at 595 nm, following the manufacturer's instructions. Data presented are the average of at least three independent experiments.

Nitrite Measurement

Nitrite accumulation in the culture medium was determined using the standard Griess reaction. Following 24 hours of cell stimulation with LPS (previously, cultures were treated with the different compounds as above mentioned), supernatants were collected and combined with an equal volume of Griess reagent (Sigma-Aldrich). The mixtures were incubated at room temperature for 15 minutes, and absorbance was measured at 492/540 nm using a plate reader.

Neuronal primary cultures and mixed neuronal and microglial primary cultures

To prepare primary cultures of striatal neurons and mixed neuronal and microglial primary cultures, brains from fetuses of pregnant C57BL/6J mice or from one day mice, respectively, were removed (gestational age: 19 days). Cells were isolated as described in Hradsky et al.^12^. Briefly, the samples were dissected and, after a careful removal of the meninges, digested for 20 min at 37ºC with 0.25% trypsin. Trypsinization was stopped by adding an equal volume of culture medium (supplemented DMEM). Cells were brought to a single-cell suspension by repeated pipetting followed by passage through a 100 μm-pore mesh. Pelleted (5 min, 200Å~g) cells were resuspended in supplemented DMEM and seeded at a density of 3.5Å~105cells/mL in 6-well plates. The day after, to prepare neuronal primary cultures, the medium was replaced by neurobasal medium supplemented with 2 mM L-glutamine, 100 U/mL penicillin/streptomycin and 2 % (v/v) B27 medium (GIBCO) while to prepare mixed neuronal and microglial primary cultures the medium was not exchanged. All cultures were assayed 12 days after.

Neurite patterning determination

Cortical neuronal primary cultures were treated with Aβ1-42 (500 nM) for 48 h on DIV 10. Next day, neurons were treated with T2 (1 µM) or vehicle without exchanging the medium to mateine the growth factors released. Then, cells were fixed in 4% paraformaldehyde for 15 min and then washed twice with PBS containing 20 mM glycine followed by permeabilization with the same buffer containing 0.2% Triton X-100 (15 min incubation). Samples were treated for 1 h with blocking solution (PBS containing 1% bovine serum albumin) and labeled with polyclonal rabbit anti-Nectin 3 antibody (Abcam, 1/1000) to detect neurite pattering. Neurons were detected with anti-F-actin antibody fused to an Alexa 488 fluorophore (ThermoFisher, 1/400) to detect the cytoskeleton and thus the cell morphology. Then, sections were incubated at RT for 2 h with a Cy3-conjugated anti-rabbit secondary antibody (1/200, 711-166-152, Jackson ImmunoResearch). Samples were washed several times with PBS and mounted with 30% Mowiol (Calbiochem, San Diego, CA, USA). Nuclei were stained with Hoechst (1/100). Samples were observed under a Zeiss 880 confocal microscope (Leica Microsystems, Wetzlar, Germany). Quantification of neurite formation was performed over segments of 15 μm. Each red dot represents a neurite formation.

Protein aggregation detection

Cortical neuronal primary cultures were treated with Aβ1-42 (500 nM), or vehicle for 48 hours and subsequently stimulated with 200 nM CBDor vehicle. Then, cells were fixed in 4% paraformaldehyde for 15 min and then washed twice with PBS containing 20 mM glycine before permeabilization with the same buffer containing 0.2% Triton X-100 (15 min incubation). The samples were treated for 1 h with blocking solution (PBS containing 1% bovine serum albumin) and labeled with a rabbit anti-Aβ antibody (1/737 100, ab201060) and subsequently marked with a Cy3 anti-rabbit (1/200, Jackson InmunoResearch) secondary antibody (red). Following 2 h of incubation, cells were washed and subsequently imaged using confocal microscope with 25X (yellow squares) and 40X (green squares) objectives (Zeiss LSM 880).

ELISA analysis of cytokine secretion

Mature iPSC-derived microglia (50 000/cm^2^) were treated with T2 (1 μM) or (UNC0642, 1 μM) for 24h and cytokine release was measured in the culture media at basal conditions or upon 24h stimulation with LPS/IFN-γ (100 µg/ml and 10 U/ml respectively). Briefly, following the 24h pre-treatements, cell culture medium was collected, centrifuged at 300*×*g for 15 min to remove cell debris and typically 50 μl (1:2 dilution) appraised for secreted TNFα or 200 μl (1:2 dilution) for secreted IL-1β using human Quantikine ELISA kits, as per the manufacturer’s instructions (R&D).

Electrophysiology

Hippocampal neurons cultured in vitro were used for recordings between 18 and 20 DIV, following a 72-hour treatment with either DMSO (as vehicle control) or 1 μM T2. Coverslips containing the neurons were mounted onto the stage of an inverted microscope (Olympus IX50 or Axio-Vert.A1 Zeiss) for electrophysiological recording using whole-cell patch-clamp configuration. Recordings were performed at room temperature (22–24 °C) using an Axopatch 200B amplifier connected to a Digidata 1440A interface and controlled via pClamp10 software (Molecular Devices).

The external recording solution consisted of (in mM): 140 NaCl, 3.5 KCl, 10 HEPES, 20 glucose, 1.8 CaCl₂, and 0.8 MgCl₂, with pH adjusted to 7.42 using NaOH. To pharmacologically isolate AMPA receptor-mediated miniature excitatory postsynaptic currents (mEPSCs), the solution was supplemented with 1 μM tetrodotoxin (TTX), 50 μM D-AP5, and 100 μM picrotoxin to block action potentials, NMDARs and GABA-A receptor-mediated currents respectively. All pharmacological agents were obtained from Abcam.

Patch electrodes were fabricated from borosilicate glass capillaries (1.2 mm outer diameter, 0.69 mm inner diameter; GC120F-10, Harvard Apparatus) using a P-97 horizontal puller (Sutter Instrument Co.), yielding a final tip resistance of 3.5 to 7 MΩ. Electrodes were filled with an internal pipette solution containing (in mM): 116 K-gluconate, 6 KCl, 8 NaCl, 10 HEPES, 0.2 EGTA, 2 MgATP, and 0.3 Na₂GTP, adjusted to pH 7.2 with KOH.

During recordings, neurons were voltage-clamped at –70 mV. Series resistance (Rs), typically ranging from 15 to 25 MΩ, was measured at the start and end of each recording. Cells exhibiting Rs fluctuations exceeding 15% were excluded from analysis. Miniature EPSCs were low-pass filtered at 2 kHz, digitized at 5 kHz, and analyzed using IGOR Pro (WaveMetrics) in combination with the Neuromatic 2.03 package.^13^

Events were detected using amplitude threshold of typically ~5–6 pA, defined based on baseline noise. Only synaptic events with a rapid, monotonic rise time (<1.5 ms) and clean decay phases were included for amplitude analysis. Statistical comparisons were conducted using GraphPad Prism version 8.0.1 for macOS (GraphPad Software, San Diego, CA, USA). Group comparisons were made using two-tailed Student’s t-tests, with significance thresholds set at **p < 0.01 and ***p < 0.001.

***In vivo* efficacy studies in *Caenorhabditis elegans* (*C. elegans*)**

C*. elegans* strains and maintenance

The WT *Caenorhabditis elegans* (*C. elegans*) strain (N2), and the transgenic CL2006 strain (dvIs2 [pCL12(unc-54/human Aβ peptide 1–42 minigene)+rol-6(su1006)]) provided by the *C. elegans* Genetic Center were used. N2 worms were propagated at 20°C, while CL2006 worms were maintained at 16°C in a temperature-controlled incubator on a solid nematode growth medium (NGM) seeded with *Escherichia coli (E. coli)* OP_50_ (Carolina Biological) strain as a food source^37^.

Treatment of *C. elegans* and compound preparation

All pharmacological experiments employed inactivated *E. coli* OP50 strain as a dietary supplement. In Luria broth (LB) media, *E. coli* OP50 bacteria were grown for 16h at 37ºC and 200 rpm. After growth, a bacterial suspension was transferred to 50 mL tubes and centrifuged for 30 min at 400 g and 4ºC. Supernatants were discarded, and pellets were subjected to three freeze/thaw cycles, switching between liquid nitrogen and a 37°C water bath to inactivate them. Lastly, liquid nitrogen was used to freeze the *E. coli* OP50 pellets, and they were kept at -80°C. All drug experiments were done in 96-well plate format (liquid culture). Adult animals that were five days old were handled with an alkaline hypochlorite solution for 5 minutes with heavy shaking before being centrifuged for 1 minute at 20°C and 2000 rpm. The egg pellets were additionally rinsed three times in M9 buffer. To get the worms to the L1 stage, egg pellets were resuspended in S-medium (30 µl 1M MgSO_4_, 30 µl CaCl_2_, 100 µl 100x trace metal solution, 100 µl 1M Potassium citrate (pH = 6.0), and 10 ml S-basal). This process took nine hours at 37°C. The L1s were pipetted to the 96-well plate the next day. Each drug well had a final volume of 60 µl, which included 20–25 animals in the L1 stage in S-medium, *E. coli* OP50 inactivated resuspended in S-medium complete (30 µl 1M MgSO_4_, 30 µl CaCl_2_, 100 µl 1M Potassium citrate (pH=6.0), 8 µL 5 mg/ml cholesterol, 50 µL Penicillin/Streptomycin, 50 µL Nystatin), and the drug tested. For 4 days, a 96-well plate was continuously gently agitated at 20 °C and 180 rpm.

T2 compound was diluted between 5nM and 0.001 nM in 100% DMSO (Sigma, St. Louis, USA). Then, respective concentrations were subsequently diluted in MilliQ purified water to reach a final concentration ranging between 50 and 0.001uM in 1% DMSO in well.

Locomotion assay

The locomotion assay for G9a/GSK-3β inhibitors was performed to establish a dose-response profile, assessing the effects of pharmacological treatment on the motor deficits exhibited by the transgenic CL2006 strains. The worms were cultured with continuous shaking at 180 rpm and 20°C for four days. Each well contained a final volume of 60 μL, which included 25-30 L1 stage larvae diluted in S-medium, G9a/GSK-3β inhibitors at the designated dose (with the working solution being 2.4 times more concentrated than the final concentration in the well), and inactivated OP50. On the fifth day, locomotion assays were conducted on 30 mm NGM plates fully coated with OP50. Five to ten adult nematodes were positioned in the center of a 1 cm diameter circle on the seeded plates. After one minute, the number of worms remaining within the circle was recorded as an indicator of locomotor defect (LD). Motor behavior assays were performed in triplicate (n=3), with at least 100 worms evaluated per compound concentration. The motor index assigns a score where treated animals showing the same motor impairment as CL2006 receive an index of 0%. Conversely, animals demonstrating improved motor behavior comparable to wild-type (N2) receive an index of 100% (REF). The results presented in the figures were calculated using the following formula: Motor index (%) = ((LD CL2006 vehicle - LD CL2006 drug))/((LD CL2006 vehicle - LD WT vehicle)) x 100.

Thioflavin-S staining Aβ aggregation

After five days of treatment, adult CL2006 *C. elegans* were fixed in 4% Paraformaldehyde/Phosphate-buffered saline (PBS) (pH 7.5) for 24 hours at 4°C. Then, worms were permeabilized in 5% fresh β-mercaptoethanol, 1% Triton X-100, and 125 mm Tris (pH 7.5) at 37°C for another 24 hours. On the last day, nematodes were stained with 0.125% Thioflavin-S (ThS) (Sigma, CAS# 1326–12-1) in 50% ethanol (EtOH) for 2 min, destained in 50% EtOH for 2 min, washed three times with PBS. To prepare the glass slide for microscopy, approximately 10 μL volume was transferred on a droplet of Fluoromount G (Electron Microscopy Sciences, CAT#17984-25). Fluorescence images were acquired using a 20 Å~ fluorescence microscope objective. Aβ in the head region of worms were quantified by counting the number of ThS favorable spots using ImageJ and were expressed as Aβ deposits/anterior area. Aβ aggregates were scored by an investigator blinded to G9a/GSK-3β inhibitor treatments.

***In vivo* efficacy studies in SAMP8**

Animals and treatment

For the in vivo study, 24-week-old female and male SAMP8, 20-week-old mice were utilized to perform cognitive and molecular studies. In this study, the SAMP8 mice were divided into two groups: The SAMP8 control group and the SAMP8 T2 group (5 mg/kg) were utilized in the study. The experimental groups were administered either a daily dose of vehicle (2% w/v, (2-hydroxypropyl)-β-cyclodextrin) or a dose of 5 mg/kg/day of T2 dissolved in water via oral gavage for a period of 15 days. The animals had unrestricted access to food and water and were maintained under standard temperature conditions (22 ± 2°C) and 12-hour light-dark cycles (300 lux/0 lux). Subsequent to the treatment period, cognitive assessments were conducted on the animals.

All procedures involving animals, including behavioral testing and dissection and removal of brains, followed ARRIVE and the standard ethical guidelines (Council of the European Communities Directive 2010/63/EU and Guidelines for the Care and Use of Mammals in Neuroscience and Behavioral Research, National Research Council 2003) and were approved by the Institutional Animal Care and Generalitat de Catalunya (#PJ_159/24, March 13, 2024). Every effort was made to minimize the number of mice used and their suffering.

Cognitive tests

Novel Object Recognition Test (NORT)

Short- and long-term recognition memory were evaluated using the Novel Object Recognition Test (NORT). The test was conducted in a black polyvinyl chloride 90º black maze, which consists of two arms measuring 25 cm in length, 20 cm in height, and 5 cm in width. The light intensity at the center of the maze was maintained at 30 lux. During the first three days, mice were individually habituated to the maze for 10 minutes each day. On the fourth day, two identical novel objects (A + A’ or B + B’) were placed at the ends of each arm, and the animals were allowed to freely explore the apparatus for 10 minutes (first trial or familiarization phase). After this session, mice were removed and returned to their home cages. The retention phase (second trial) was performed 2h and 24h after first trial, to study the short- and long-term memory, respectively. For the short-term memory test, one of the familiar objects was replaced with a different one (A + B’ or B + A’), and mice were again allowed to explore the maze for 10 minutes. For long-term memory assessment, 24 hours later, both objects were replaced with entirely new ones (A + C or B + C), differing in shape and color, and exploration was measured for another 10 minutes. The time (in seconds) the animals spent exploring the new object (TN) and the familiar one (TO) was recorded, and a Discrimination Index (DI) was calculated using the formula: (TN - TO) / (TN + TO). The maze, the surface, and the objects were cleaned with 70% ethanol between the animals’ trials to eliminate olfactory cues.

Open Field

The Open Field Test (OFT) was conducted to assess anxiety-like behaviors. Mice were placed in the center of a white polywood arena (50 × 50 × 25 cm) and allowed to explore freely for a duration of 5 minutes. Behavior was subsequently analyzed using SMART® version 3.0 software, and each trial was recorded for later evaluation. The following behavioral parameters were quantified: the duration spent in the center of the arena, rearing frequency, grooming behavior, and the total distance traveled.

Object Location Test (OLT)

Spatial memory was evaluated using the Object Location Test (OLT) over a period of three days, conducted in a wooden box measuring 50 × 50 × 25 cm, in which three walls were white and one wall black to provide a visual cue. On the first day, mice were placed in the empty box for a 10-minute habituation session to acclimate to the open-field environment. On the second day, two identical objects (10 cm in height) were positioned equidistantly in front of the black wall. Animals were then placed in the arena and allowed to explore both objects and the surrounding area for 10 minutes. On the third day, one of the objects was relocated to a position in front of a white wall to assess spatial memory. Mice were again given 10 minutes to explore the arena. The time (in seconds) spent exploring the object in the novel location (TN) and the object in the familiar location (TO) was recorded. A Discrimination Index (DI) was calculated using the formula: (TN − TO) / (TN + TO), to quantify spatial memory performance. To eliminate olfactory cues that could influence behavior, the apparatus, objects, and surfaces were cleaned with 70% ethanol between trials.

Biochemical experiments

Sample collection

SAMP8 mice were euthanized two days after the completion of the behavioral test by cervical dislocation. Brains were immediately removed from the skull. Cortex and hippocampus were then isolated and frozen on powdered dry ice. They were maintained at -80 °C for biochemical experiments. For the Thioflavin, Immunohistochemistry and Golgi staining protocol, see the procedure in the sections “Aβ plaques histology and Immunofluorescence assay” and “Spine density and Golgi staining”, respectively.

Approximately 0.1 ml of blood was extracted via intracardiac injection and collected in EDTA-K2 tubes. The tubes were then placed on wet ice until the process of centrifugation proceeded. Blood samples were centrifugally treated at 3200g for 10 minutes to obtain plasma, subsequently transferred to polypropylene tubes, and flash frozen on dry ice.

**ELISA determinations**

Cortical tissue and plasma samples were collected and used for several Elisa determinations. Quantification of amyloid-β40 (Invitrogen, #KMB3481, #KHB3481) and amyloid-β42 (Invitrogen, #KMB3441, # KHB3441), p-Tau (T181) (Fine test, #EM2110), p-Tau (T217) (Fine test, #EM2108), and H3K9me2 (Epigentek, #P-3116-96), were performed following the manufacturer’s instructions.

**Protein Level Determination by Western Blotting (WB)**

Cortical mouse tissue samples, and parietal human samples were homogenized in lysis buffer supplemented with phosphatase and protease inhibitors (Cocktail II, Sigma-Aldrich). Protein concentrations were determined using the Bradford assay. Equal amounts of protein were subjected to sodium dodecyl sulfate–polyacrylamide gel electrophoresis (SDS-PAGE; 10%–14%) and transferred onto polyvinylidene difluoride (PVDF) membranes (Millipore). Membranes were blocked with 5% bovine serum albumin (BSA; Sigma) in Tris-buffered saline containing 0.1% Tween 20 (TBS-T) for 1 hour at room temperature. Following blocking, membranes were incubated overnight at 4°C with primary antibodies (detailed in Table S10), diluted in 5% BSA in TBS-T. The following day, membranes were washed and incubated for 1 hour at room temperature with the corresponding secondary antibodies (also listed in Table S10). Protein detection was carried out using an enhanced chemiluminescence (ECL) kit (Millipore), and digital imaging was performed with the Imager 680 system (Amersham Bioscience). Band intensities were quantified using ImageLab software (BioRad) and expressed in Arbitrary Units (AU), considering the control mice group as 100%. Values were normalized to glyceraldehyde 3‐phosphate dehydrogenase (GAPDH).

**RNA extraction and gene expression determination**

Total RNA was extracted from hippocampal tissue and *C. elegans* samples using TRIsure™ reagent (Bioline Reagents, #BIO-38032), following the manufacturer’s protocol. RNA concentration, purity, and integrity were assessed spectrophotometrically using a NanoDrop™ ND-1000 and an Agilent 2100 Bioanalyzer. Complementary DNA (cDNA) was synthesized from total RNA using the High-Capacity cDNA Reverse Transcription Kit (Applied Biosystems, Foster City, CA, USA). Quantification of mRNA expression for selected target genes (listed in Table S11) was performed using SYBR® Green-based real-time PCR on a StepOnePlus™ Real-Time PCR System (Applied Biosystems). Relative gene expression was calculated using the comparative threshold cycle method (ΔΔCt), where the housekeeping gene level (β-actin) was used to normalize differences in sample loading and preparation.

**MDA Levels**

The levels of malondialdehyde (MDA) in the tissues were determined using the Lipid peroxidation assay colorimetric following the manufacturer’s instructions (Canvax, Córdoba, Spain). The absorbance was measured at 532 nm using a NanoQuant microplate reader (Tecan Trading AG, Männedorf, Switzerland).

**ROS Levels**

Reactive oxygen species (ROS) in cells were estimated as following; tissues was homogenized in ice-cold Tris-HCl buffer (40 mM, pH 7.4). Samples were mixed with 2′, 7′-dichlorofluorescein diacetate (1 μM) prepared in Tris-HCl buffer (40 mM, pH 7.4). The mixture was incubated for 30 min at 37 °C. Finally, the fluorescence intensity of the samples was measured using a NanoQuant microplate reader (Tecan Trading AG, Männedorf, Switzerland) (λ excitation 485 nm and λ emission 525 nm).

**Detection GSH/GSSG Levels**

Oxidized glutathione (GSSG) was measured as follows; briefly, after Pyroglutamic acid extraction (Sigma-Aldrich, Madrid, Spain), reduced glutathione (GSH) was derivatized with 2-vinylpyridine at room temperature for 1 h, and the reaction was monitored as follows. Reduced GSH was obtained by subtracting GSSG levels from total glutathione levels. To determine the ratio GSH:GSSG in the tissues in a luminescent-based assay kit (Promega, WI, USA) was used.

**Determination Iron (II) Levels**

Iron Assay Kit (ThermoFisher Scientific) was used to determine the amount of iron (II) present in cell samples and sEV as a marker for ferroptosis. Absorbance at 592 nm was measured using a NanoQuant microplate reader (Tecan Trading AG, Männedorf, Switzerland).

**Aβ plaques histology**

Mice were euthanized by cervical dislocation and brains were carefully removed from the skull and fixed in 4% paraformaldehyde (PFA) at 4 °C overnight. The next day, tissues were transferred to a cryoprotective solution consisting of 4% PFA supplemented with 15% sucrose. Finally, brains were frozen in isopentane and stored at − 80 °C. Frozen brains were embedded in OCT Cryostat Embedding Compound (Tissue-Tek, Torrance, CA, USA) and sectioned at a thickness of 30 μm using a cryostat (Leica Microsystems, Germany) maintained at −20 °C.

For Thioflavin-S (Th-S) staining, three coronal brain sections per animal were rehydrated in 1× phosphate-buffered saline (PBS) for 5 minutes at room temperature. Sections were then sequentially washed with 50%, 70%, and 80% ethanol solutions. Immediately after, brain slices were incubated in a 0.3% Th-S solution (Sigma-Aldrich) for 10 minutes in the dark at room temperature. Post-staining, sections underwent successive 1-minute washes in 80%, 70%, and 50% ethanol, followed by a final wash in 1× PBS for 5 minutes. Slides were then mounted using Fluoromount-G™ (Electron Microscopy Sciences, Hatfield, NJ, USA) and left to dry overnight in the dark. Fluorescence imaging was performed using a laser fluorescence microscope (Leica Thunder, Germany) with a 10× objective. For quantification of Th-S-positive plaques, ImageJ software was used. Images were captured from anatomically comparable regions, specifically focusing on the cortex and hippocampus. Image analysis and quantification were performed using ImageJ software. The final plaque count for each animal was obtained by averaging data from the three analyzed sections.

**Spine density and Golgi staining**

Mice were euthanized by cervical dislocation, and the brains were removed from the skull. The FD Rapid Golgi Stain kit (FD NeuroTechnologies incs, #PK401) was used for Golgi staining according to the manufacturer's instructions. Neuronal images for dendritic branching analysis were captured at 20x magnification using a Leica Thunder microscope (Germany). Dendritic neurite length and complexity were quantified using NeuronJ macros and Advanced Sholl Analysis software. The number of intersections (branch points) within concentric circles of 10 µm radius was calculated and compared between groups. For spine density analysis, neuronal images were acquired at 50x magnification using an oil-immersion objective. Only neurites with a length of approximately 18 µm and within a 150 µm radius from the soma were analyzed.

**Transcriptomic analysis**

Bulk RNA-seq analysis

Paired-end RNA-seq reads were QC’d using fastQC v. 0.11.9. All of them had good quality and were used in the analysis. Reads were aligned against the mouse transcriptome (Ensembl annotation v99, based on mm10) with HISAT 2.1.0 using the following non-standard parameters: --no-unal --no-mixed --no-discordant. Alignments with QS <10 or falling within Encode blacklisted regions were eliminated. Read count tables were obtained with featureCounts from the Rsubred package (2.10.5, R version 4.2.1) using the following parameters: isGTFAnnotationFile=T, useMetaFeatures=T, minMQS=10, largestOverlap=T, isPairedEnd=T, requireBothEndsMapped=T, with the Ensembl annotation v99 as reference. The read count table was analyzed with edgeR v3.38.4. Lowly expressed genes (cpm<3) were eliminated. After calculating normalization factors and estimating dispersion, differentially expressed genes between the four groups were calculated using glm modeling. Only genes with an absolute log2 fold change > 0.5 and a FDR <1e-2 were considered as DEG. Heatmaps of DEG were generated using heatmap.2 from the gplots package v3.1.3. Gene ontology analyses were performed using DAVID using the list of all expressed genes as background. RNA age was calculated using the RNAAgeCalc package from Bioconductor. Mouse gene symbols were converted to human using the mouse2human_eset function from the IOBR package. Converted IDs were used in the predict_age function with tissue set to “brain” and type set to “all”.

ChIP-seq and ATAC-seq analysis

Paired-end single-cell RNA-seq reads were QC’d using fastQC v. 0.11.9. Reads were trimmed using fastp 0.26.0. Reads were aligned to mouse genome (mm39) using bowtie2 v2.5.4. Significant peaks were calculated with macs3 v3.0.2, using a --broad-cutoff of 1e-2 for H3K9me2 and a regular narrow peak calling with an FDR of 1e-5 for ATAC-seq. Enrichment plots were created using deeptools v3.5.5. Enrichment within peaks was calculated with computeMatrix from deeptools and fold enrichments were calculated using R v4.5.0. Genes associated with peaks were obtained with the closestBed program from the beedtools suite v2.31.1. Gene annotations were obtained with our get gencode annotation tool (<https://github.com/david-valle/gencode_annotation>). Lists comparisons between ChIP-seq and single-cell-RNA-seq analyses were made with FilterLine.pl from our useful scripts suite (<https://github.com/david-valle/useful_scripts>).

Single-Cell RNA-seq analysis

Paired-end single-cell RNA-seq reads were QC’d using fastQC v. 0.11.9. All of them had good quality. The analysis was performed using the CeleScope pipeline v2.0.7. Briefly, UMIs were extracted, filtered (quality score >30) and corrected (UMIs with a hamming distance >1). Reads were mapped to the mouse transcriptome (Ensembl annotation v99, based on mm10) using STAR. Analysis was performed with Seurat 5.3.0 in an R 4.5.0 environment. Cells with low UMI count and, or high mitochondrial counts were filtered out. PCA, Clustering, UMAP and gene marker finding was performed using default values unless noted otherwise. 15 dimensions from PCA with a resolution of 0.5 were used for clustering, which generated 11 clusters for SAMP8 control and 13 for T2. Cell type annotation was performed with singleR using the mouse_rnaseq (2024-02-2) database from celldex. Gene Ontology, Pathway and Disease analyses were performed using Enrichr from the genes that were unique markers in T2 compared to SAMP8. Only categories with FDR values <0.05 were considered significant.

Proteomic analysis

Brain tissue samples were sent to HaploX Biotechnology Co., Ltd. (Shenzhen, China) for proteomic analysis by data independent acquisition (DIA) mass spectrometry.  After peptide extraction and trypsin digestion, a peptide library was constructed and subjected to in-house quality control prior to LC-MS/MS analysis on an Orbitrap Astral high-resolution mass spectrometer. The system integrates quadrupole, Orbitrap and Astral analysers to enable high-throughput, high-resolution acquisition. Internal retention time standards (iRT kit) were used to normalise retention times between samples.

Protein identification and quantification was performed with DIA-NN software<https://github.com/vdemichev/DiaNN>, searching the SwissProt Mus musculus UniProt database (version 2024_07_26, 17,224 entries; <https://www.uniprot.org/>). A total of 7,850 proteins were identified and quantified. Proteins showing a fold change >1.2 or <0.83 with a p-value <0.05 were considered differentially expressed.

To explore group-wise variation in the proteomic profiles, Principal Coordinates Analysis (PCoA) based on Bray–Curtis dissimilarity was applied to the quantified protein data, enabling visualization of sample dispersion in multidimensional space. Functional interpretation of differentially expressed proteins was carried out through Gene Ontology (GO) and Kyoto Encyclopedia of Genes and Genomes (KEGG) enrichment analyses. Additionally, protein–protein interaction (PPI) networks were generated using the STRING database (v12.0) <https://string-db.org/>. Confidence thresholds for interaction scores were adapted to the size of each protein set to ensure robust and interpretable networks, which were used to identify functionally enriched modules and potential regulatory hubs.

**Human cases**

Tissue samples were obtained from the Institute of Neuropathology-IDIBELL Brain Bank, Hospitalet de Llobregat, following the guidelines of Spanish legislation on this matter and the approval of the local ethics committee (Table S12). The postmortem interval between death and tissue processing was 1 to 10 h and samples were processed to minimize postmortem delay artifacts. The brain tissue was immediately frozen on metal plates over dry ice, placed in individual air-tight plastic bags, and maintained at -80 °C for biochemical experiments. The average age in the ND group is 69.16 ± 15.06, and in the AD group is 81.13 ± 7.03. The neuropathologic diagnosis of AD was based on the classification of Braak.

CSF samples were obtained through lumbar puncture, and were centrifuged (2000rpm x 10 mins), aliquoted and stored at -80ºC until analysis. Concentrations of Aβ1–42, Aβ1–40, tTau and pTau181 in CSF were measured using commercially available kits in the Lumipulse fully-automated platform (Fujirebio-Europe), as previously described (PMID: 31464088). Plasma samples were collected in EDTA-K2 tubes and subsequently centrifuged (2000rpm x 10 mins) within 2 hours after extraction. Plasma was aliquoted and stored at -80ºC until analysis. Plasma and CSF samples were collected simultaneously. Full protocol for CSF and blood sample collection in our center has been previously reported in detail (PMID: 31650016).

Additional determinations of amyloid-β40 (Invitrogen, #KHB3481) and amyloid-β42 (Invitrogen, #KHB3441), p-Tau (T181) (RayBio, #PEL-Tau-T181-Q ),TNF-α (Millipore, #EZHTNFA-150K), GFAP (Invitrogen, #EEL098), NF-L (UmanDiagnostic, #20-8002 RUO), H3K9me2 (Epigentek, #P-3116-96, SMOC1 (MyBiosource, #MBS3804789-96) were performed following the manufacturer’s instructions.

**Statistical Analysis**

The statistical analysis was conducted using GraphPad Prism version 9.2 statistical software. Group size may differ depending on power analysis and expertise of the authors. Shapiro–Wilk test to verify data normality for all groups. Data were expressed as the mean ± Standard Error of the Mean (SEM). For normally distributed data, means were compared in One-Way or Two-Way ANOVA analysis of variance, (ANOVA), followed by Tukey’s post-hoc analysis. Moreover, unpaired t-tests were used in several instances to compare the performance of the groups. Non-linear regressions were used to calculate the IC50. In contrast, the Mann-Whitney or Kruskal–Wallis test followed by Dunn’s post-hoc analysis was used for non-normally distributed data. Statistical significance was considered when *p*-values were less than 0.05. For behavioral tests, a blinded analysis was performed.
